# Temperature-sensitive cortical condensation modulates mechano-response at Tricellular Junctions

**DOI:** 10.1101/2024.12.11.628049

**Authors:** Qi-Hong Zheng, Chengge Zhang, Ming-Xin Wang, Xiaoxiang Xiang, Shaohong Zhang, Yaru Wang, Huapeng H. Yu

**Author notes:** These authors contributed equally (Qi-Hong Zheng, Chengge Zhang, Ming-Xin Wang). Correspondence (H.H.Y).

## Abstract

A general mechanism to maintain epithelial integrity during cell movements is to couple adhesion strength with mechanical force. Mechanosensitive proteins are recruited to adhesion sites to reinforce cell-cell or cell-matrix linkage in response to elevated tension. However, subcellular mechanisms for highly dynamic protein transport into and out of tension-bearing structures remain elusive. Furthermore, it is still unknown whether mechanosensitivity can be modulated by physiological signals *in vivo*. Here we revealed a direct “edge-to-vertex” route of Canoe/Afadin transport to tricellular junctions under tension, which relies on an obligatory binding partner Mbt/PAK. Without Mbt/PAK, Canoe/Afadin becomes insensitive to cytoskeletal tension and forms “tactoid-shaped” condensates directly at bicellular junctions. Remarkably, a temporary increase in temperature within the physiological range dissolves these condensates, allowing Canoe/Afadin to regain mechanosensitivity and enrichment at tricellular junctions. Functionally, the localization of Mbt/PAK at cell junctions oscillates during synchronous cell division within embryonic mitotic domains, enabling dynamic adjustments of Canoe/Afadin mechanosensitivity to preserve epithelial integrity in rapidly dividing tissues. Collectively, our findings suggest a physiological process in which the state of protein condensation, regulated by biochemical inputs, can supersede mechanical signals to modulate adhesion strength during cell division.

## Introduction

A defining characteristic of multicellular organisms is the physical assembly of hundreds to billions of single cells into a unified and coordinated entity. This integration relies on a fundamental cellular apparatus named cell-cell junctions, which serves to mechanically link neighboring cells together^1–3^. Furthermore, while both unicellular and multicellular organisms are capable of rapid and repeated cell divisions, the establishment of physical connections among daughter cells post-division represents a pivotal moment in the evolution of multicellularity to ensure tissue continuity during proliferation^4–6^. Previous studies have extensively investigated how individual cells divide and remodel their adhesion with neighboring non-dividing cells^7–10^. However, in the early stage of animal development, cell divisions are often synchronized, with all or a subset of adjacent cells dividing simultaneously^11–13^, a scenario that presents a unique challenge in terms of re-establishing adhesion post-division without the physical support from neighboring non-dividing cells^11,14^. How synchronously dividing cells remodel and restore adhesion post-division are poorly understood.

Cell-cell junctions serve as crucial sites where mechanical and biochemical signals converge to regulate cell adhesion in cultured cells and animal models^3,15^. These junctions exist in two modular forms: bicellular junctions, where two cells establish an edge contact, and tricellular junctions, where three cells come together to form a vertex^16,17^. Analogous to focal adhesion at cell-matrix contacts^18^, cell-cell adhesion typically reinforces in response to increased tension by recruiting mechanosensitive actin-binding molecules like Vinculin^19,20^, Ajuba^21,22^, ZO-1^23^ and Canoe/Afadin^24,25^, to strengthen the membrane-cytoskeleton linkage. The “knee-jerk” reaction-alike mechanism serves to maintain tissue integrity and barrier functions during force-induced events such as cell rearrangements and collective migration^26,27^. Similarly, cell division is a multi-step force-driven process mediated by cytoskeleton re-organization and actomyosin contraction, characterized by extensive cell shape changes such as mitotic rounding at metaphase and membrane cleavage at telophase^5,28,29^. In a multicellular context, each cell division results in the *de novo* formation of a bicellular junctions between the two daughter cells and two tricellular junctions with neighboring cells^30^. However, it remains largely unknown whether cell adhesion is similarly coupled to cytoskeletal forces during mitosis at pre-existing or newly formed cell junctions, and if not, what mechanisms underlies this decoupling.

A universal property of cells is the ability to sense and respond to both biochemical and mechanical inputs. Biochemical signals can trigger changes in protein conformations through post-translational modifications or a “key-lock” allosteric mechanism. Mechanical forces, on the other hand, can physically stretch proteins to expose hidden active site^31–33^ or induce a switch into an alternate conformation^34^. Given that both biochemical and mechanical inputs influence biological processes by altering protein conformations, a crucial question arises regarding how two types of signals are coordinated at the molecular level. Previous studies have mainly found evidence of synergistic effects, demonstrating that mechanical stretching can activate kinase activities or phosphorylation events can enhance mechanosensitivity^25,35,36^. However, the molecular basis and physiological context of how cells respond when confronted with conflicting inputs from biochemical and mechanical signals remain largely unexplored.

Early gastrulation of *Drosophila* embryos involves rapid collective cell movements and patterned synchronous mitosis^12,13,37^, offering a valuable model to investigate the coordination of mechanical forces and cell adhesion in a physiological setting. Here, we identified a novel tricellular junction protein named Mbt/PAK, which is utilized by biochemical signals to desensitize Canoe/Afadin from mechanical inputs during synchronous cell division. Oscillation of junctional Mbt/PAK during synchronous cell division enables dynamic modulation of tricellular adhesion independent from actomyosin activities and prevents epithelial rupture in hyperproliferative tissues. Mechanistically, Mbt/PAK facilitates the direct “edge-to-vertex” transport of Canoe/Afadin from bicellular to tricellular junctions in a kinase-independent manner. In the absence of this mechanism, endogenous Canoe/Afadin forms tactoid-shaped condensates at bicellular junctions *in vivo*, rendering it inert to changes in cytoskeletal forces. Strikingly, the condensation of endogenous Canoe/Afadin exhibits temperature-sensitive behavior, undergoing phase transitions within physiologically relevant temperature ranges *in vivo*. Together, our findings suggest that condensational states of proteins can be modulated by physiological cues to regulate mechanosensitivity of cell junctions.

## Results

### Canoe/Afadin tricellular localization is uncoupled from tension during synchronous cell division

To investigate how cell division affects adhesion at tricellular junctions, we first analyzed single-cell mitotic events (Extended Data Fig. 1a) surrounded by non-dividing neighboring cells at stage 9 (after egg laying, AEL 6 hrs) *Drosophila* embryos. We found that transmembrane adhesion receptor Sidekick^38–40^ and mechano-sensitive adaptor Canoe/Afadin^24,25^ remain enriched at tricellular junctions throughout mitosis (Extended Data Fig. 1b, c). This result suggests that mitotic cells maintain close intercellular contacts with neighboring cells and cytoskeletal tension at cell junctions is not significantly altered during cell cycle. However, due to microscopic resolution limits, one cannot assign the fluorescent signal at tricellular junctions to the mitotic cell or the other two non-dividing neighboring cells (Extended Data Fig. 1a, question mark). Therefore, it was not possible to conclude whether mitotic signals affected Sidekick or Canoe/Afadin tricellular localization in a cell-autonomous manner. To circumvent the resolution limit, we switched to examine mitotic domains in stage 7-8 embryos (AEL 3-4 hrs) where three cells connected through the same tricellular junction can divide all at once (Fig. 1a, Supplementary Video 1). The position and timing of division at these mitotic domains are strictly defined by morphogen patterns and consistent between embryos^12,13^. Combining high-speed fluorescent imaging and genetically encoded fluorescent markers, we can track the dynamics of cytoskeletal tension and adhesion at bicellular and tricellular junctions, before, during and after synchronous cell division. With mitotic phases distinguished by stereotypical spindle shapes visualized via mCherry-Tubulin, we found that tricellular contacts and Sidekick localization are well-maintained at metaphase during synchronous division (Fig. 1b), a pattern similar to single-cell mitosis. In contrast, despite of intact tricellular junctions, Canoe/Afadin gradually lost tricellular enrichment as the cell cycle progressed from interphase to metaphase (Fig. 1c, d, g, Supplementary Video 1). Canoe/Afadin tricellular enrichment has been shown to be dependent upon actomyosin-mediated cytoskeletal tension^25^. Therefore, we hypothesized that cortical actomyosin activity might be similarly downregulated. However, fluorescent intensities of Myosin light chain at the cell cortex remained largely unchanged from interphase to metaphase (Fig. 1e, f, j), consistent with cytoskeletal and membrane tension measurements during single-cell mitotic events by previous studies^7,8,41–43^. These results implicate a non-mechanical signal that might contribute to the delocalization of Canoe/Afadin from tricellular junctions during mitosis.

**Fig. 1:**
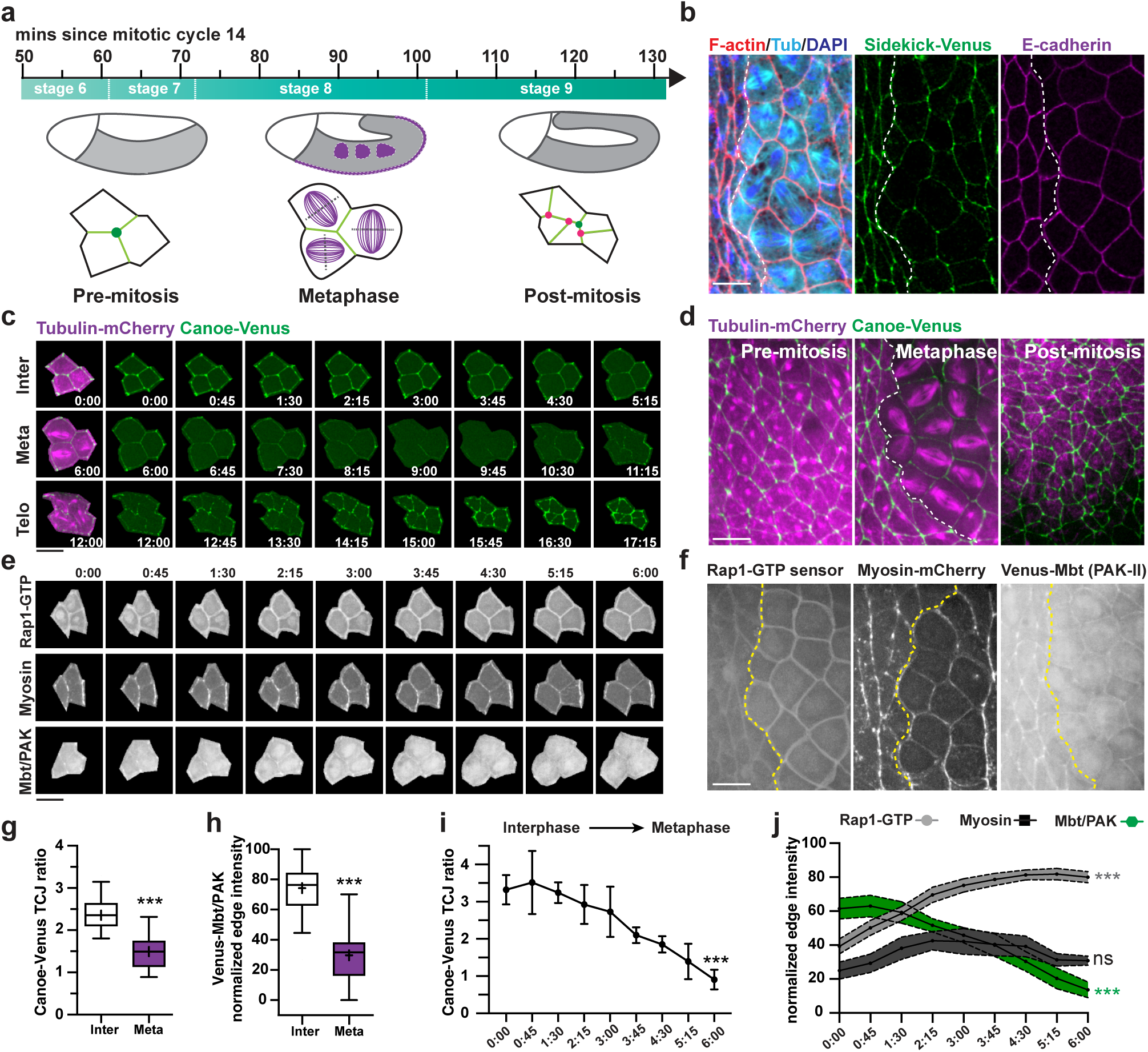
Canoe/Afadin is uncoupled from mechano-regulation during synchronous cell division. **(a)** Time scales of *Drosophila* embryonic stages and the location of mitotic domains (Purple) where cells divide in a synchronous manner (top); schematics of synchronous cell division of three cells connected to the same tricellular junction (bottom). Note the de novo formation of three additional tricellular junctions after one round of division (red dot). **(b)** Localization of tricellular and bicellular junction integrity marker Sidekick and E-cadherin in synchronously dividing cells at metaphase, characterized through rounded cell shapes (F-actin) and aligned mitotic spindles (Tub, tubulin). A dotted line separates cells at metaphase within the mitotic domain (right) from non-dividing neighboring cells (left). **(c, d)** Still images of a dual-color time lapse movie recording Tubulin-mCherry and endogenously Venus-tagged Canoe/Afadin every 45 seconds (c), representative images of cells before, during and after synchronous cell division were shown in (d). **(e)** Still images of a dual-color time lapse movies recording Venus-Ralgds-RBD (Rap1-GTP sensor) and Myosin-mCherry (top and middle), and a separate movie recording Venus-Mbt/PAK alone (bottom). **(f)** Localization of Rap1-GTP sensor, mCherry-Myosin and Venus-Mbt/PAK in synchronous dividing cells (right side of the dotted line) and neighboring non-dividing cells (left). **(g)** Quantification of Canoe/Afadin enrichment at tricellular junctions during interphase and metaphase using TCJ ratio as the metric. **(h)** Quantification of cortical localization of Mbt/PAK during interphase and metaphase using normalized edge intensity as the metric. **(i, j)** Quantification of changes in Canoe/Afadin TCJ enrichment, junctional intensity of Rap1-GTP sensor, myosin, and Mbt/PAK over time as cells progress from interphase (time point 0:00) to metaphase (time point 6:00). All Venus-tagged Canoe/Afadin at the endogenous locus simplified as “Canoe-Venus” hereafter. Ectopically expressed Venus-tagged Canoe transgene will be referred to as “Venus-Canoe”. Boxes are 2^nd^ and 3^rd^ quartiles and whiskers are 5^th^ to 95^th^ percentiles; horizontal line is the median and “+” sign is the mean value. Mean±SD in (i) and Mean±SEM in (j). ***P<0.001; Not significant (ns) P>0.1. Welch’s t-test for all comparisons. See table S4 for detailed quantification values. Fixed embryos in (b) and live embryos in all other panels. Embryos are stage 7 (before), 8 (during) and 9 (after synchronous division), positioned with anterior left, ventral down. Scale bars, 10 μm.

### Mbt/PAK is required for Canoe/Afadin mechanosensitivity

To search for non-mechanical upstream signals that might delocalize Canoe/Afadin during mitosis, we next examined a well-established biochemical regulatory input of Canoe/Afadin named Rap1. As a Ras-family small GTPase, Rap1 has been shown to be critical for Canoe/Afadin tricellular localization (Extended Data Fig. 2a). Downregulation of Rap1 or a Rap1-Guanine Exchange Factor (GEF) named Dizzy decreased Canoe/Afadin junctional localization and tricellular enrichment in cells at interphase^44,45^. Therefore, one possibility is that Rap1 activity is temporally down-regulated as cells progress into metaphase. To probe for Rap1 activity, we newly developed a Rap1-GTP sensor by fluorescently-tagging the Rap1-binding domain of a well-conserved downstream effector named RALGDS^46^ (Extended Data Fig. 2b). When expressed in pre-mitotic stage-7 embryos, the Rap1-GTP sensor localized evenly to the cell cortex with no apparent enrichment at tricellular junctions (Extended Data Fig. 2c). Notably, knockdown (KD) of Rap1 or Rap1 activator Dizzy through RNA-interference (RNAi) completely delocalized the sensor into cytoplasm, suggesting that the cortical localization of RALGDS-RBD is dependent on the Rap1 activity (Extended Data Fig. 2c, e). However, the dependency might be indirect since it is possible that the delocalization results from cell-cell adhesion defects caused by Rap1- or Dizzy-RNAi, exemplified by apparent gaps between myosin cables (arrowheads, Extended Data Fig. 2d). In other words, all junction or cortical localized protein might be similarly affected as the Rap1-GTP sensor by Rap1- or Dizzy-RNAi. To examine this possibility, we expressed the Rap1-GTP sensor in Abl- and Canoe/Afadin-RNAi embryos that exhibited similar adhesion defects (Extended Data Fig. 2d). However, the Rap1-GTP sensor localized normally at the cell cortex in these embryos (Extended Data Fig. 2c, e), suggesting that its cortical localization does not require intact cell-cell adhesion. Therefore, the localization of Rap1-GTP sensor likely depends on Rap1 activation and therefore detects endogenous Rap1 activity in living embryos. Next, we tracked Rap1 activity over time with this sensor during synchronous cell division in stage-8 embryos. To our surprise, Rap1 activity increased when cells enter metaphase (Fig. 1e, f, j, Supplementary Video 2), which would not account for the decreased tricellular localization of Canoe/Afadin (Fig. 1i), as a positive regulator (Extended Data Fig. 2a). Therefore, there might exist a Rap1-independent signal to uncouple Canoe/Afadin from mechanical inputs during cell cycle.

To search for potential regulators of Canoe/Afadin tricellular localization, we took two independent approaches. First, we systematically characterized proteins enriched at tricellular junctions using a TurboID-based proximity proteomic assay. Second, we performed RNAi-screens against known Canoe/Afadin interactors using tricellular enrichment ratio (TCJ ratio) of endogenously Venus-tagged Canoe/Afadin as the quantitative metric. Both approaches led us to focus on a highly conserved serine/threonine kinase named *Mushroom body tiny* (Mbt/PAK), the only member of group-II p21-activated kinase family (PAK) in *Drosophila*. Mbt/PAK has been previously shown to localize at bicellular adherens junctions by immunofluorescence in adult *Drosophila* eye and mammalian cultured cells^47–49^. Here, we observed a slight but consistent enrichment of Venus-tagged Mbt/PAK at tricellular junctions in *Drosophila* embryonic epithelia (Fig.1e, f, Extended Data Fig. 3a, b), a feature only pronounced when examined in live embryos but lost after formaldehyde fixation (Extended Data Fig. 3a, b). Next, we compared Mbt/PAK junction localization between cells at interphase and metaphase in live embryos during synchronous cell division (stage-8). We found that the localization of Mbt/PAK is significantly reduced at junctions between synchronously dividing cells (Fig.1e, h, Extended Data Fig. 3c). Time-lapse tracking revealed that Mbt/PAK junctional intensity gradually decreased as cells progressed from interphase to metaphase (Fig.1e), coinciding with the loss of Canoe/Afadin tricellular enrichment (Fig.1i, j). We further validated the temporal correlation using dual-color imaging to visualize dynamics of mCherry-tagged Mbt/PAK and Venus-tagged Canoe/Afadin at the same time (Supplementary Video 3). Together, these results implicate Mbt/PAK as a potential regulator of Canoe/Afadin tricellular localization.

To determine whether Mbt/PAK is required for Canoe/Afadin tricellular enrichment, we quantified Canoe/Afadin TCJ ratio in control and Mbt/PAK-RNAi embryos. In addition, we newly generated a *Mbt/PAK* mutant *Drosophila* line with an early stop codon using CRISPR (Extended Data Fig. 3d). We found that endogenous Canoe/Afadin almost entirely lost tricellular enrichment in either Mbt/PAK-RNAi or *Mbt/PAK* homozygous mutant embryos (Fig.2a, c, Extended Data Fig. 3e, Supplementary Video 4), similar to Dizzy- or Rap1-RNAi embryos. One possibility is that Mbt/PAK regulates Rap1 activity, which in turn is required for Canoe/Afadin localization. However, Mbt/PAK knockdown has no impact on the localization or intensity of Rap1-GTP sensor at cell junctions (Extended Data Fig. 2c, e). Another potential explanation is that Mbt/PAK downregulation disrupts the integrity of bicellular or tricellular adherens junctions, which is prerequisite for Canoe/Afadin tricellular enrichment^44,50^. However, we found that Mbt/PAK knockdown only caused local epithelial breaches at multicellular vertices, with transmembrane receptors Sidekick and E-cadherin localized mostly normal at tricellular and bicellular adherens junctions (Extended Data Fig. 3f). Therefore, the local adhesion defects observed in Mbt/PAK-RNAi embryos are mostly likely a result rather than a cause of the embryo-wide loss of Canoe/Afadin tricellular localization. In addition, Canoe/Afadin expressed at similar protein levels in control and Mbt/PAK knockdown embryos (Extended Data Fig. 3g). These findings indicate that Mbt/PAK is likely directly required for Canoe/Afadin recruitment to tricellular junctions, without affecting its protein stability or overall cell-cell adhesion integrity.

To further probe whether Mbt/PAK affects Canoe/Afadin mechanosensitivity, we analyzed Canoe/Afadin dynamics and localization under physiological and ectopically-enhanced actomyosin activity in control and Mbt/PAK-RNAi embryos. The following evidence led us to propose that Canoe/Afadin becomes insensitive to actomyosin-generated tension without Mbt/PAK. First, Canoe/Afadin is not enriched at tricellular junctions in Mbt/PAK-RNAi or Mbt/PAK mutant embryos, despite of normal actomyosin localization and cortical intensity (Fig.2b-d, Supplementary Video 4). Second, spatial and temporal correlations between Canoe/Afadin and myosin observed in control embryos becomes statistically insignificant after Mbt/PAK knockdown (Fig.2e-g). Third, we performed high-speed imaging to track movement of endogenous Venus-tagged Canoe/Afadin at single-junction resolution. In control embryos, we noted that Canoe/Afadin can form individual puncta-like structures at bicellular junctions. These puncta can collide and merge, exhibiting a “edge-to-vertex” flow towards tricellular junctions (Fig.2b, right panel top). However, in Mbt/PAK-RNAi embryos, Canoe/Afadin initially forms “rod-shaped” structures at bicellular junctions, which transformed into a “tactoid-shape” with a long tail over time, as if proteins are disproportionally pushed to one side within a confined space (Fig.2b, right panel bottom). In other words, the cortical movement of Canoe/Afadin was confined to the inside of tactoid-shaped structure at bicellular junctions and failed to eventually arrive at tricellular junctions. Lastly, we ectopically enhanced actin cytoskeleton tension by over-expressing a Rho-kinase activator named Shroom-A^51^ in a striped tissue pattern using *paired*-GAL4. Myosin intensity precisely increased within the tissue stripe where ShroomA was locally expressed in a cell-autonomous manner (Fig.2h, k), offering a mosaic system to directly compare mechano-response between cells inside and outside the stripe. As a proof of principle, a canonical mechanosensitive actin-binding protein named Ajuba^21,22,50^ exhibits nearly two-fold increase in junctional intensity within the stripe (Fig.2i, k), demonstrating heightened mechano-response at cell junctions upon striped expression of Shroom-A and increased myosin cortical localization. Similarly, Canoe/Afadin becomes further enriched at tricellular junctions within the stripe in control embryos (Fig.2j, l). In contrast, Canoe/Afadin exhibits no enrichment at tricellular junctions within or outside of stripe in Mbt/PAK-RNAi embryos (Fig.2j, l), suggesting that simply increasing cytoskeletal force is not sufficient to reactivate mechanosensitivity of Canoe/Afadin. Together, these results led us to propose that Mbt/PAK is required for Canoe/Afadin mechanosensitivity to either physiological or enhanced cytoskeletal tension.

### Mbt/PAK promotes adhesion re-establishment after mitosis

To explore physiological relevance of modulating Canoe/Afadin mechanosensitivity by Mbt/PAK during cell division, we characterized embryonic phenotypes of Mbt/PAK-RNAi and *Mbt/PAK* mutant embryos. Although homozygous *Mbt/PAK* mutant embryos is fertile and viable, hatch rate is significantly reduced (from 94.1% to 64.6%) and adult rate is not furtherly reduced, indicative of impaired embryogenesis (Extended Data Fig. 4a). Temporal analysis of embryonic phenotypes across developmental stages revealed earliest defects during ventral midline formation, a well-conserved process that resembles neural tube closure in mammals^52^. At the onset of gastrulation, a group of cells along the entire length of ventral side undergo apical constriction and bend inward to form a tubular structure named ventral furrow^53^ (Extended Data Fig. 3a, Supplementary Video 5). Internalization of ventral furrow brings two parallel rows of cells named mesectoderm into contact (Fig.3a, Extended Data Fig. 4b, orange colored), which then undergoes synchronous cell division and seal the furrow to form the ventral midline^54^ (Fig.3a, Extended Data Fig. 4c, Supplementary Video 5). The sealing process relies on *de novo* junction formation between two newly apposed rows of mesectoderm cells (Extended Data Fig. 4c, d, top panel). Whereas ventral furrow formation, juxtaposition of mesectoderm and subsequent synchronous cell division all progress normally in Mbt/PAK-RNAi or mutant embryos (Fig.3b, Supplementary Video 6), mesectoderm cells failed to establish stable adhesion after division (Supplementary Video 6), particularly at tricellular junctions between apposed rows of cells (Extended Data Fig. 4d, lower panel). Interestingly, the epithelial defects coincide with abnormal accumulation of Canoe/Afadin at bicellular junctions and thus failed tricellular recruitment (Fig.3b, Extended Data Fig. 4d, lower panel). Consequently, the ventral midline failed to be sealed exemplified by exposed underlying mesoderm (Fig.3b, c, Extended Data Fig. 4d, Supplementary Video 6), quantified as markedly increased epithelial gap area (Fig.3d). In contrast, the epithelial adhesion remained largely intact among the non-dividing germ band cells, with minor defects only observed at multicellular vertices (Fig.3e, f). These results led us to hypothesize that Mbt/PAK might be particularly required for maintain adhesion in tissues with higher mitotic activities.

To investigate potential contributions of mitotic activity to *Mbt/PAK* mutant phenotypes, we examined epithelial integrity under hypo-and hyper-proliferative conditions by manipulating a well-established regulator of embryonic mitosis named String/Cdc25^55^ (Fig.3a, lower right). Consistent with previous reports^54^, down-regulation of String/Cdc25 completely inhibited synchronous cell division of ventral midline cells in both control and *Mbt/PAK* mutant embryos (Fig.3c, Supplementary Video 7). We found that blocking cell division prevented ventral midline opening and significantly reduced midline sealing defects of *Mbt/PAK* mutant, to a level similar to control embryos (Fig.3c, d, Supplementary Video 7). On the other hand, ectopic expression of String/Cdc25 induced embryo-wide synchronous cell division at stage 7, including germ band tissues that are normally non-dividing at this stage (Fig.3e, Supplementary Video 8). Whereas control embryos displayed remarkable resilience to hyper-proliferation and maintained intact adhesion between cells after division, global induction of mitosis caused severe epithelial integrity defects in *Mbt/PAK* mutant embryos (Fig.3f, Supplementary Video 9). Based on these results, we propose that Mbt/PAK might be critical to establish stable adhesion after synchronous cell division, by promoting dynamic recruitment of Canoe/Afadin to newly established tricellular junctions after mitosis.

### Mutual dependency between Mbt/PAK and Canoe/Afadin

To explore molecular mechanisms of how Mbt/PAK regulates Canoe/Afadin mechanosensitivity, we started by pinpointing minimal protein features required for normal Mbt/PAK localization and function in embryos. As a Group-II p21-activated kinase, Mbt/PAK shares a well-conserved domain organization with the N-terminal Cdc42-Rac interaction/binding (CRIB) domain followed by the auto-inhibitory domain (AID) and the C-terminal catalytic domain with Serine/Threonine kinase activity (Extended Data Fig. 5a). *In vitro* kinase assays and cell culture studies on PAK4, a mammalian homolog of Mbt/PAK, demonstrated that Cdc42-GTP binding releases the auto-inhibition mediated by self-interactions between a pseudo-substrate (PS) motif and the kinase domain, and promoted its localization to bicellular junctions^47,56,57^ (Extended Data Fig. 5b). However, the mechanism and physiological functions of junctional recruitment of Group-II PAK remain largely unclear. By investigating the sole member of Group-II PAK in *Drosophila*, we now provide *in vivo* evidence to support a model that Cdc42-binding relieves Mbt/PAK auto-inhibition by exposing a previously uncharacterized Adherens Junction-localizing Motif (AJM), which mediates interactions with Canoe/Afadin and promote mechanosensing at tricellular junctions (Fig.4b).

To examine the impact of auto-inhibition on Mbt/PAK localization *in vivo*, we generated a deletion variant (ΔPS) lacking the R-P-L-P motif (aa. 54-57) that was shown to mediate auto-inhibition by serving as a pseudo-substrate for the kinase domain (Extended Data Fig. 5a, b). Compared to wild type, Mbt/PAK-ΔPS becomes highly enriched at tricellular junctions (Fig.4a second panel), with two-fold increase in TCJ ratio and junction-cytoplasm ratio (Fig.4c, d), despite of similar protein expression level as Mbt/PAK-WT (Extended Data Fig. 6g). This result suggests that release of auto-inhibition potentially activated a mechanism for Mbt/PAK tricellular junction recruitment. Next, to examine the requirement of Cdc42 interaction, we introduced point mutations (H19L, H22L, named as ΔCdc42) to the CRIB domain that supposedly disrupts Cdc42 binding (Extended Data Fig. 5a). Mbt/PAK-ΔCdc42 localized exclusively to the cytoplasm with no apparent junctional localization (Fig.4a, fourth panel, 4c), consistent with a requirement for Cdc42-binding to release the auto-inhibition state of Mbt/PAK (Fig.4b). Interestingly, further deletion of the PS motif within the ΔCdc42 variant (ΔPS+ΔCdc42) rescued junction localization (Fig.4a fifth panel, 4c), therefore bypassed the requirement of Cdc42-binding. These results suggest that Cdc42-binding is likely required for releasing auto-inhibition, but not directly for junctional localization. Release of auto-inhibition normally enhances kinase activity, therefore we next tested whether kinase-activity is required for Mbt/PAK junction localization. Surprisingly, a kinase-inactivating point mutation^49^ (T525A) of Mbt/PAK (Extended Data Fig. 5a) exhibited increased junction localization and tricellular enrichment (Fig.4a third panel, 4c, d). Based on the auto-inhibition model, the enhanced localization likely resulted from release of auto-inhibition through reduced binding to the PS motif, similar to the effect of removing PS motif entirely. The findings that neither Cdc42-binding or kinase-activity is directly required for junctional localization raise the possibility that might exist additional junction-localization motif(s) within Mbt/PAK that are normally hidden but become exposed after release of auto-inhibition to promote junction recruitment (Fig.4b).

To search for the unknown junction localization motif, we performed protein-wide structure-function analysis by generating a series of Mbt/PAK deletion variant and examined their localization in living embryos (Extended Data Fig. 5a). First, we found that Mbt/PAK^1–345^ is sufficient for junctional localization whereas the rest of the protein region (Mbt/PAK^Δ1–345^) localized entirely in the cytoplasm (Extended Data Fig. 5a, c, f), suggesting that the junction-localizing motif likely resides within the N-terminal region. Next, sequential deletion of this region at 50-aa intervals revealed that a short region (aa. 58-100) immediately after the PS motif is essential for junctional localization (Extended Data Fig. 5a, d, g). To examine the requirement of this motif for the localization of full-length Mbt/PAK, we selectively deleted this region but find that this variant (Mbt/PAK ^Δ54–100^) localized normally at junctions (Extended Data Fig. 5a, e, h). This result suggests that there might exist a secondary junction localizing motif. To examine this possibility, we generated another round of sequential deletions within the full-length Mbt/PAK (Extended Data Fig. 5a). Compared with the normal localization of Mbt/PAK ^Δ54–300^, further deletion of another 45 amino acid (Mbt/PAK ^Δ54–345^ or Mbt/PAK ^ΔPS+ΔAJM^) rendered the protein completely cytoplasmic (Extended Data Fig. 5e, h). Therefore, we identified two adherens junction localizing motifs (AJM): one immediately after the PS motif and the other just before the kinase domain. The location of these two AJM implies a model that release of auto-interaction between the PS motif and kinase domain exposed the “buried” AJM, which becomes accessible to a junction recruitment mechanism (Fig.4b).

To identify potential anchors for Mbt/PAK junctional recruitment, we examined Mbt/PAK localization in embryos lacking canonical adherens junction components. Unexpectedly, Mbt/PAK retained significant junction localization after alpha-catenin knockdown, a core component required for cadherin complex integrity and cell-cell adhesion (Extended Data Fig. 6a, b). In contrast, Canoe/Afadin knockdown resulted in embryo-wide delocalization of Mbt/PAK into the cytoplasm (Fig.4e, f, Extended Data Fig. 6a, b). In addition, Canoe/Afadin knockdown did not affect Mbt/PAK protein stability (Fig.4g). These results suggests that Canoe/Afadin might be directly required for Mbt/PAK junction recruitment. One possible mechanism is that Canoe/Afadin knockdown might affect Cdc42 activity and therefore locked Mbt/PAK in a constitutive auto-inhibited state. To examine this possibility, we tested the localization of Mbt/PAK-ΔPS, a variant that bypasses requirement of Cdc42 binding and releases auto-inhibition, in control and in *canoe*-RNAi embryos. We found that Mbt/PAK-ΔPS is similarly delocalized to the cytoplasm as Mbt/PAK-WT in *canoe*-RNAi embryos, despite of normal expression levels (Extended Data Fig. 6c-e), suggesting that Canoe/Afadin is likely directly required for Mbt/PAK junction recruitment. In addition, based on the model, exposing the AJM through deletion of PS motif should allow constitutive interactions between Mbt/PAK with Canoe/Afadin (Fig.4b). Indeed, spatial correlation analysis demonstrated that the localization of Mbt/PAK-ΔPS is almost indistinguishable from Canoe/Afadin (Extended Data Fig. 6f, h). Together, we propose that Mbt/PAK is an obligatory partner of Canoe/Afadin and they are mutually dependent for tricellular localization and functions to maintain epithelial integrity.

To understand physiological relevance of the self-inhibition mechanism to control Group-II PAK activity, we examined whether Mbt/PAK variants that specifically blocks each regulatory step can recapitulate tricellular localization or rescue adhesion defects when expressed in Mbt/PAK homozygous mutant. First, selectively blocking Cdc42 binding prevented Mbt/PAK junction localization and Mbt/PAK^ΔCdc42^ failed to rescue adhesion defects (Fig.4h, i, third panel), suggesting that Cdc42 might serve as a critical regulatory signal to modulate Mbt/PAK function *in vivo*. In comparison, release of the auto-inhibitory state by removing the PS motif rescued both localization and function of Mbt/PAK (Fig.4h, i, fourth panel). However, additional removal of the AJM motif rendered the protein completely cytoplasmic and Mbt/PAK^ΔPS+ΔAJM^ failed to rescue cell adhesion defects despite released from auto-inhibition (Fig.4h, i, fifth panel), validating an essential role of junction recruitment for Mbt/PAK function. Auto-inhibition is a common mechanism to regulate serine/threonine kinase activities. To our surprise, the Kinase-Dead (T525A) form of Mbt/PAK recapitulated tricellular localization and completely rescued homozygous mutant phenotypes (Fig.4h, i, sixth panel). In addition, expression of Mbt/PAK^T525A^ reverted Canoe/Afadin localization to normal TCJ enriched patterns in mutant embryos (Fig.4j, k) and rescued ventral midline sealing defects (Fig.4j, l). These results strongly support an unexpected conclusion that junctional localization, but not kinase activity, is required for Mbt/PAK function to modulate Canoe/Afadin mechanosensitivity and tricellular adhesion *in vivo*.

### Temperature-sensitive tactoid-shaped Canoe/Afadin junctional condensation underlies mechanical desensitization

In light of observations that Canoe/Afadin formed tactoid-shaped cortical structures in the absence of Mbt/PAK, particularly in mesectoderm cells after synchronous cell division (Fig.2b, Extended Data Fig. 4d), we tracked the pattern of Canoe/Afadin junctional distribution in control, Mbt/PAK-RNAi and *Mbt/PAK* mutant embryos before and after embryonic mitotic cycles, from stage-7 to stage-10. (Extended Data Fig. 7a). We also developed a new metric named JEDI (Junctional Equal Distribution Index) to quantitatively determine whether a junctional protein is uniformly distributed or unevenly concentrated at one or two edges within a single cell (Fig.5b, Table S3). Strikingly, the percentage of cells with junctional Canoe/Afadin condensation considerably increased from pre-mitotic to post-mitotic stage in Mbt/PAK-RNAi (13% to 45%) or *Mbt/PAK* mutant embryos (19% to 51%) (Extended Data Fig. 7b, c). Whereas the tactoid-shaped Canoe/Afadin condensation was never observed in control embryos (Extended Data Fig. 7b, c), Canoe/Afadin disproportionally accumulate at one or two edges of each cell at an embryo-wide scale after mitosis at stage 10 (Fig.5a, Supplementary Video 10), resulting in a three-fold increase in JEDI values (Fig.5b). Together, these results implicate mitotic cycles as one contributing factor to the aberrant formation of tactoid-shaped Canoe/Afadin cortical structures. In other words, there might be a unique requirement for Mbt/PAK in proliferative tissues to prevent aberrant Canoe/Afadin localization.

The tactoid-shape of Canoe/Afadin cortical structures prompted us to examine a possibility that they might represent a liquid- or gel-like phase-separated entity^58,59^, which normally formed as spherical granules in the cytoplasm^58,60^. To examine this possibility, we first analyzed the protein dynamics of Canoe/Afadin condensation directly in living embryos. Whereas the majority of Canoe/Afadin condensates is very stable with intensity fluctuations limited to less than 20% (Fig.5c, d, grey curves), kymographs of single cortical condensation events demonstrated that the structure can also be dynamic, exhibiting steady accumulation and dissipation patterns over time (Fig.5c, d, magenta and blue curves). This result suggests that there are exchanges in Canoe/Afadin molecules between the inside and outside of tactoid-shaped structures. To further probe molecular diffusion rates, we performed fluorescence recovery after photobleaching (FRAP) in live embryos. Compared to rapid recovery in control embryos, FRAP half-life of Canoe/Afadin increased by 2.5-fold in Mbt/PAK-RNAi embryos (Fig.5e, f). implicating a slower molecular diffusion rate of Canoe/Afadin when forming cortical condensates.

Slower FRAP recovery is often observed in fiber- or gel-like condensates *in vitro*, and the threshold concentration required for condensation is sensitive to environmental factors such as temperature, PH, and pressure^61–63^. To determine biophysical properties of Canoe/Afadin directly in living embryos, we chose temperature as the viable factor to manipulate. Unlike mammals, *Drosophila* embryo development is highly robust across a wide temperature range^64,65^. Therefore, we subjected embryos to a temporal temperature-shifting scheme and tracked endogenously Venus-tagged Canoe/Afadin localization over time. Whereas temperature shifts had no impact on Canoe/Afadin localization in control embryos, the tactoid-shaped condensates in Mbt/PAK-RNAi and Mbt/PAK mutant embryos notably dissolved (Fig.5g) and Canoe/Afadin regained tricellular junction enrichment after 10-minute incubation at 34°C, quantified as a decreased of JEDI (Fig.5h) and increase of TCJ ratio (Extended Data Fig. 7d). Remarkably, Canoe/Afadin condensates rapidly re-appeared after 10-minute cooldown at 22°C and fully reverted to pre-thermal shift levels after 20 minutes (Fig.5g, h, Supplementary Video 11), demonstrating that temporary shifting embryos to 34°C does not cause lethality or other cellular catastrophes. Together, the thermodynamics of Canoe/Afadin junctional condensates *in vivo* are consistent with features of a phase-separated entity.

Melting and cooling analysis have been commonly used to characterize phase transitions and define critical solution temperatures of molecules *in vitro*^66^. If even- and uneven-junctional distributions of Canoe/Afadin measured by JEDI values indeed reflect two distinctive phases, we should be able to utilize similar analysis to quantitatively characterize the thermodynamics of Canoe/Afadin, directly in living embryos. Similar to melting curve analysis, we subject the embryos to increasing shifting temperatures. By qualitative junction localization patterns and plotting JEDI values across different shifting temperatures, we found that Canoe/Afadin condensates dissolved in a temperature-sensitive manner (Fig. 5i, j). After shifting embryos back to 22°C, Canoe/Afadin condensates all re-appeared efficiently within 20 minutes (Fig. 5j), except the 38°C incubation that likely exceeded the upper limit of physiological temperature range of *Drosophila* embryos (Fig.5j, dark red). Next, by plotting the ratio of JEDI values between pre- and thermal-shift embryos against increasing temperatures, we found that Canoe/Afadin condensates exhibited a characteristic “melting” behavior, with critical solution temperature determined at around 29°C (Fig.5k). Lastly, taking advantage of the efficient re-condensation after cooling, we tracked the dynamics of single Canoe/Afadin condensation events over time at 22°C after 10-minute 36°C incubation. Living recordings and temporal plotting of JEDI values (normalized by time zero) clearly demonstrated a time-dependent phase-transiting behavior of Canoe/Afadin in embryos (Fig.5l), similar to a process *in vitro* when liquids solidify over time. Together, we provide evidence that thermodynamic properties of proteins can be analyzed directly in living embryos, and Canoe/Afadin condensation at cell junctions likely represents a two-phase system.

### A conserved N-terminal alpha-helix is essential for Canoe/Afadin condensation, localization, and function

Biomolecular condensates typically involve multivalent interactions with certain components serving as scaffolds and others as co-partitioned “clients”^61,62^. To examine whether Canoe/Afadin serves as scaffolds or depends upon other junctional proteins for condensation, we compared localization of major adherens junction components between control and Mbt/PAK-RNAi embryos. We found that E-cadherin, alpha-catenin, Ajuba, Sidekick and Echinoid that are previous reported to interact with Canoe/Afadin all localized similarly between control and Mbt/PAK-RNAi embryos, with no apparent junctional condensation (Extended Data Fig. 8a, c). In addition, although purified actin filaments can form tactoid-shaped phases^67^ and Mbt homolog PAK4 was shown to regulate actin cytoskeleton dynamics^68^, we found no signs of actin filament condensation in either control or Mbt/PAK-RNAi embryos (Extended Data Fig. 8a, c). Further, F-actin intensity is similar between junctions with- and without Canoe/Afadin condensates within the same cells (Extended Data Fig. 8b, inserts). These results suggest that Canoe/Afadin does not co-condensate with other major junction components or actin cytoskeleton and likely serves as the scaffold itself.

Therefore, we hypothesize that there might exist certain phase-promoting motifs within Canoe/Afadin that mediate condensation. To pinpoint such structural elements, we ectopically expressed a series of Canoe/Afadin deletion and mutated variants in control and Mbt/PAK-RNAi embryos. First, over-expressed full-length Venus-Canoe fully recapitulated tactoid-shaped condensation in Mbt/PAK-RNAi but not in control embryos (Fig.6a, c), suggesting that simply increasing protein concentration is not sufficient to drive junctional condensation. In contrast, deletion of the N-terminal region completely abrogated Canoe/Afadin condensation (Extended Data Fig. 9a-c). The N-terminal region contains two Rap1-binding domain (RBD) and a N-terminal structurally conserved but functionally unclear alpha-helix (Extended Data Fig. 9b). Selectively disrupting Rap1-binding by mutating four conserved lysine within the RBD (4KA) abolished Canoe/Afadin condensation, suggesting that Rap1-binding is necessary (Extended Data Fig. 9a-c). To examine whether Rap1-binding is sufficient for Canoe/Afadin condensation, we substituted the N-terminal region of Canoe/Afadin with the Rap1-binding domain of canonical Rap1 effector RALGDS (Extended Data Fig. 9b). However, RALGDS-RBD-Canoe did not regain the ability to form condensates in either control or Mbt/PAK-RNAi embryos (Extended Data Fig. 9a-c), implicating an additional role for the evolutionally conserved alpha-helix located N-terminal to the Canoe/Afadin-RBD (Fig.6d, Extended Data Fig. 9b). Indeed, deletion of the alpha-helix alone (1-24 amino acids) completely abrogated Canoe/Afadin junctional condensation (Fig.6a, c). Moreover, simply fusing the single alpha-helix with monomeric GFP (msVenus) resulted in embryo-wide cytoplasmic puncta formation (Supplementary Video 12). These puncta exhibit highly dynamic liquid-like behaviors to fuse into larger particles or split into smaller ones within seconds (Fig. 6b), consistent with a well-established activity of alpha-helices to induce multivalent self-interactions. In light of these observations, we also examined the other two alpha-helices at the C-terminal region of Canoe/Afadin but found no requirements for condensation (Extended Data Fig. 9a-c). Together, we uncovered a novel role for the conserved N-terminal alpha-helix, together with Rap1-binding, to promote Canoe/Afadin junctional condensation.

The N-terminal alpha-helix promotes aberrant Canoe/Afadin junctional condensation and rendered the protein less functional, in a condition when Mbt/PAK is down-regulated. Then what are physiological benefits for the extensive conservation of this structural element from worms to human (Fig.6d)? With observations that endogenous Canoe/Afadin exhibited a “edge-to-vertex” flow towards tricellular junction in the form of spherical structures, we hypothesized that an intermediate level of condensation mediated by the N-terminal alpha-helix might be required for Canoe/Afadin localization or function in wild type embryos, while Mbt/PAK serves as a gatekeeper to prevent excessive condensation. Therefore, we tested whether the Canoe/Afadin variant lacking the N-terminal alpha-helix (ΔαH1) can rescue *canoe* mutant embryos. In contrast to full-length (WT), Canoe-ΔαH1 failed to enrich at tricellular junctions despite of normal bicellular localization (Fig.6e, f). Dual-color live imaging revealed that deletion of the alpha-helix abrogated correlations between Canoe/Afadin and actomyosin activity (Fig.6e insets, 6g). In other words, Canoe-ΔαH1 becomes insensitive to mechanical force, despite the presence of Mbt/PAK and absence of aberrant condensation. Functionally, Canoe-ΔαH1 failed to rescue adhesion defects in *canoe* mutant embryos, characterized as epithelial gaps between myosin cables (Fig.6h, blue area, Supplementary Video 13). These results demonstrated that N-terminal alpha-helix is indeed critical for normal tricellular recruitment and function of Canoe/Afadin. Together, self-interaction might be a general requirement for the edge-to-vertex transport of Canoe/Afadin to tricellular junctions, with Mbt/PAK serving as a co-factor to keep this activity in check to prevent excessive condensation (Fig. 6i).

## Discussion

Our studies provide direct evidence of the physiological regulation of mechanosensitivity through the modulation of endogenous protein condensational states. In non-dividing cells, Canoe/Afadin is directly transported from bicellular to tricellular junctions to enhance adhesion under tension. During synchronous cell division, the temporary displacement of junctional Mbt/PAK desensitizes Canoe/Afadin mechano-response at metaphase. After mitosis, the restoration of Mbt/PAK junction localization promotes Canoe/Afadin recruitment to newly formed tricellular junctions. In the absence of this mechanism, Canoe/Afadin aggregates into tactoid-shaped condensates at the cell cortex, rendering them unresponsive to mechanical forces. This impedes the re-establishment of adhesion post-division, leading to significant epithelial integrity defects in proliferative tissues. At the molecular level, an auto-inhibitory mechanism of Mbt/PAK acts as an “on-off” switch for Canoe/Afadin mechanosensitivity by regulating protein condensation and cortical mobility (Model in Fig.6i).

Cell adhesion, tension and division are three interconnected processes crucial for animal development. A “Knee-jerk” mechanism was proposed to uphold cell adhesion during movements, by coupling of adhesion strength with mechanical force^3,18,26,69^. Here we propose a potential decoupling mechanism to temporarily dampen mechanosensitivity of tricellular junctions during synchronous cell division. A similar phenomenon has been observed at bicellular junctions, as evidenced by the diminished mechano-response of Vinculin in mitotic cultured cells^43^. The precise physiological rationale for desensitizing mechano-response during mitosis is not entirely clear. Cell division relies on cytoskeletal force to drive extensive cell shapes such as mitotic rounding and cleavage furrow formation^5,28,29^. Scaling up adhesion strength accordingly during these events might be counteractive to required cell shape changes for mitosis. For example, the localized reduction of E-cadherin coincides with cleavage furrow progression^10^, while a constitutive mechano-sensitive alpha-catenin hinders mitotic rounding^43^. Therefore, the mechanism we described here could offer an *in vivo* solution that reconciles conflictive demands on cytoskeletal force levels by temporarily decouple adhesion from mechanical inputs and thereby enables cell shapes changes during mitosis.

Phase separation have been recognized to be involved in many aspects of cellular and signaling events. However, detecting phase transitions *in vivo* with molecules at endogenous concentrations remains uncommon^58,60,70^. Here we observed that endogenously-tagged Canoe/Afadin forms spherical or tactoid-shaped condensates within living *Drosophila* embryos, with these transitions facilitated by an obligatory co-factor Mbt/PAK. When in spherical form, Canoe/Afadin moves directly from bicellular junctions towards tricellular junctions under actomyosin flow, strengthening adhesion under tension. A liquid-liquid phase separation mechanism has been reported for a tight junction protein named ZO-1^71^, which transports from the non-junctional cell cortex to bicellular junctions in droplet form and confers mechanosensitivity of tight junctions^23^. However, whether phase-separation can be regulated by physiological cues to modulate mechanosensitivity remains unknown. Our findings demonstrated that down-regulation of Mbt/PAK caused transition of Canoe/Afadin into tactoid-shaped condensates, reducing molecule diffusion rates and cortical mobility, thereby decreasing mechanosensitivity. Importantly, this mechanism is utilized by synchronously dividing cells within mitotic domains to dynamically remodel cell-cell adhesion during embryonic proliferation.

Biomolecular condensation *in vitro* can be modulated by physical parameters such as salt, PH, and temperature^61–63^. Whereas living organisms are known to sense and respond to changes in these factors, an important conceptual gap exists regarding whether modification of cellular protein condensation contributes to osmosensing or thermosensing. Taking advantage of the fact that *Drosophila* embryogenesis is highly robust to shifting temperatures, we examined Canoe/Afadin condensational states at varying temperatures within the physiological range. Remarkably, tactoid-shaped Canoe/Afadin condensates rapidly dissolved at 30°C or higher, but dynamically re-established after cooldown to 22°C. Canoe/Afadin regained tricellular localization after condensate dissolution, despite of the absence of Mbt/PAK. In other words, temperature variations could bypass the requirement of important genetic information for regulating cellular activities during embryogenesis. Allosteric or physical switches of protein conformations mediates signal transduction in response to biochemical or mechanical inputs. Similarly, the switch of protein condensational states might present a regulatory hinge point for thermal signaling, an exciting emerging concept exploring temperature variations as a signal to drive biological events^72^. Similar to mechanical signaling, it is important to identify cellular sensors and responders to thermal variations. Combining powerful genetics, diverse live imaging tools and a wide-range thermal tolerance, *Drosophila* embryos presents a unique system to systematically screen for molecules involved in thermal signaling.

## Methods

### *Drosophila* stocks and genetics

All embryos were collected in cylinder cages at 25 °C during the day or room temperature overnight and processed for imaging. Room temperature of the imaging facility was set at 22°C. *Drosophila* genotypes of each figure and supplemental figure were listed in Table S1. Catalog number of all commercially available stocks, *Drosophila* lines gifted from other labs and made in this study were listed in Table S2. *Drosophila* lines expressing shRNA for genetic knockdown analysis were obtained from Tsinghua Fly Center (THFC) and Bloomington Drosophila Stock Center (BDSC). *Drosophila* lines expressing Venus-tagged proteins at endogenous locus were obtained from the Kyoto stock center as part of the Cambridge Protein Trap Insertion (CPTI) collection. The following stocks were generous gifts including the (*ubi-Ecadherin-GFP*, *sqh(MRLC)-mCherry),* (*nanos::Cas9)*, (*y,w,nanos::phic31*;*attp40)*, (*y,w,nanos::phic31*;*attp2), (UAST-ShroomA)*, (Ajuba-GFP, BAC) and (*matatub67; Spider-GFP, sqh-mCherry*) from Jennifer Zallen (Sloan Kettering Institute, NY, USA), the *cno*^R2^ mutant allele from Mark Peifer (University of North Caroline Chapel Hill, NC, USA), the (*matatub67, sqh-GFP; matatub15, GAP43-mCherry*) from Adam Martin (Massachusetts Institute of Technology, Cambridge, MA, USA), the (*matatub67*; *matatub15)* Gal4 stock from Daniel St. Johnston (University of Cambridge, Cambridge, UK). Genomic recombination was used to generate (*matatub67*, *MRLC-mCherry*), (*matatub67*, *Tubulin-mCherry*), (*matatub67*, *UASp-Venus-RALGDS-RBD)*, and (*matatub67*, *UAS-P{TRiP.HMS04518}attP40*) on the same chromosome. Briefly, males carrying desired transgenes were crossed to *matatub67* virgin females, individual male F1 progeny was crossed to *Sp/Cyo* virgin females for 5 days before sacrificed for PCR screens. Primers for detection of *matatub67* are 5’-GCGCCAAGTGTCTGAAGAACAAC(Fwd) and 5’-CCAAGGGCATCGGTAAACATCTG (Rev). Primers for detection of mCherry are 5’-GCCAAGCTGAAGGTGACCAAG (Fwd) and 5’-GGTGTAGTCCTCGTTGTGGGA (Rev). Primers for detection of msVenus are 5’-GATAGCACTGAGAGCCTGTTCAC (Fwd) and 5’-GGTCACGAACTCCAGCAGGA (Rev). Primers for detection of *TRiP.HMS04518 (Mbt/PAK shRNA)* are 5’-ACCAGCAACCAAGTAAATCAAC (Fwd) and 5’-TAATCGTGTGTGATGCCTACC (Rev). Progenies of screened males with the presence of both *matatub67* and *mCherry* or msVenus or *TRiP.HMS04518* were selected to make stable stocks. The maternal *canoe* mutant embryos analyzed in this study were generated using FLP recombinase dominant female sterile system, described in detail previously. The *mbt* mutant embryos are homozygous viable and therefore, all *mbt* mutant embryos analyzed are depleted of both maternal and zygotic Mbt/PAK proteins.

### *Drosophila* CRISPR mutant generation

For generation of CRISPR/Cas9 mutants of the *Mbt* (CG18582) gene, two gRNAs were expressed ubiquitously using the pCFD4 plasmid backbone. gRNAs were designed using the target finder tool of flycrispr.org website with template genomic sequence retrieved from flybase.org website. gRNA#1 (5’-GCGGTAAAAGGGCAGAAATT) targeted the 5’UTR of *Mbt* locus and gRNA#2 (GCTGCTGCTGTTGAGCGAGG) targeted the first exon. The *pCFD4-Mbt-gRNA* plasmid was cloned by gibson assembly of BbsI-digested pCFD4 and PCR fragment amplified from pCFD4 using gRNA primers 5’TATATAGGAAAGATATCCGGGTGAACTTCGCGGTAAAAGGGCAGAAATTGTTTTAGAGCTAGAAATAGCAAG (Fwd) and 5’-ATTTTAACTTGCTATTTCTAGCTCTAAAACCCTCGCTCAACAGCAGCAGCGACGTTAAATTGAAAATAGGTC (Rev). The *pCFD4-Mbt-gRNA* plasmid was then injected at 80 ng/μL into *nanos::Cas9* embryos within one hour after egg laying. F0 adult flies were individually crossed to FM7 virgin flies. The F1 progeny of injected flies were screened by sequencing using primers covering both CRISPR sites (Fwd 5’-CACCACCTCTATTTCTGGCATCG; Rev 5’-GAGAGGTCTTCGGCAGGACT). One CRISPR mutant was identified with a single cut at gRNA#2 site resulting in two missing nucleotides within the Exon 1 and thereby frameshift mutations starting at amino acid L90 and an early stop codon at amino acid A179. This mutant was selected to establish a stable Drosophila line without the FM7 balancer and used as the *mbt* homozygous mutant for this study. (Extended Data Fig. 3d)

### Cloning and transgenic lines

To generate N-terminal Venus-tagged variants of Mbt/PAK, the full-length *mbt* open reading frame (Isoform-RA, 1917nt, not including the stop codon) was PCR-amplified using Phanta Max Super-Fidelity DNA Polymerase (Vazyme) from a *Drosophila* cDNA library and cloned into MluI-digested UASp N-terminal msVenus vector by Gibson assembly using ClonExpress Ultra One Step Cloning Kit (Vazyme). Mbt/PAK transgenes containing deletions and point mutations were cloned as two- or three-part Gibson assembly reactions using the same linearized UASp-N-msVenus vector and PCR fragments amplified from UASp-Venus-Mbt/PAK-WT. The entire open reading frame of all transgenes were sequence verified and injected into the attP2 sites on Chromosome III. The exact deletion and mutations were listed below: Mbt/PAK^ΔPS^ (aa.54-57 deleted); Mbt/PAK^ΔCdc42^ (H19 and H22 mutated to Leucine); Mbt/PAK^kinase-dead^ (T525 mutated to Alanine); Mbt/PAK^ΔCdc42+ΔPS^ (H19 and H22 mutated to Leucine, and aa.54-57 deleted); Mbt/PAK^ΔAJM+ΔPS^ (aa. 54-345 deleted); Mbt/PAK^Δ1–345^ (aa. 1-345 deleted); Mbt/PAK^1–345^ (aa. 346-639 deleted); Mbt/PAK^1–300^ (aa. 301-639 deleted); Mbt/PAK^1–250^ (aa. 251-639 deleted); Mbt/PAK^1–200^ (aa. 201-639 deleted); Mbt/PAK^1–150^ (aa. 151-639 deleted); Mbt/PAK^1–100^ (aa. 101-639 deleted); Mbt/PAK^1–57^ (aa. 58-639 deleted); Mbt/PAK^Δ54–100^ (aa. 54-100 deleted); Mbt/PAK^Δ54–150^ (aa. 54-150 deleted); Mbt/PAK^Δ54–150^ (aa. 54-150 deleted); Mbt/PAK^Δ54–200^ (aa. 54-200 deleted); Mbt/PAK^Δ54–300^ (aa. 54-300 deleted). To generate N-terminal mCherry-tagged variants of Mbt/PAK, coding region of mCherry was synthesized directly, coding region of Mbt/PAK variants were amplified from UASp-Venus-Mbt/PAK plasmids, and then both fragments were cloned into a linearized UASp backbone vector through a three-part Gibson assembly. To generate N-terminal Venus-tagged variants of Canoe/Afadin, transgenes containing deletions or point mutations were amplified from UASp-Venus-Canoe-WT with distinct sets of primers and cloned into MluI-linearized UASp N-terminal msVenus backbone. The exact deletion and mutations were listed below: Canoe/Afadin^ΔNTR^ (aa. 1-350 deleted); Canoe/Afadin^4KA^ (K57, K77, K274, K294 mutated to Alanine); Canoe/Afadin^alpha-helix-1^(aa. 1-20); Canoe/Afadin^Δalpha-helix-1^ (aa. 1-24 deleted); Canoe/Afadin^Δalpha-helix-2^ (aa. 1583-1612 deleted); Canoe/Afadin^Δalpha-helix-3^ (aa. 1618-1659 deleted). To generate the Rap1-GTP sensor, the Rap1-binding domain (RBD) of RALGDS (aa 793-914) was PCR amplified from pET28-MHL-His-RALGDS: N793-F914 (H. sapiens RALGDS, Addgene #25331, deposited by Cheryl Arrowsmith), cloned into a UASp N-terminal msVenus backbone using Gibson assembly, and inserted into the attP40 site on chromosome II. To generate RalgdsRBD-Canoe/Afadin^ΔNTR^ -Chimera, coding region of Ralgds-RBD and coding region of Canoe/Afadin^ΔNTR^ were cloned into UASp N-terminal msVenus vector via a three-part Gibson assembly reaction. All Canoe/Afadin variants were injected into the attP40 sites on Chromosome II.

### Still frame and time-lapse live imaging

Embryos were collected for 6 hours (for stage 7-8) at 25°C or overnight (for stage 9-10) at room temperature using apple juice plates with yeast paste. Embryo chorions were removed by adding 2ml 50% bleach to the plate and swirl for 90 seconds. Embryo/bleach mix was transferred into a 70 μm nylon cell strainer and washed thoroughly by deionized water for one minute. Dry the cell strainer with paper towels and transfer the embryo immediately into halocarbon oil-27 (Sigma) within a clean apple juice plate without yeast paste. Hand pick embryos at the correct stage using tweezers based on morphological features under bright field microscope and mount them in a 1:1 mixture of halocarbon oil 27 and 700 (Sigma) between a gas-permeable membrane (YSI) and 22X22 mm coverslip (Epredia). Still-frame live images were acquired on a Zeiss LSM980 confocal microscope with a Zeiss Plan-Apochromat 63X /1.4 oil-immersion objective (1 μm optical section and 0.5 μm z-steps), including Fig. 2a, 2h, 2i, 2j, 3c, 3e, 4a, 4e, 4h, 4j, 5a, 5g, 5i, 6a, 6e, 6h, Extended Data Fig. 2c, 2d, 3a (left), 5c, 5d, 5e, 6c, 7b, 8a, 9a. Dual-color still-frame and time-lapse images were acquired on a Nikon FLUOVIEW microscope equipped by a spinning disk module CSU-X1FW with a Olympus UPlanSApo 60X /1.35 oil-immersion objective (0.5 μm z-steps, 21 z-stacks taken for each time frame), acquired at 45-second intervals, including Fig. 1c, 1d, 1e, 1f, 2b, 3b, 5c, 6b, 6e (inset), Extended Data Fig. 1c, 3c, 4b, 4c, 4d, 6a, 6f. Maximum-intensity projections of 10-15 optical slides were generated for analysis using Fiji software. Time-lapse videos were bleach corrected using Fiji plugins by histogram matching.

**Fig. 2:**
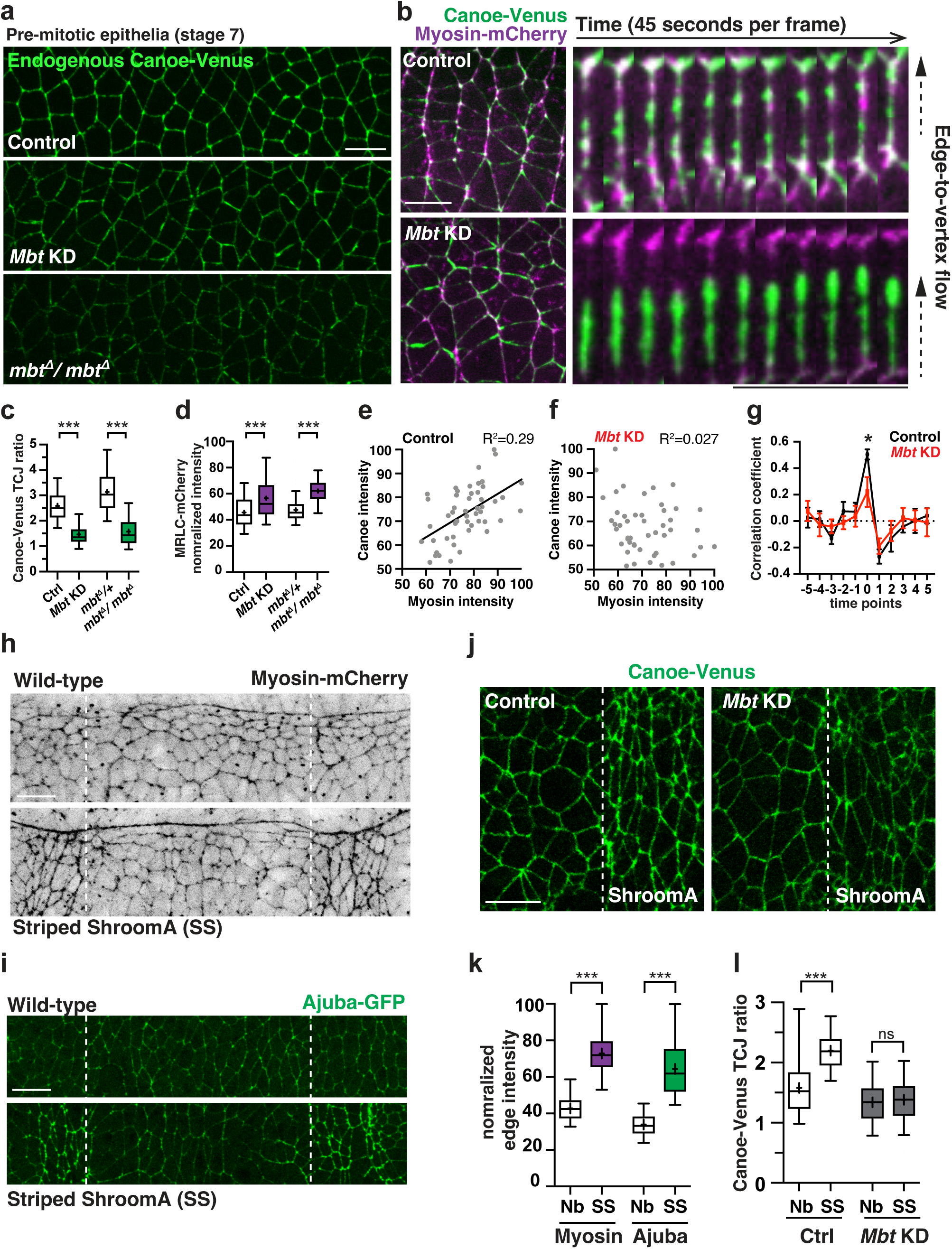
Mbt/PAK is required for Canoe/Afadin mechanosensitivity. **(a)** Still images of Canoe-Venus in control, *Mbt* knockdown and *mbt* homozygous mutant embryos at pre-mitotic stage 7. **(b)** Still images of a dual-color time lapse movie recording Canoe/Afadin-Venus together with Myosin-mCherry in control and *Mbt* knockdown embryos (Left), with kymographs of a single bicellular junction connecting two tricellular junctions shown on the right. Arrows with dotted line points the direction of edge-to-vertex protein flow. **(c, d)** Quantification of Canoe/Afadin TCJ enrichment and myosin junctional intensity in control, *Mbt* knockdown, *mbt* heterozygous (*mbt^Δ^*/+) and homozygous (*mbt^Δ^*/*mbt^Δ^*) mutant embryos. **(e, f)** Spatial correlation analysis of Canoe/Afadin and myosin at tricellular junctions in control and *Mbt* knockdown embryos. **(g)** Correlation coefficient for the rates of change in Canoe-Venus and Myosin-mCherry at TCJs in control and *Mbt* knockdown embryos. **(h)** Induction of local myosin activity in a mosaic manner through expression of Shroom-A in striped patterns (left and right side of dotted lines, bottom panel), compared to neighboring cells within the same embryo (middle, between dotted lines, bottom panel) or embryos without ShroomA expression (upper panel). **(i)** Induction of increase mechano-response at adherens junctions visualized by enhanced junctional localization of Ajuba-GFP specifically in cells with elevated myosin junctional intensities (left and right side of dotted lines, bottom panel). **(j, l)** Localization and quantification of Canoe/Afadin TCJ enrichment in stripes with increased myosin activity and neighboring cells with normal myosin activity, in control and *Mbt* knockdown embryos. **(k)** Quantification of junctional myosin and Ajuba localization in stripes with increased myosin activity and control neighboring cells. Boxes are 2^nd^ and 3^rd^ quartiles and whiskers are 5^th^ to 95^th^ percentiles; horizontal line is the median and “+” sign is the mean value. Mean±SD in (c), (d), (k) and (l); Mean±SEM between embryos in (g). ***P<0.001; *P<0.1; Not significant (ns) P>0.1. Welch’s t-test for all comparisons. Live embryos in all panels. Embryos are stage 7 in (a) and (b), stage 9 in (h), (i) and (j) for zygotic striped expression of ShroomA. All positioned with anterior left, ventral down. Scale bars, 10 μm.

**Fig. 3:**
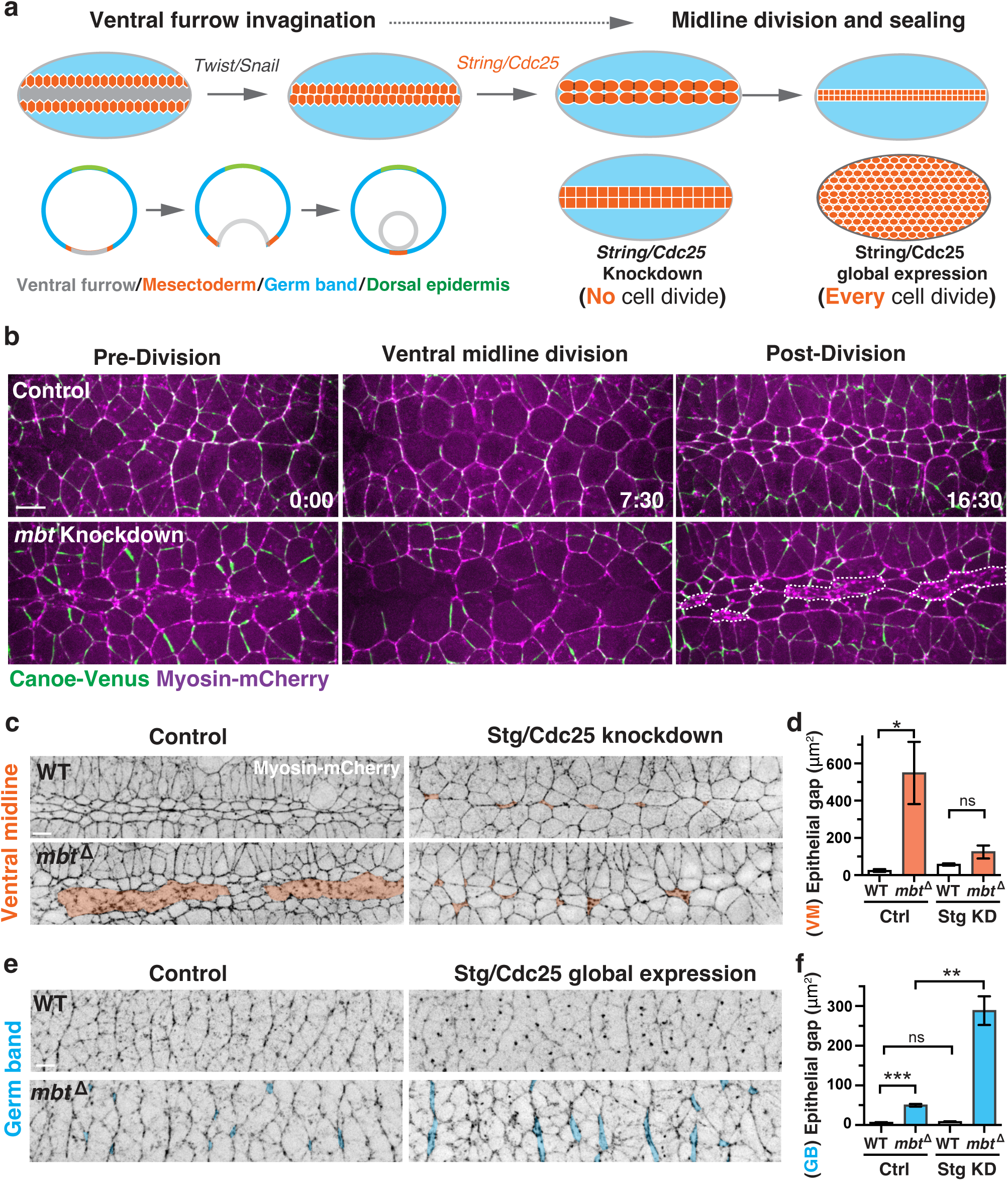
Mbt/PAK is required for re-establishing cell adhesion after mitosis in proliferative tissues. **(a)** Schematics of ventral midline formation, including furrow invagination initiated by patterned expression of Snail/Twist (grey region), parallel juxtaposition of mesectoderm cells (orange) along the midline after furrow closure, followed by synchronous cell division and adhesion re-establishment of mesectoderm cells, and eventually midline sealing (top panel). Sagittal view of the process shown at bottom left. Effects of decreasing or increasing protein levels of cell cycle regulator String/Cdc25 on embryonic divisions illustrated at bottom right. **(b)** Still images of a dual-color live imaging of Canoe/Afadin-Venus (green) and Myosin-mCherry (magenta) during ventral midline formation in control (top) and *Mbt* knockdown embryos (bottom). Epithelial gaps along the ventral midline delineated by dotted lines. **(c)** Still images of Myosin-mCherry in wild type (top) and mbt mutant embryos (bottom), with (right) or without (left) expression of shRNAs against Stg/Cdc25 driven by GAL4/UAS system. Epithelial gaps along the ventral midline delineated by orange coloring. **(d)** Epithelial adhesion defects quantified by the area of gaps between cells in wild type and mbt mutant embryos, with (Stg KD) or without (Ctrl) expression of shRNAs against Stg/ Cdc25. **(e)** Still images of Myosin-mCherry in wild type (top) and mbt mutant embryos (bottom), with (right) or without (left) ectopic expression of HA-tagged Stg/Cdc25. Epithelial gaps in the germ band tissue delineated by blue coloring. **(f)** Epithelial adhesion defects quantified by the area of gaps between cells in wild type and *mbt* mutant embryos, with (Stg OE) or without (Ctrl) ectopic expression of HA-tagged Stg/ Cdc25. Mean±SEM between embryos in (d), (f). ***P<0.001; **P<0.01; *P<0.1; Not significant (ns) P>0.1. Welch’s t-test for all comparisons. Live embryos in all panels. All positioned with anterior left, ventral down. Scale bars, 10 μm.

**Fig. 4:**
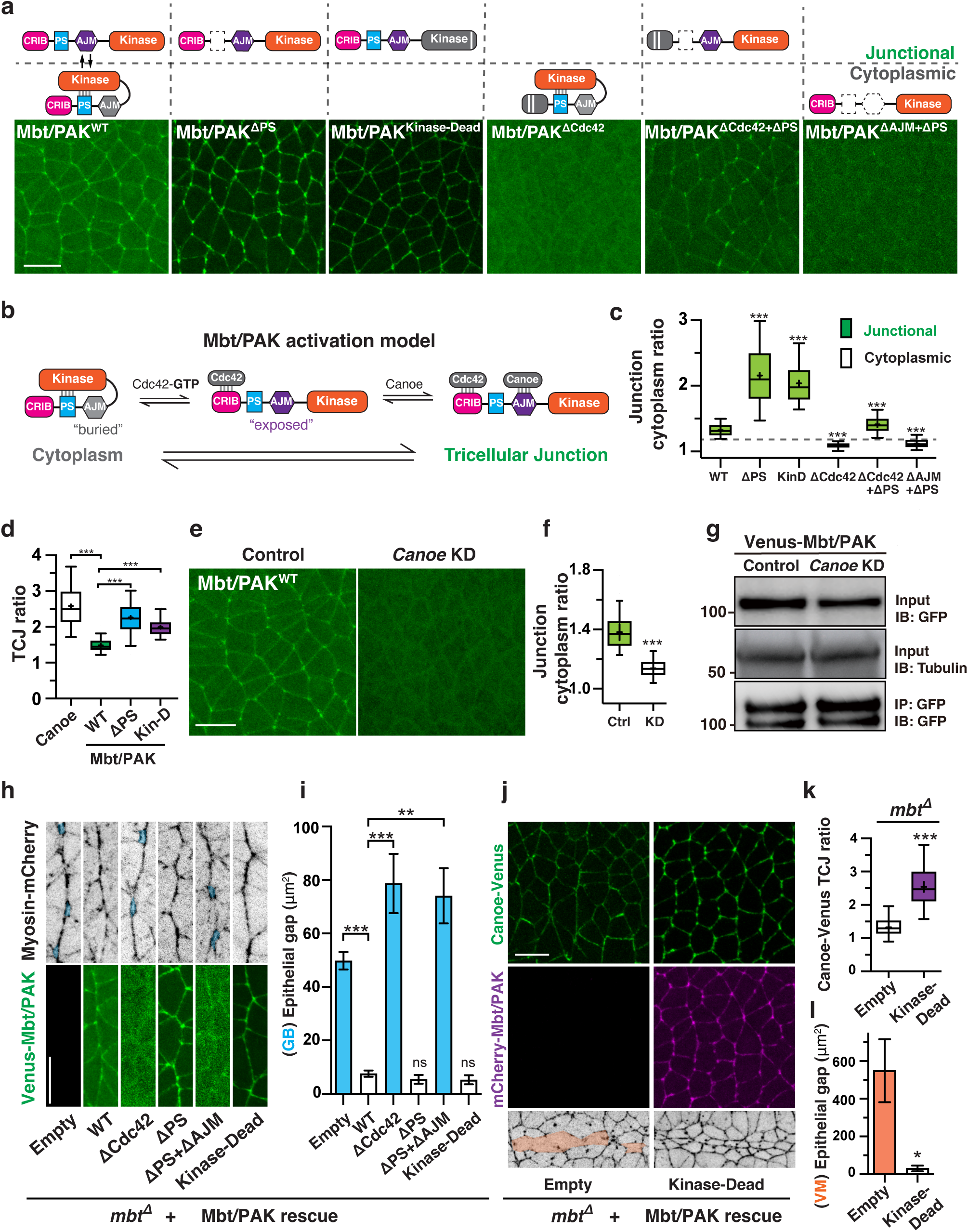
Mutual dependency between Mbt/PAK and Canoe/Afadin in localization and functions. **(a)** Localization of Venus-tagged ectopically expressed Mbt/PAK wild type or variant transgenes in pre-mitotic stage 7 embryos using the GAL4/UAS system. Schematics of domain organizations for each transgene shown above, with junction-localized ones at the upper panel and cytoplasm-localized at the bottom. **(b)** The updated model of Mbt/PAK activation: binding of Cdc42-GTP to the CRIB domain releases Mbt/PAK from the auto-inhibition state, which is mediated by a self-interaction between the Pseudo-Substrate (PS) (aa. 54-57) motif and kinase domain (known). Release of auto-inhibition exposes a hidden Adherens Junction localizing Motif (AJM), which mediates Mbt/PAK junctional recruitment in a Canoe/Afadin-dependent manner (this study). **(c)** Junctional enrichment of Mbt/PAK transgenes, quantified by the ratio between junctional and cytoplasmic intensities. **(d)** Comparison of tricellular enrichment between Canoe/ Afadin and Mbt/PAK-WT or variants, measured by TCJ ratios. **(e, f)** Localization of wild type Venus-Mbt/ PAK in control and *canoe* knockdown embryos (e), quantified by junction-cytoplasm ratio (f). **(g)** Ectopically expressed Venus-Mbt/PAK-WT in embryos using the GAL4/UAS system was immunoprecipitated from 0-5 hrs control or *canoe* knockdown embryos with an antibody to GFP and immunoblotted with antibodies to GFP (bottom). Input lysate was blotted with antibodies to GFP (top) and alpha-tubulin (middle). **(h)** Still images of Myosin-mCherry (top) in *mbt* homozygous mutant embryos expressing Venus-Mbt/PAK transgenes, with the localization of each Venus-tagged Mbt/PAK variant shown at the bottom panel. **(i)** Epithelial adhesion defects quantified by the area of gaps between cells in *mbt* homozygous mutant embryos expressing different Venus-Mbt/PAK transgenes. **(j, k)** Still images and TCJ ratios of Canoe-Venus (j, top) in *mbt* homozygous mutant embryos with (right) or without (left) expressing mCherry-Mbt/ PAK-Kinase-Dead. **(j, l)** Still images of Myosin-mCherry (j, bottom) and quantifications of epithelial gaps along the ventral midline (l) in *mbt* homozygous mutant embryos with (right) or without (left) expressing Venus-Mbt/PAK-Kinase-Dead. Boxes are 2^nd^ and 3^rd^ quartiles and whiskers are 5^th^ to 95^th^ percentiles; horizontal line is the median and “+” sign is the mean value. Mean±SD in (c), (d), (f), (k); Mean±SEM between embryos in (i), (l). ***P<0.001; **P<0.01; *P<0.1; Not significant (ns) P>0.1. Welch’s t-test for all comparisons. Live embryos in all panels. All positioned with anterior left, ventral down. Scale bars, 10 μm.

### Embryo temperature-shift and live imaging

Procedures for live images before temperature shifting is the same as normal live imaging as described above. Next, slides with mounted embryos were incubated inside the PECON incubator and heating chamber -P-S compact, with temperature controlled by the TempModule-S. The chamber was put aside on the desk instead of on top of objective lens, so that the bottom lens hole of the chamber can be sealed completely with tape to prevent temperature fluctuations. The temperature of chamber was controlled by the Zeiss software ZEN and set to desired temperature (34°C for example) 10 minutes before the incubation to minimize fluctuations. Embryos were incubated for 10 minutes, and then taken out immediately and put on stage for live imaging, with this image labeled as 34°C 10min. After this image taken, embryos remained on the stage and another two live images were taken after 10 minutes and 20 minutes, labeled as 34°C 10min+22°C 10min and 34°C 10min+22°C 20min. Procedures are the same for experiments with different shifting temperatures as in Fig. 5i, with “post” images acquired at 20min after temperature shift.

**Fig. 5:**
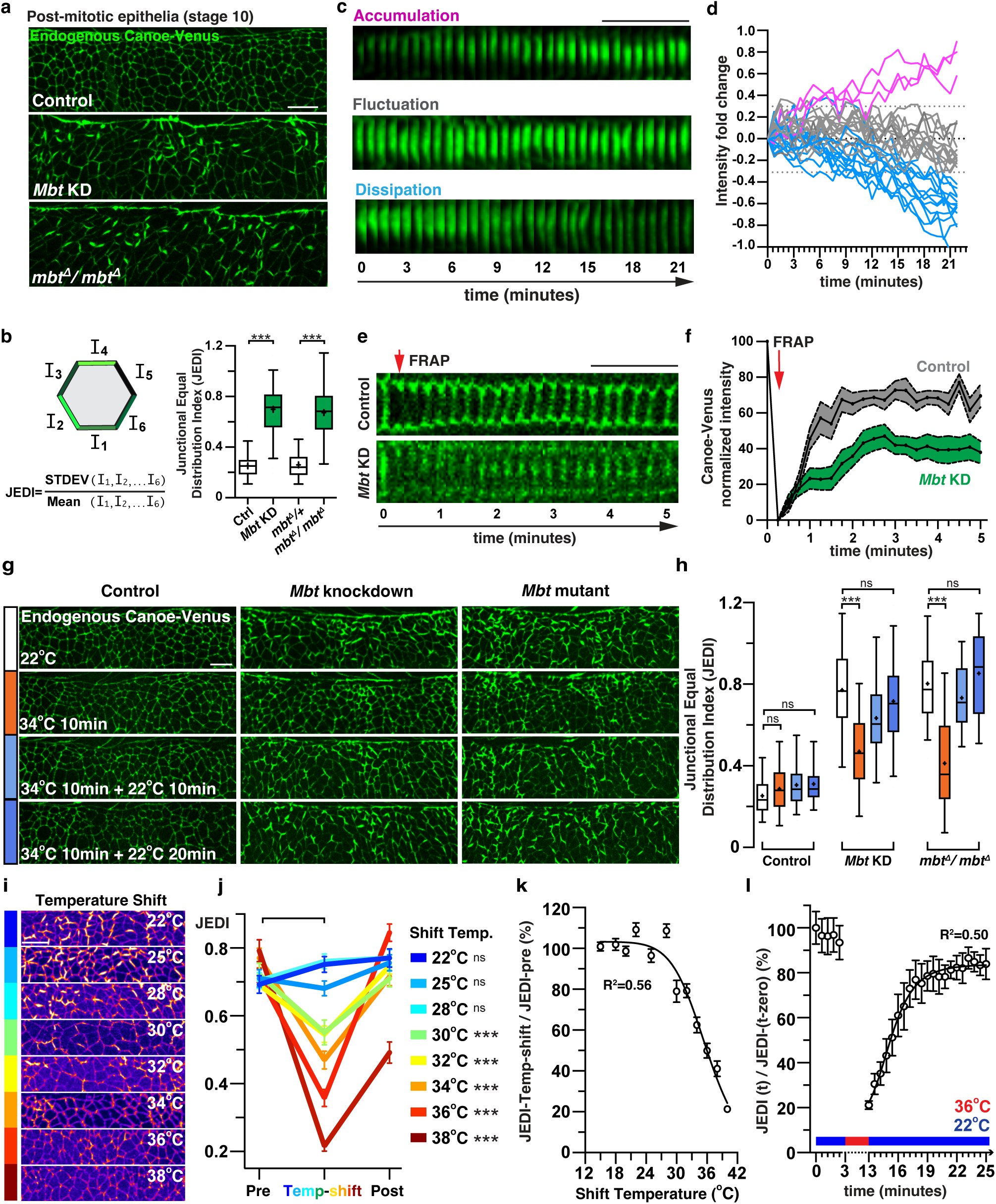
Temperature-sensitive cortical condensation of Canoe/Afadin. **(a)** Localization of Canoe-Venus in control (top), *Mbt* knockdown (middle) and *mbt* homozygous mutant (bottom) embryos at post-mitotic stage 10. **(b)** Schematic and calculation equation of JEDI, a metric to quantify protein distribution at junctions (left), and JEDI values in control, *Mbt* knockdown, heterozygous (*mbt^Δ^*/+) and homozygous (*mbt^Δ^*/ *mbt^Δ^*) mutant embryos (right). **(c, d)** Kymographs (c) and mean intensities fold changes over time (d) of Canoe/Afadin-Venus condensation over time, categorized into three distinctive patterns: Accumulation (> 0.3-fold increase, magenta), Fluctuation (< 0.3-fold increase or decrease, grey) and Dissipation (> 0.3-fold decrease, blue). **(e, f)** Kymographs of Canoe-Venus before and after bleaching over time at 20 second intervals (e) and quantification of Canoe-Venus diffusion rate using fluorescence recovery after photobleaching (f). **(g, h)** Localization and JEDI values of Canoe-Venus in control, *Mbt* knockdown and *mbt* homozygous mutant embryos under a temperature-shifting regime. Embryos firstly imaged at 22°C (white), shifted to 34°C for 10 minutes and then imaged the second time (orange), shifted back to 22°C and imaged the third (blue) and fourth (dark blue) time after another 10 and 20 minutes. **(i)** Localization of Canoe-Venus in *Mbt* knockdown embryos under different temperatures, colored by Fire LUT using ImageJ to highlight condensation patterns. **(j)** JEDI values of Canoe-Venus in *Mbt* knockdown embryos under the same temperature-shifting regime, with different shifting temperatures. Pre, images taken at 22°C before temperature shift; Post, images taken at 22°C 20 minutes after temperature shift. Statistical comparisons of JEDI values between Pre- and Temp-shift embryos shown aside of different shifting temperatures. **(k)** A melting curve analysis to determine the critical solution temperature of Canoe-Venus condensates, using JEDI values at different temperatures as a proxy for phase changes. **(l)** Condensation recovery dynamics measured as changes in normalized JEDI values by time zero over time in a temp-shift regime with shifting temperature set at 36 °C. Boxes are 2^nd^ and 3^rd^ quartiles and whiskers are 5^th^ to 95^th^ percentiles; horizontal line is the median and “+” sign is the mean value. Mean±SD in (b) and (h); Mean±SEM between cells in (f), (j), (k) and (l). ***P < 0.001; Not significant (ns) P > 0.1. Welch’s t-test for all comparisons. Live embryos in all panels. All positioned with anterior left, ventral down. Scale bars, 10 μm.

**Fig. 6:**
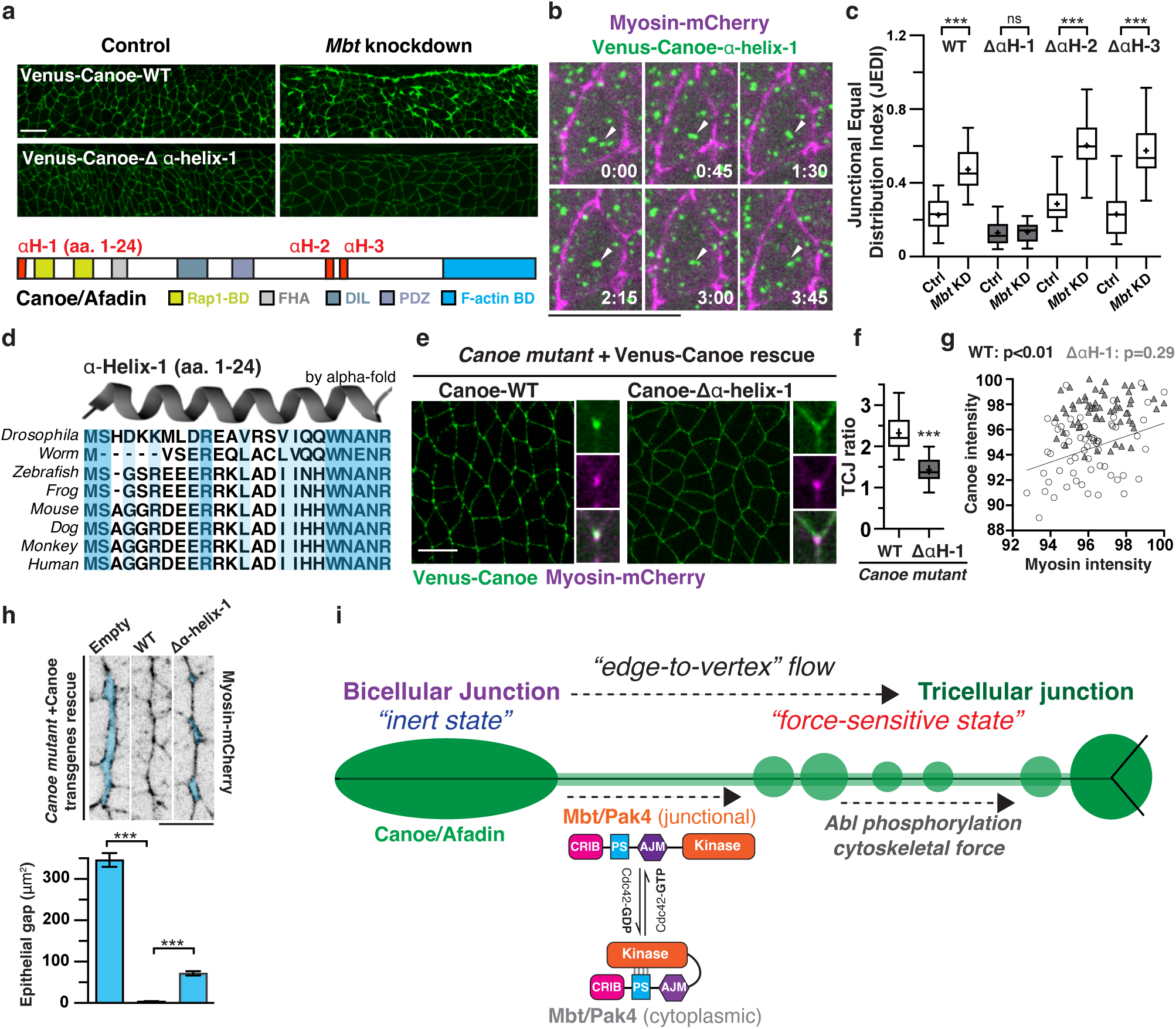
A conserved N-terminal alpha-helix is essential for Canoe/Afadin condensation and mechanosensitivity. **(a)** Localization of ectopically expressed Venus-Canoe-WT and Venus-Canoe-Δα-helix-1 in control and *mbt* knockdown embryos at post-mitotic stage 10, with domain organization of full-length Canoe shown below. **(b)** Still images of a dual-color time lapse movie recording Venus-Canoe-α-helix-1 (green) and Myosin-mCherry (magenta), note the merging and splitting of cytoplasmic puncta, arrowheads. **(c)** JEDI values of Venus-Canoe-WT, Venus-Canoe-Δα-helix-1, Venus-Canoe-Δα-helix-2, and Venus-Canoe-Δα-helix-3 in control and *mbt* knockdown embryos. **(d)** Protein sequence alignment of the first alpha-helix of Canoe/Afadin in different model organisms, with the predicted structure in *Drosophila* by alpha-fold shown above. **(e, f)** Localization (e), TCJ ratio (f) of Venus-Canoe-WT and Venus-Canoe-Δα-helix-1 expressed in *canoe* mutant embryos at pre-mitotic stage 7, with insets showing merged images with Myosin-mCherry. **(g)** Spatial correlation analysis between Myosin-mCherry and Venus-Canoe-WT (circles) or Venus-Canoe-Δα-helix-1 (triangles) expressed in *canoe* mutant embryos. **(h)** Still images of Myosin-mCherry (top) and quantifications of epithelial gaps (bottom) in the germ band tissue of *canoe* mutant embryos with or without (left) expressing Venus-Canoe-WT (middle) or Venus-Canoe-Δα-helix-1 (right). **(i)** Proposed model of mechanosensitive edge-to-vertex flow of Canoe/Afadin controlled by condensational states and Mbt/PAK activation. Boxes are 2^nd^ and 3^rd^ quartiles and whiskers are 5^th^ to 95^th^ percentiles; horizontal line is the median and “+” sign is the mean value. Mean±SD in (c) and (f); Mean±SEM between embryos in (h). ***P < 0.001; Not significant (ns) P > 0.1. Welch’s t-test for all comparisons. Live embryos in all panels. All positioned with anterior left, ventral down. Scale bars, 10 μm.

### Fluorescence Recovery After Photobleaching (FRAP)

FRAP experiments were performed on an Olympus FVMPE-RS Multiphoton Excitation Microscope with a UPlanSApo 100X1.35 oil-immersion objective. Embryos were mounted in a live imaging setup as described above, between a gas-permeable membrane and a coverslip. Bicellular junctions were identified by the fluorescent signal of endogenously Venus-tagged Canoe/Afadin. Time lapse movies were recorded first to determine the optimal laser intensity and frame rate to minimize photobleaching within the 5 minutes of data acquisition window. Next, after one pre-bleach image were taken, one bicellular junction was beached by the LSM stimulation module of Olympus FV30S-SW software, under laser power setting at 69%, duration setting at 201 milliseconds and scanning speed at 8 microsecond/pixel. A time-lapse movie was taken immediately after the bleaching for 5 minutes at 20 second per frame. One bleaching experiment was performed per imaging window, but multiple bleaching experiments were performed on the same embryo under different imaging windows. Maximum-intensity projections of 5-10 optical slides were generated for analysis using Fiji software.

### Immunofluorescence

Formaldehyde-fixed embryos were analyzed in Fig.1b, Extended Data Fig. 1b, 3a, 3f. All other images of this study were taken with live embryos. For immunofluorescence, dechorionated embryos were transferred into a glass jar with a mixture of 750mL deionized water, 750mL 19% Formaldehyde (Sigma) and 1.5mL Heptane (Sigma). Shaken for 15 minutes at 300 rpm and then vitelline membranes were manually removed using a glass needle. Primary antibodies used were mouse anti-alpha-tubulin (1:50, 12G10, Drosophila hybridoma bank (DSHB), supernatant and stock concentration unspecified), rabbit anti-GFP (2 μg/mL, TP401, Torry Pines), rat anti-Ecadherin (1:50, DCAD2, DSHB, supernatant and stock concentration unspecified). AlexaFluor-conjugated secondary antibodies (−488nm, -546nm or -647nm, Invitrogen) were used at 10 μg/mL. Filamentous actin was visualized by AlexaFluor-conjugated phalloidin (647nm, dissolved with 1ml DMSO, used at 1:100). Embryos were mounted in Prolong Gold (Invitrogen) and acquired on a Zeiss LSM980 confocal microscope with a PlanNeo 63×/1.3-NA oil-immersion objective (1 μm optical section and 0.5 μm z-steps) and a maximum-intensity projection of 3-4 optical slices was generated for analysis.

### Immunoprecipitation and immunoblotting

All biochemical analysis were preformed using *Drosophila* embryos. For immunoprecipitation, embryos were collected for 6 hours at 25°C and lysed in RIPA buffer with protease and phosphatase inhibitors. [50 mM Tris-HCl (pH 7.5), 125 mM NaCl, 5% glycerol, 0.5% NP-40, 1 mM phenylmethylsulphonyl fluoride, 2 mM sodium orthovanadate, protease inhibitor cocktail and phosphatase inhibitor cocktail (Epizyme). Lysates were homogenized by passing through a 1mL syringe with 0.6 X 32 mm needle and rotate at 4°C for 20 minutes before centrifuge for 30 minutes at 4°C. Supernatants were transferred to a new Eppendorf tube and quantified using a Pierce BCA protein assay kit (Thermo Fisher) and NanoDrop (Thermo Fisher). For input immunoblotting, 100μg total protein was diluted with lysis buffer to 80 μL and 20 μL 10XSDS sample buffer was added and boiled at 100°C for 6 minutes before gel loading. For immunoprecipitation, 20μL GFP-Trap magnetic beads was used for 1mg total protein in a total volume of 1mL. Lysate/Beads mixture was incubated at 4°C for 2 hrs. and then washed three times in lysis buffer using a magnetic stand (Epizyme). After final wash, magnetic beads were incubated with 2XSDS sample buffer and analyzed by Bis-Tris 4-10% pre-cast protein gels (Epizyme) and immunoblotting. Primary antibodies used were mouse anti-GFP (1 μg/mL, Roche), mouse anti-alpha-tubulin (12G10, DSHB, 1:100). Secondary antibodies used were IgG (H&L)-HRP Conjugated Goat anti-mouse and Goat anti-Rabbit (Easybio). Blots were visualized by ECL western blotting substrate (Pierce) and imaged by VILBER Fusion SOLO-S chemiluminescence imaging system.

### Quantification and statistical analysis

The workflow and calculation process for quantitative metrics repeatedly used in this study were described in a quantification manual (Table S3). All the N (number of measurements) and p values of group comparisons for each plot were compiled into Table S4 with references to each figure. All image processing and quantification were performed using Fiji software (2.1.0) and all plots were generated using GraphPad Prism 9. Quantitative metrics used were TCJ ratio for measure protein enrichment at tricellular junctions (Fig. 1g, 1i, 2c, 2l, 4d, 4k, 6f, Extended Data Fig. 7d); normalized edge intensity for measuring protein levels at bicellular junctions (Fig. 1h, 1j, 2d, 2k, Extended Data Fig. 2e, 3e), normalized TCJ intensity for measuring protein levels at tricellular junctions (Fig. 2e, 2f, 6g, Extended Data Fig. 3e, 6h); Junction cytoplasm ratio to measure protein enrichment at cell junctions versus the cytoplasm (Fig. 4c, 4f, Extended Data Fig. 5f-h, 6b, 6d); spatial co-localization and correlation analysis between two proteins (Fig.2e, 2f, 6g, Extended Data Fig. 6g); correlation coefficient for analyzing temporal correlation between two proteins (Fig. 2g); epithelial gap to measure adhesion defects between cells in embryos (Fig.3d, 3f, 4i, 4l, 6h); Intensity fold change to measure temporal dynamics of protein levels (Fig.5d); Junctional Equal Distribution Index (JEDI) to measure protein distribution among all edges within a single polygonal cell (Fig.5b, 5h-l, 6c, Extended Data Fig. 6e, 8c, 9c). In box and whiskers plots, boxes show the 25-75^th^ percentile, whiskers show the 5-95^th^ percentile, a horizontal line shows the median and a plus sign shows the mean. In column plots, heights show mean values and upper and lower whiskers show standard error values. Statistical analyses were performed using unpaired Student’s t-test with Welch’s correction without assuming equal standard deviations.

## Acknowledgements

We thank generous gifts of *Drosophila* lines and reagents from the Zallen Lab at Sloan Kettering Institute and support for initial proteomic screens of tricellular junction components. Zilong Wen, Ruoxi Wang, Yan Yan, Mingjie Zhang, Yan Zhao, Zhiyi Wei, Eric Brooks and Adam Pare for comments on the manuscript and helpful discussions; Bloomington Drosophila Stock Center, Tsinghua Fly Center, Kyoto Stock Center for fly strains; Genetivision for fly transgenic services; SUSTech University’s shared Imaging Core for technical support, confocal and spinning disk microscope usage; and Mark Peifer for his pioneering theories on Canoe/Afadin IDR domains. This works was supported by the National Natural Science Foundation of China (32270809 to H.H.Y) and seed funding support from School of Life Sciences, SUSTech.

## Author contributions

H.H.Y conceived of the project, designed and performed experiments, and wrote the paper. Q.Zheng, C.Zhang, M. Wang designed and performed experiments and analyzed data, Q.Zheng generated part of transgenic flies, designed and generated *mbt* CRISPR mutants. X. Xiang performed initial shRNA screens and noted preliminary abnormalities of Mbt/PAK4 knockdown embryos. S.Zhang performed all immunoprecipitation and immunoblot analysis. Y. Wang performed immunofluorescence experiments and analyzed data. All authors participated in interpreting results and discussions at all stages of the manuscript preparation.

## Captions for Supplementary Video 1 to 13

**Supplementary Video 1. Live imaging of Canoe-Venus and mCherry-Tubulin**. Dual-color time-lapse movie of mCherry-tubulin (middle, magenta) and endogenous Canoe-Venus (Left, green) during mitotic cycles in wild type embryos starting from stage 7, with merged image shown on the right. Note the synchronized cell division at mitotic domains at the upper half of images. Bar 10μm.

**Supplementary Video 2. Live imaging of Rap1-GTP sensor and mCherry-MRLC**. Dual-color time-lapse movie of mCherry-MRLC (middle, magenta) and ectopically expressed Venus-RALGDS-RBD (Left, green) in during mitotic cycles in wild type embryos starting from stage 7, with merged image shown on the right. Bar 10μm.

**Supplementary Video 3. Live imaging of Canoe-Venus and mCherry-Mbt/PAK**. Dual-color time-lapse movie of ectopically expressed mCherry-Mbt/PAK-WT (middle, magenta) and endogenous Canoe-Venus (Left, green) in during mitotic cycles in in wild type embryos at stage 7, with merged image shown on the right. Bar 10μm.

**Supplementary Video 4. Live imaging of Canoe-Venus and mCherry-MRLC in wild type and mbt knockdown embryos**. Dual-color time-lapse movie of mCherry-MRLC (middle, magenta) and endogenous Canoe-Venus (Left, green) in wild type (upper panel) and *mbt* knockdown (lower panel) embryos at stage 7, with merged image shown on the right. Note local condensation of Canoe-Venus at bicellular junctions and split of myosin cables in *mbt* knockdown embryos. Bar 10μm.

**Supplementary Video 5. Live imaging of MRLC-GFP and GAP43-mCherry**. Dual-color time-lapse movie of merged mCherry-GAP43 (magenta) and MRLC-GFP (green) during mitotic cycles in wild type embryos starting from stage 6. Note the ventral furrow invagination followed by synchronous cell division of mesectoderm cells, and the eventual sealing of ventral midline. Bar 10μm.

**Supplementary Video 6. Live imaging of Canoe-Venus and mCherry-MRLC in wild type and *mbt* knockdown embryos during ventral midline division and sealing.** Dual-color time-lapse movie of merged mCherry-MRLC (magenta) and endogenous Canoe-Venus (green) during synchronous cell division of mesectoderm cells and ventral midline sealing in wild type (left) and *mbt* knockdown embryos starting from stage 7. Note the apparent gaps along the ventral midline and failed sealing after division in *mbt* knockdown embryos. Bar 10μm.

**Supplementary Video 7. Live imaging of Canoe-Venus and mCherry-MRLC in wild type and *mbt* mutant embryos after *string* knockdown during ventral midline sealing.** Dual-color time-lapse movie of merged mCherry-MRLC (magenta) and endogenous Canoe-Venus (green) during ventral midline sealing without cell division after *string* knockdown in wild type (left) and *mbt* mutant embryos (right) starting from stage 7. Note that mesectoderm cells along the ventral midline failed to divide and ventral midline achieved normal sealing in both wild type and *mbt* mutant embryos. Bar 10μm.

**Supplementary Video 8. Live imaging of Spider-GFP in wild type and String/Cdc25 over-expressed embryos.** Time lapse movie of Spider-GFP (membrane marker, grey) in wild type and String/Cdc25 over-expressed embryos starting from stage 6 to stage 8. Whereas cell division only occurs within mitotic domains (ventral midline) in wild type embryos, all cells divided once in a nearly synchronized manner starting at stage 7 after ventral furrow invagination. Bar 10μm.

**Supplementary Video 9. Live imaging of MRLC-mCherry when String/Cdc25 is over-expressed in wild type and *mbt* mutant embryos.** Time lapse movie of MRLC-mCherry (grey) in String/Cdc25 over-expressed wild type and *mbt* mutant embryos starting from stage 7. In both wild type and *mbt* mutant embryos, all cells divided once in a nearly synchronized manner. In wild type embryos, cell quickly re-establish close contact after division, exemplified by the appearance of single-lined myosin cables. In contrast, cells failed to re-establish adhesion efficiently after division in *mbt* mutant embryos, visualized by the split of myosin into parallel cables. Bar 10μm.

**Supplementary Video 10. Live imaging of Canoe-Venus in post-mitotic wild type and *mbt* knockdown embryos.** Time-lapse movie endogenous Canoe-Venus in post-mitotic wild type (upper panel) and *mbt* knockdown (lower panel) embryos starting from stage 10. Note the extensive tactoid-shaped condensation of Canoe-Venus at bicellular junctions across the entire imaging region of *mbt* knockdown embryos. Bar 10μm.

**Supplementary Video 11. Live imaging of Canoe-Venus in post-mitotic *mbt* knockdown embryos before and after temperature-shift.** Time-lapse movie endogenous Canoe-Venus in a post-mitotic *mbt* knockdown embryo starting from stage 10. The embryo was imaged at 22°C for five time points and the incubated at 36°C for 10 minutes, then imaged again at 22°C for another 20 minutes. Note the extensive tactoid-shaped condensation of Canoe-Venus markedly disappeared after 36°C incubation and then gradually recovered after shift back to 22°C. Bar 10μm.

**Supplementary Video 12. Live imaging of Venus-Canoe-alpha-helix-1 and mCherry-MRLC**. Dual-color time-lapse movie of MRLC-tubulin (middle, magenta) and ectopically expressed Venus-tagged alpha-helix-1 of Canoe/Afadin (Left, green) in wild type embryos at stage 7, with merged image shown on the right. Note the extensive puncta formation of Canoe-alpha-helix-1 in the cytoplasm across the entire imaging region of wild type embryos. Bar 10μm.

**Supplementary Video 13. Live imaging of canoe mutant embryos rescued with either Canoe/Afadin-WT or Canoe/Afadin^Δalpha-helix-^**^1^. Dual-color time-lapse movie of MRLC-mCherry (magenta) and ectopically expressed Venus-tagged Canoe/Afadin transgenes (green) in *canoe* maternal mutant embryos at stage 7, with merged image shown. Note the extensive split of myosin cables and therefore epithelial gaps in *canoe* maternal mutant embryo rescued with Canoe/Afadin^Δalpha-helix-1^. Bar 10μm.

---

**Extended Data Fig.1:**
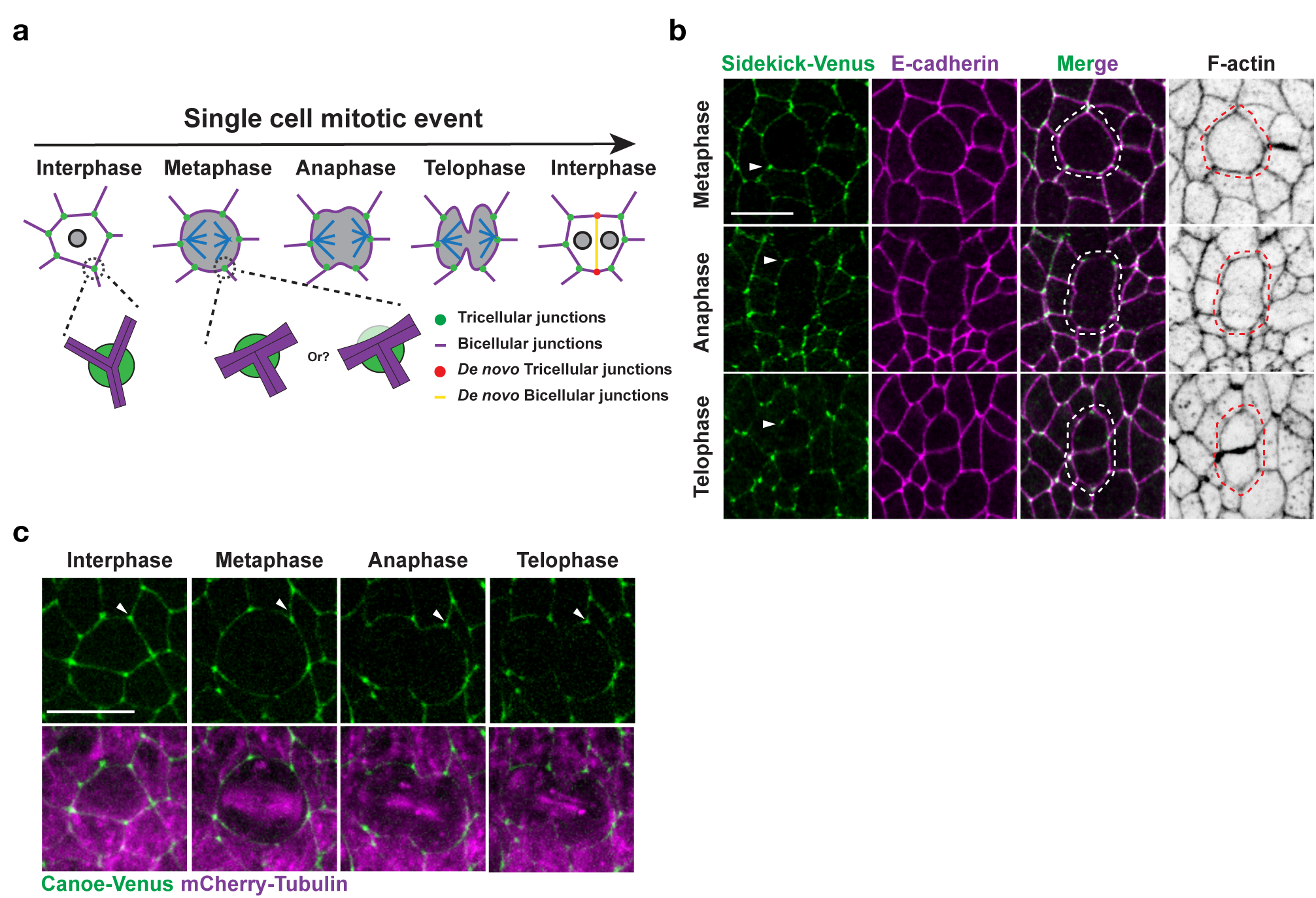
Bicellular and tricellular junction dynamics during single-cell mitotic events. **(a)** Schematic of tricellular junction remodeling during single-cell mitotic events. Each division generates one bicellular junction and two tricellular junctions *de novo*. Note the resolution limit illustrated by the question mark. **(b)** Localization of tricellular and bicellular junction transmembrane receptor Sidekick and E-cadherin by immunofluorescence, at different mitotic phases, characterized by stereotypical cell shapes changes visualized by F-actin. **(c)** Still images of a dual-color time lapse movie recording Canoe-Venus and mCherry-Tubulin during a single cell mitotic event. Arrowheads, Canoe-Venus of the same tricellular junction at different mitotic phases. Fixed embryos in (b), live embryos in (c). All positioned with anterior left, ventral down. Scale bars, 10 μm.

**Extended Data Fig.2:**
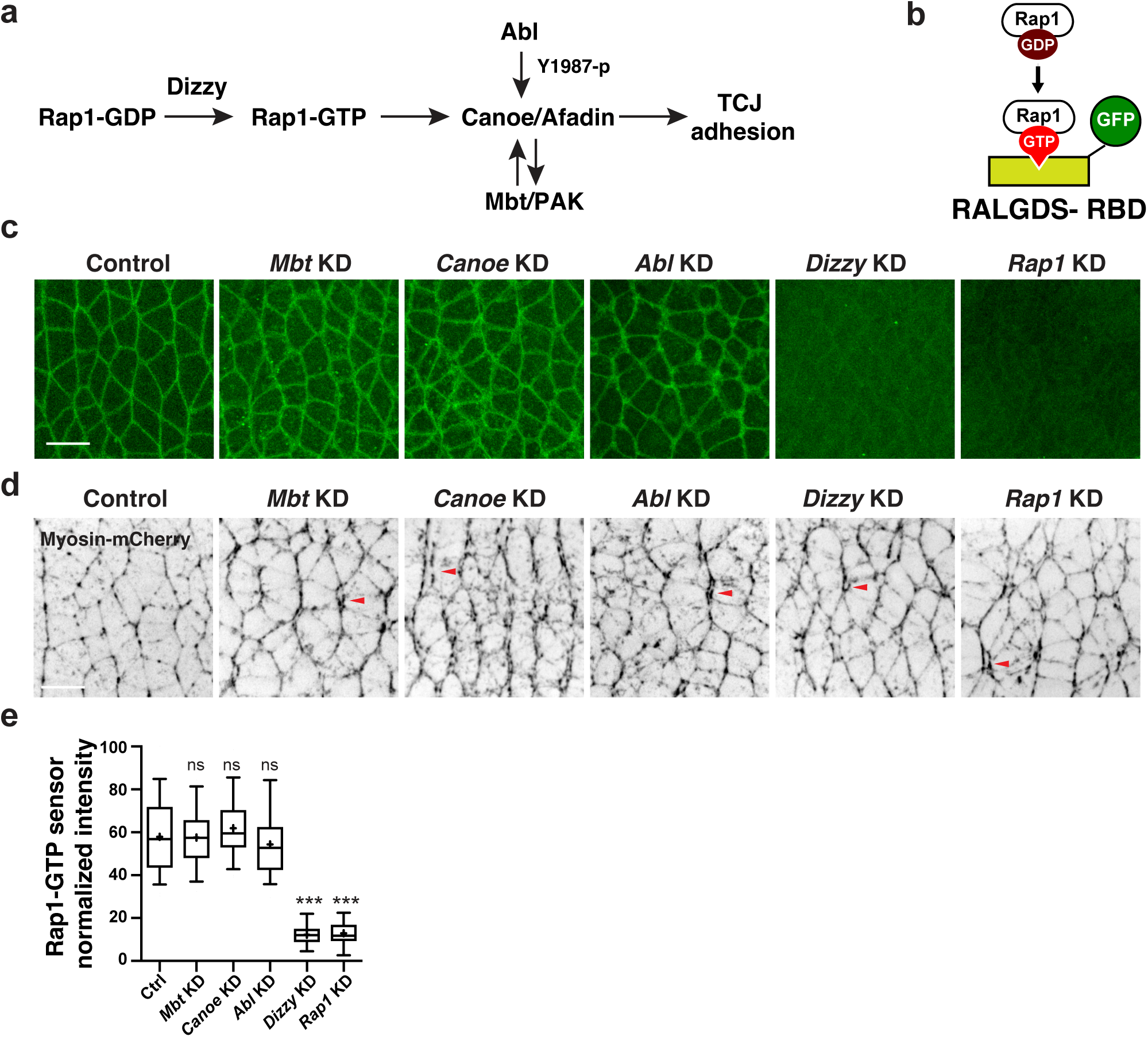
Development and validation of Rap1 activity sensor in live embryos. **(a)** Regulatory pathways of Canoe/Afadin localization and tricellular adhesion. **(b)** Schematic of Rap1-GTP sensor design. **(c, d)** Localization of Venus-Ralgds-RBD (c) and Myosin-mCherry (d) in control and embryos expressing shRNAs against Mbt, Canoe, Abl, Dizzy (Rap1-GEF) or Rap1 using the GAL4/UAS system. **(e)** Quantification of junctional intensity of Rap1-sensor in control or embryos with downregulated Mbt, Canoe, Abl, Dizzy (Rap1-GEF) or Rap1. All statistical comparisons were performed against control embryos. Boxes are 2^nd^ and 3^rd^ quartiles and whiskers are 5^th^ to 95^th^ percentiles; horizontal line is the median and “+” sign is the mean value. Mean±SD in (D), ***P < 0.001; Not significant (ns) P > 0.1. Welch’s t-test for all comparisons. Live embryos in all panels. All positioned with anterior left, ventral down. Scale bars, 10 μm.

**Extended Data Fig.3:**
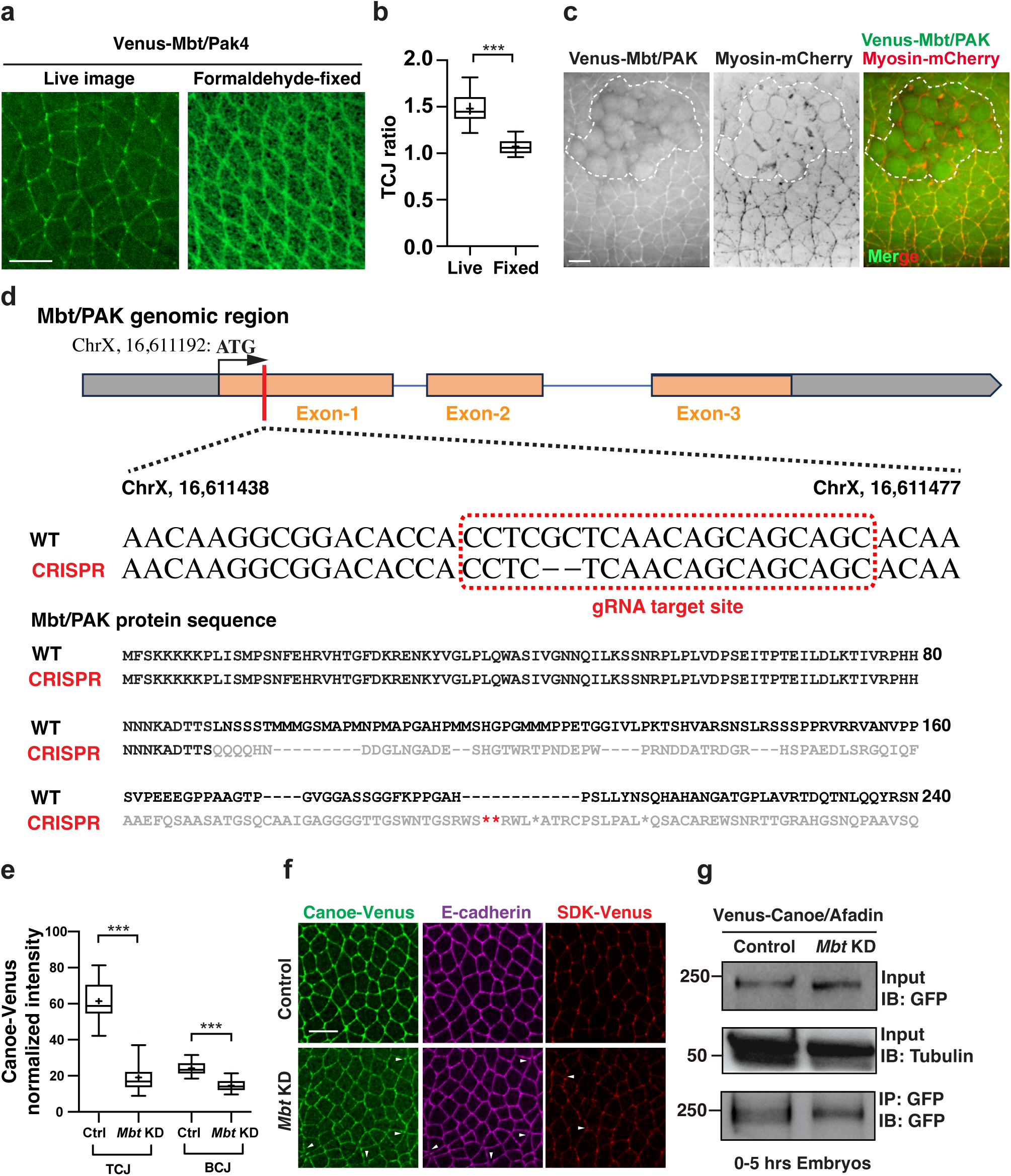
Mbt/PAK regulates Canoe/Afadin tricellular junction enrichment without affecting protein stability or general adherens junction integrity. **(a, b)** Localization and TCJ ratio of Venus-tagged Mbt/PAK in live (left) or fixed (right) embryos. **(c)** Still images of a dual-color time lapse movie recording Venus-Mbt/PAK and Myosin-mCherry during synchronous cell division of mitotic domains (dotted line) in live embryos. **(d)** Schematic of Mbt/PAK CRISPR design and sequencing results of *mbt* locus and alignment of translated protein before and after gRNA-mediated editing. **(e)** Quantification of tricellular and bicellular junction intensity of Canoe/Afadin in control and Mbt/PAK knockdown embryos. **(f)** Localization of tricellular and bicellular junction transmembrane receptor Sidekick and E-cadherin, with Canoe/Afadin by immunofluorescence in control and Mbt/PAK knockdown embryos. **(g)** Ectopically expressed Venus-Canoe-Afadin using the GAL4/UAS system was immunoprecipitated from 0-5 hrs wild type and Mbt/PAK knockdown embryos with an antibody to GFP and immunoblotted with antibodies to GFP (bottom). Input lysate was blotted with antibodies to GFP (top) and alpha-tubulin (middle). Boxes are 2^nd^ and 3^rd^ quartiles and whiskers are 5^th^ to 95^th^ percentiles; horizontal line is the median and “+” sign is the mean value. Mean±SD in (b), (e). ***P < 0.001. Welch’s t-test for all comparisons. Live embryos in (a, left) and (c), fixed embryos in (a, right) and (f). All positioned with anterior left, ventral down. Scale bars, 10 μm.

**Extended Data Fig.4:**
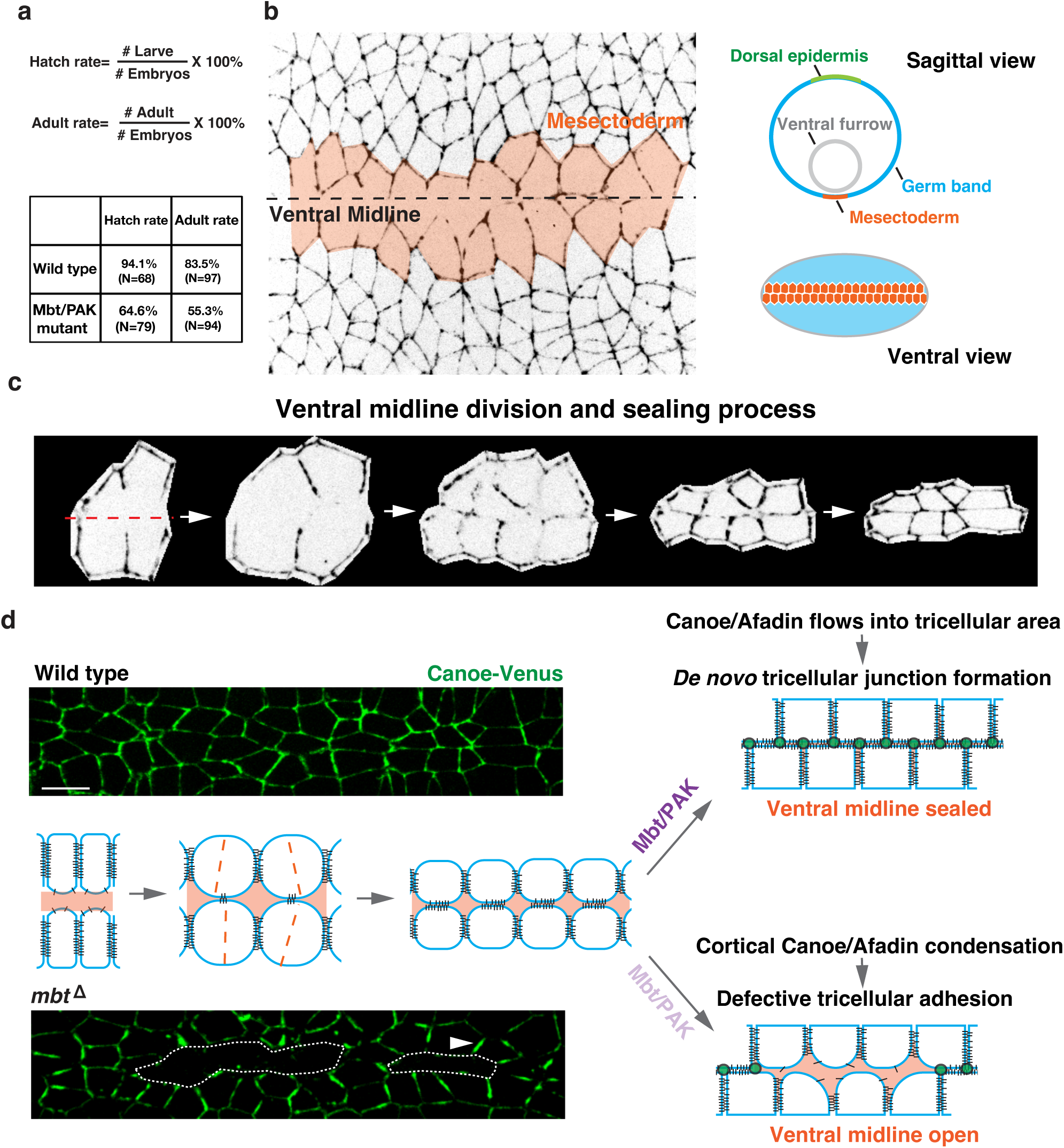
Physiological functions of the Mbt/PAK during ventral midline formation. **(a)** Hatch and adult rate of wild type and *mbt* mutant *Drosophila*, measuring fitness of embryogenesis and larvae-to-adult transition. **(b)** Positions of ventral midline, mesectoderm, ventral furrow, germ band tissues in pre-mitotic stage-7 embryos. **(c)** Dynamic remodeling of cell-cell interactions during synchronous cell division of mesectoderm cells along the ventral midline (dotted line, red). **(d)** Canoe-Venus localization in mesectoderm cells after synchronous divisions and illustration of adhesion defects progression in control (top) and *mbt* mutant embryos (bottom). Arrowheads, cortical condensation of Canoe-Venus after mitosis in *mbt* mutant embryos. Live embryos in all panels. All positioned with anterior left, ventral down. Scale bars, 10 μm.

**Extended Data Fig.5:**
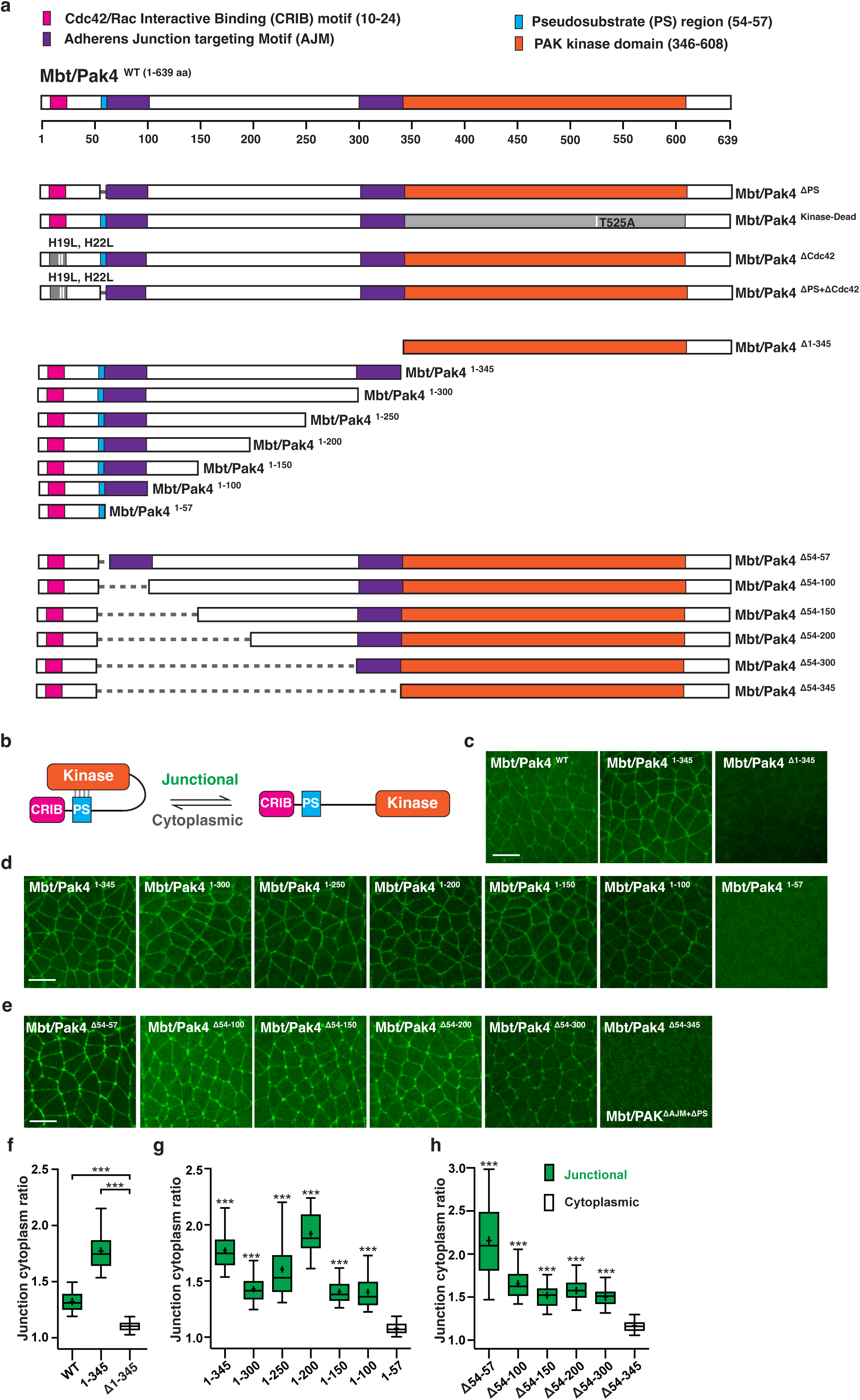
Identification of novel adherens junction localizing motifs (AJM) by systematic structural and functional analysis of Mbt/PAK. **(a)** Schematics of Mbt/PAK protein domains and variants generated. **(b)** Illustration of the auto-inhibition mechanism mediated by interactions between the Pseudo-Substrate motif and kinase domain. **(c, f)** Localization of wild type (full-length), N-terminal region (aa. 1-345), and C-terminal kinase domain (aa. 346-639) of Mbt/PAK in pre-mitotic embryos at stage 7 (b), quantified by junction-cytoplasm intensity ratios (f), with all statistical comparisons were performed against embryos expressing Venus-Mbt/PAKΔ1-345. **(d, g)** Localization of the N-terminal region of Mbt/PAK with decreasing length in pre-mitotic embryos (d), quantified by junction-cytoplasm intensity ratios (g), with all statistical comparisons were performed against embryos expressing Venus-Mbt/PAK1-57. **(e, h)** Localization of Mbt/PAK variants with or without the complete AJMs (e), quantified by junction-cytoplasm intensity ratios (h), with all statistical comparisons were performed against embryos expressing Venus-Mbt/PAK54-345. Boxes are 2nd and 3rd quartiles and whiskers are 5th to 95th percentiles; horizontal line is the median and “+” sign is the mean value. Mean±SD in all plots. ***P < 0.001. Welch’s t-test for all comparisons. Live embryos in all panels. All positioned with anterior left, ventral down. Scale bars, 10 μm.

**Extended Data Fig.6:**
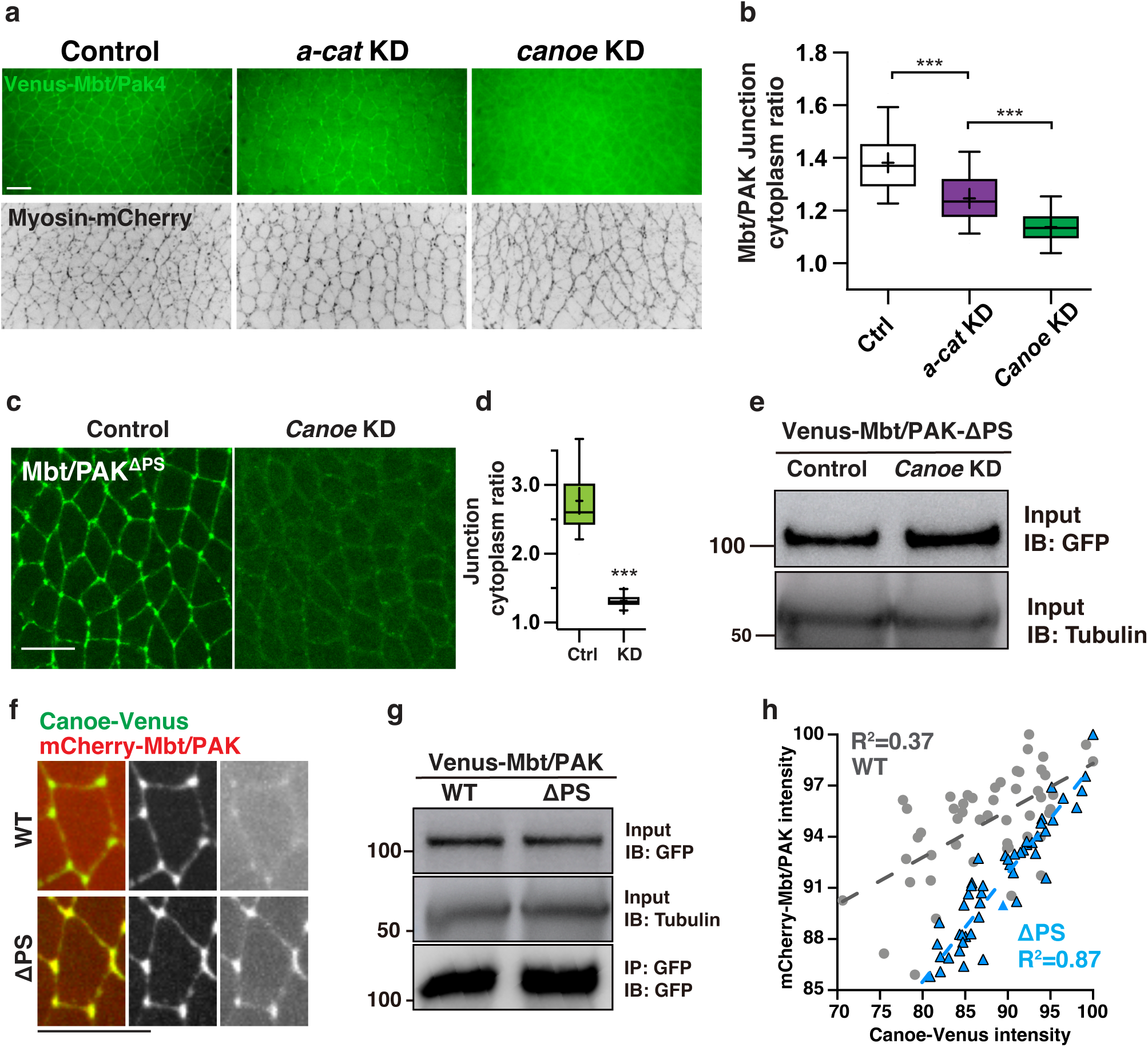
Mbt/PAK junctional localization is dependent on Canoe/Afadin. **(a, b)** Localization of Venus-Mbt/PAK (top) and Myosin-mCherry (bottom) in control and embryos expressing shRNAs against alpha-catenin or Canoe/Afadin using the GAL4/UAS system (a), quantified by junction-cytoplasm intensity ratios. Note the remaining Mbt/PAK junctional signal in alpha-catenin knockdown embryos, and complete absence of Mbt/PAK junctional signal in Canoe/Afadin knockdown embryos, despite of similar adhesion defects in both. **(c, d)** Localization of Venus-Mbt/PAK-ΔPS in control and *canoe* knockdown embryos (c), quantified by junction-cytoplasm ratio (d). **(e)** Ectopically expressed Venus-Mbt/ PAK-ΔPS in embryos using the GAL4/UAS system was lysed from 0-5 hrs control or *canoe* knockdown embryos and immunoblotted with antibodies to GFP and alpha-tubulin. **(f)** Still images of a dual-color time lapse movie recording Canoe/Afadin-Venus (green) together with ectopically expressed mCherry-Mbt/PAK-WT (top) or mCherry-Mbt/PAK-ΔPS (red) in embryos (bottom). **(g)** Ectopically expressed Venus-Mbt/PAK-WT and Venus-Mbt/PAK-ΔPS using the GAL4/UAS system was immunoprecipitated from 0-5 hrs wild type embryos with an antibody to GFP and immunoblotted with antibodies to GFP (bottom). Input lysate was blotted with antibodies to GFP (top) and alpha-tubulin (middle). **(h)** Spatial co-localization and correlation analysis between Canoe/Afadin and Mbt/PAK-WT (grey dots) or Mbt/PAK-ΔPS (blue triangles). Boxes are 2^nd^ and 3^rd^ quartiles and whiskers are 5^th^ to 95^th^ percentiles; horizontal line is the median and “+” sign is the mean value. Mean±SD in (b), (d), ***P<0.001. Welch’s t-test for all comparisons. Live stage 7 embryos in all panels. All positioned with anterior left, ventral down. Scale bars, 10 μm.

**Extended Data Fig.7:**
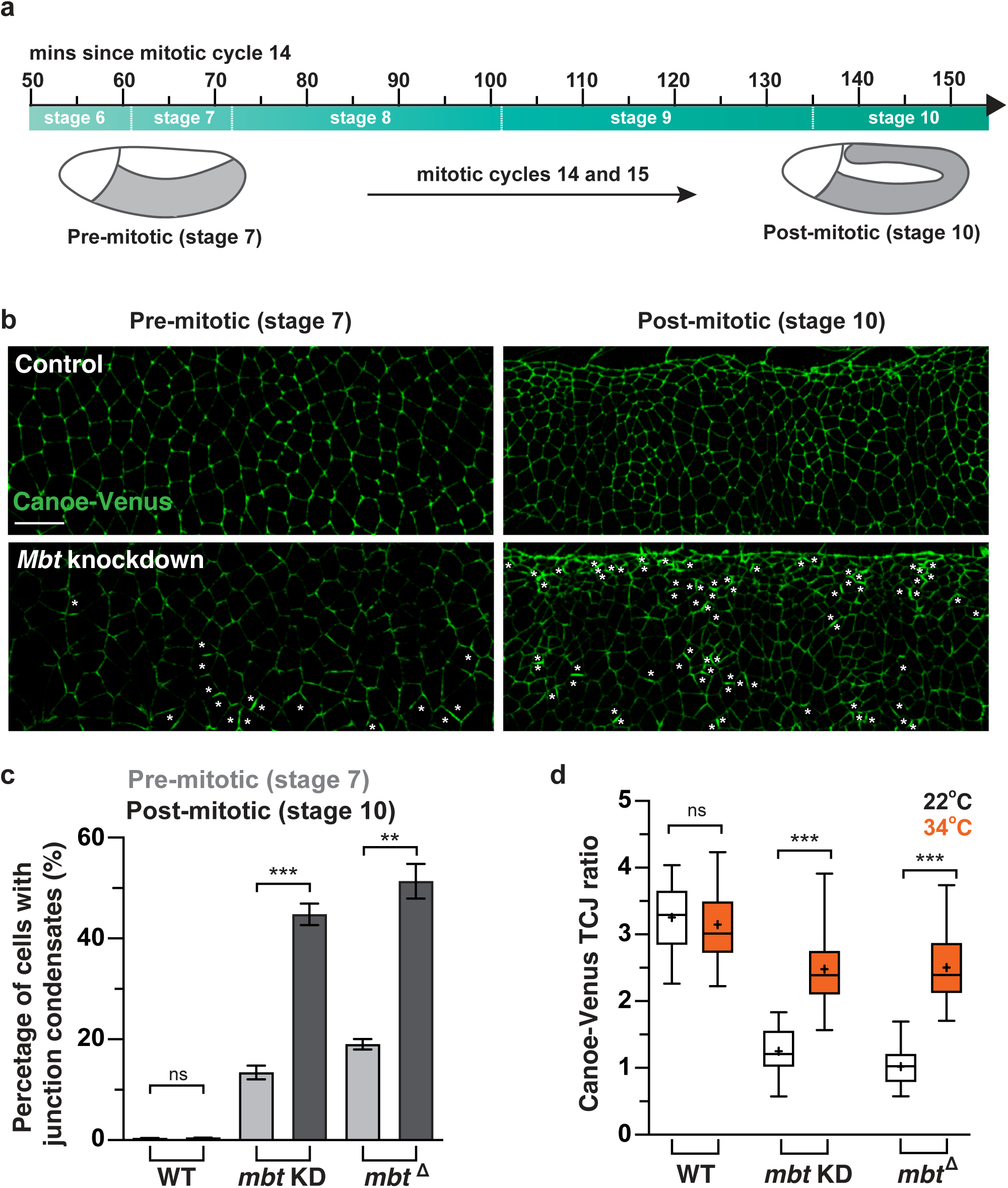
Increase of Canoe/Afadin cortical condensation after embryonic mitosis. **(a)** Time scales of embryogenesis from pre-mitotic stage 7 to post-mitotic stage 10 after two consecutive mitotic cycles (14 and 15). **(b)** Localization of Canoe-Venus in control and Mbt/PAK knockdown embryos before (stage 7) and after mitosis (stage 10). Asterisks marking cells with apparent Caneo/Afadin cortical condensation. **(c)** Percentage of cells with Canoe-Venus junction condensation in control, Mbt/PAK knockdown and *mbt* mutant embryos, before and after mitosis. Mean±SEM between embryos in (c). **(d)** TCJ ratios of Canoe-Venus in control, Mbt/PAK knockdown and *mbt* mutant embryos, before (white, 22°C) and after shifted to 34°C (orange). ***P < 0.001, **P < 0.01, Not significant (ns) P > 0.1. Welch’s t-test for all comparisons. Live embryos in all panels. All positioned with anterior left, ventral down. Scale bars, 10 μm.

**Extended Data Fig.8:**
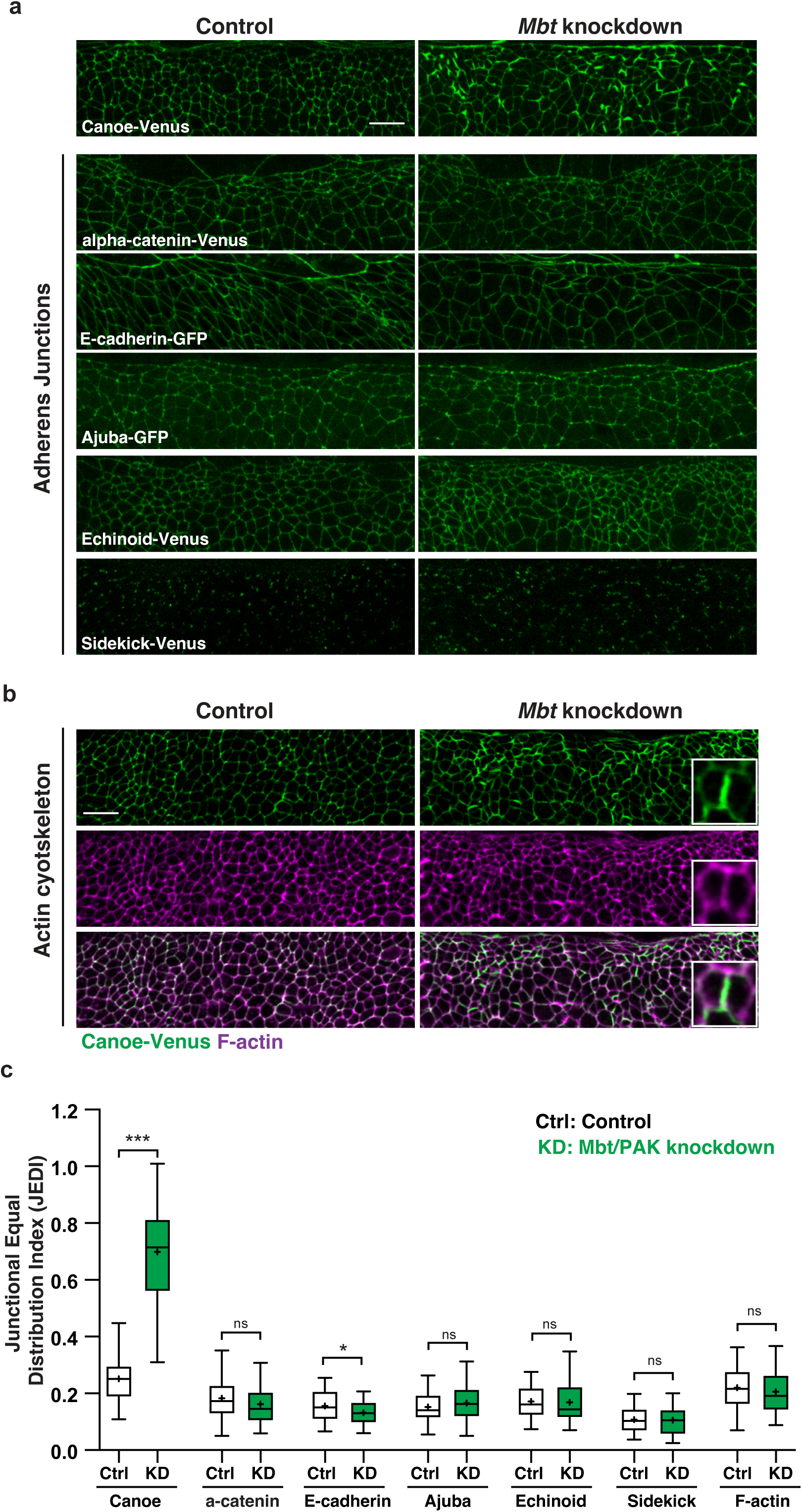
Canoe/Afadin forms cortical condensates despite normal localization of other adherens junction components and actin cytoskeleton organization. **(a)** Localization of canonical bicellular and tricellular adherens junction components in post-mitotic control and Mbt/PAK knockdown embryos, including Canoe, alpha-catenin, E-cadherin, Ajuba, Echinoid and Sidekick. **(b)** Localization of Canoe-Venus and F-actin by immunofluorescence in control and Mbt/PAK knockdown embryos. Insets, zoom-in images showing the apparent lack of condensation of F-actin. **(c)** JEDI values of canonical bicellular and tricellular adherens junction components in post-mitotic control and Mbt/PAK knockdown (green) embryos. Boxes are 2^nd^ and 3^rd^ quartiles and whiskers are 5^th^ to 95^th^ percentiles; horizontal line is the median and “+” sign is the mean value. Mean±SD in all plots. ***P < 0.001, *P < 0.1, Not significant (ns) P > 0.1. Welch’s t-test for all comparisons. Live embryos in (a) and fixed embryos in (b). All positioned with anterior left, ventral down. Scale bars, 10 μm.

**Extended Data Fig.9:**
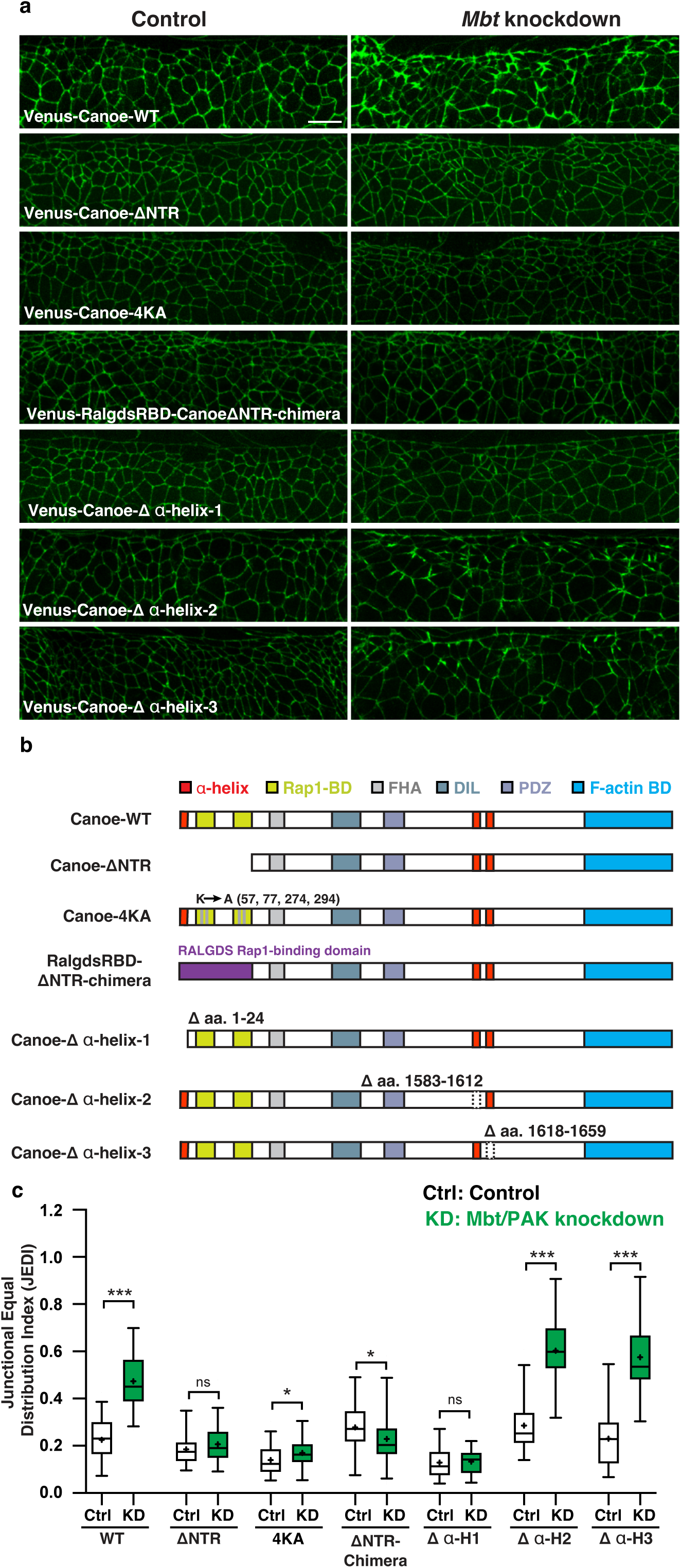
N-terminal alpha-helix and Rap1-binding are both necessary for Canoe/Afadin cortical condensation. **(a, b)** Localization of ectopically expressed Venus-Canoe-WT, Venus-Canoe-ΔNTR, Venus-Canoe-4KA, Venus-RalgdsRBD-CanoeΔNTR-chimera, Venus-Canoe-Δα-helix-1, Venus-Canoe-Δα-helix-2, and Venus-Canoe-Δα-helix-3 in control and *mbt* knockdown embryos at post-mitotic stage 10 (a), with domain organization of each Canoe/Afadin transgenes shown below (b). **(c)** JEDI values of ectopically expressed Venus-Canoe variants in post-mitotic control and Mbt/PAK knockdown (green) embryos. Boxes are 2^nd^ and 3^rd^ quartiles and whiskers are 5^th^ to 95^th^ percentiles; horizontal line is the median and “+” sign is the mean value. Mean±SD in all plots. ***P < 0.001, *P < 0.1, Not significant (ns) P > 0.1. Welch’s t-test for all comparisons. Live embryos in all panels. All positioned with anterior left, ventral down. Scale bars, 10 μm.

**Table S1.**
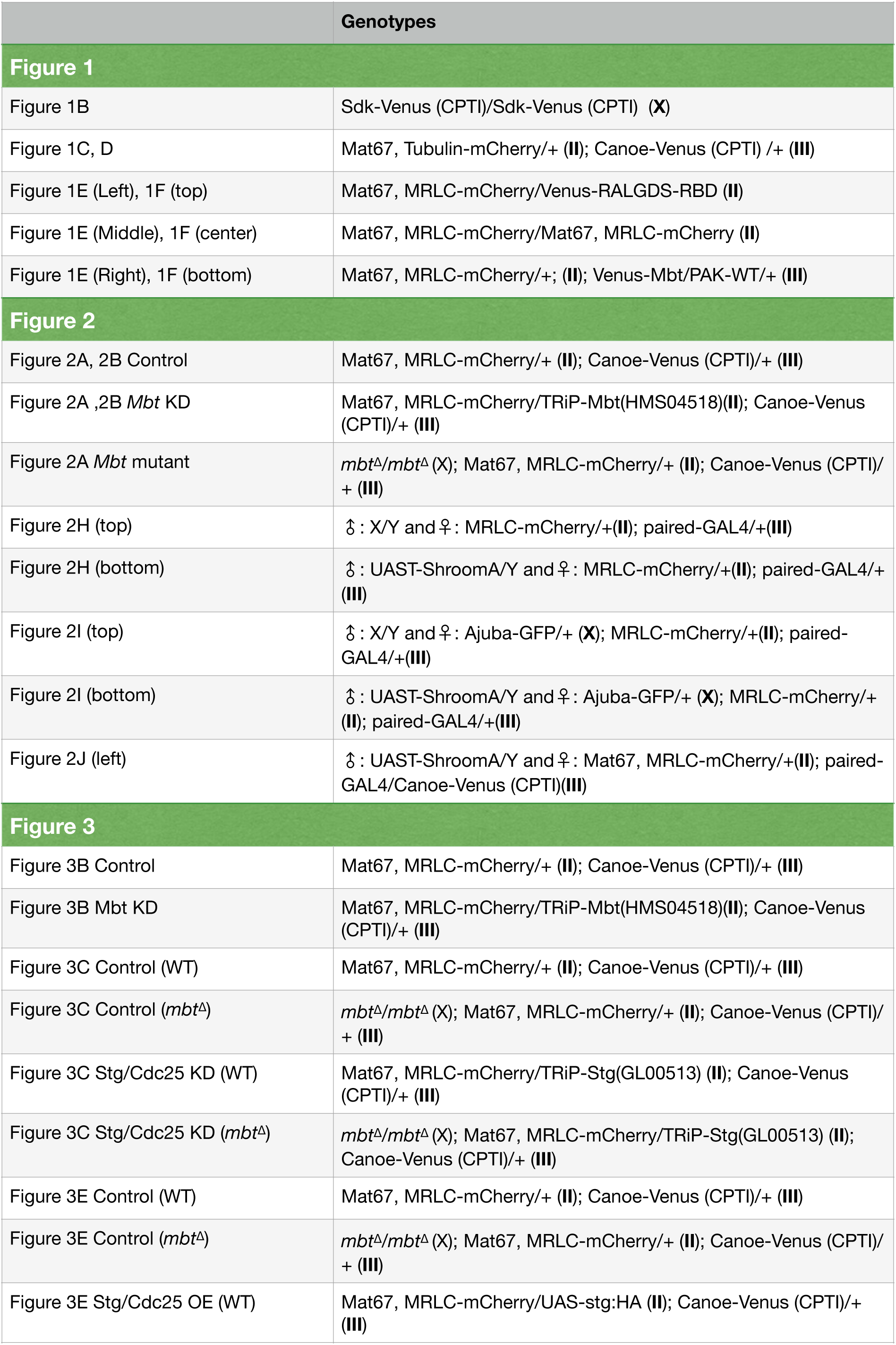
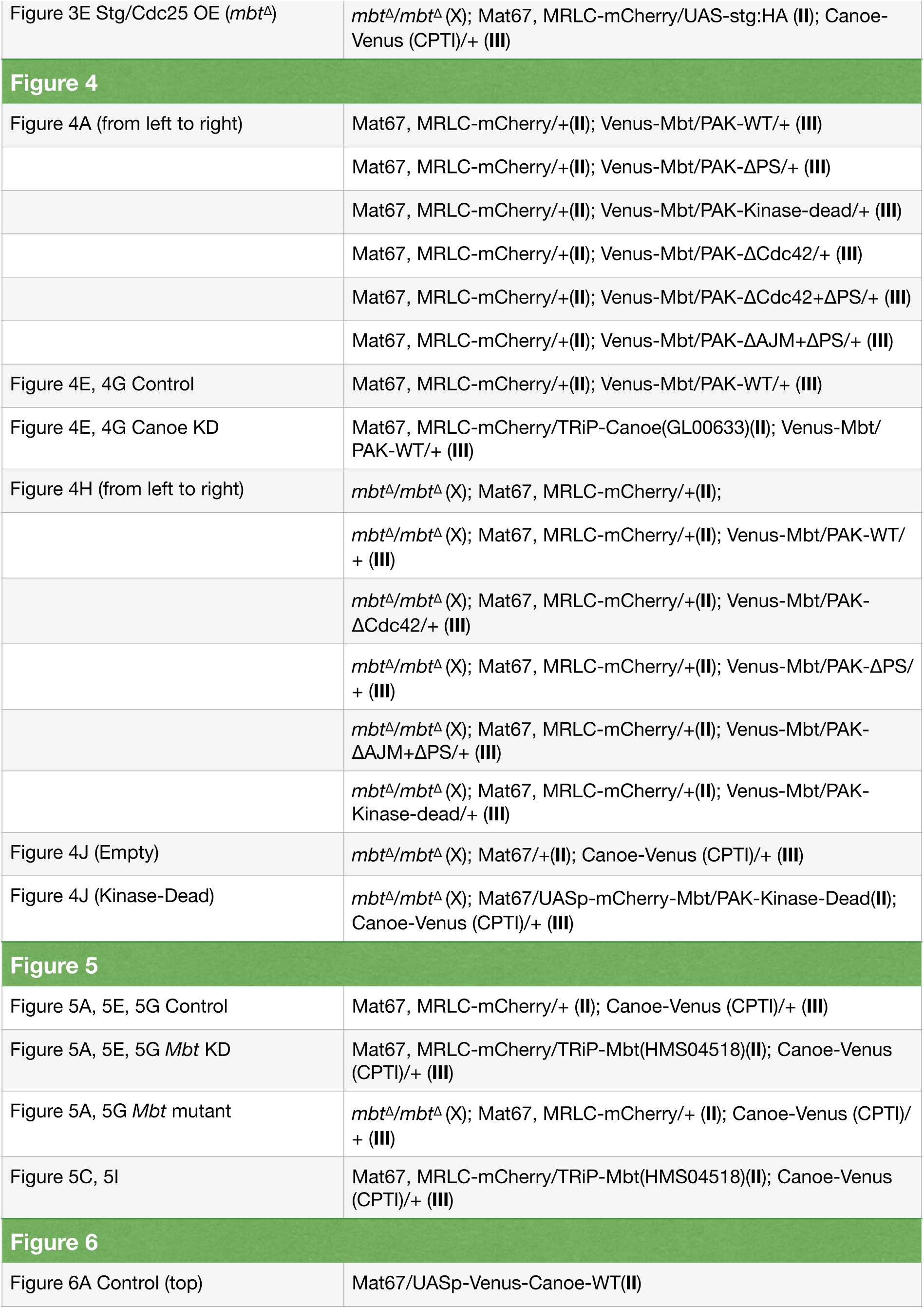
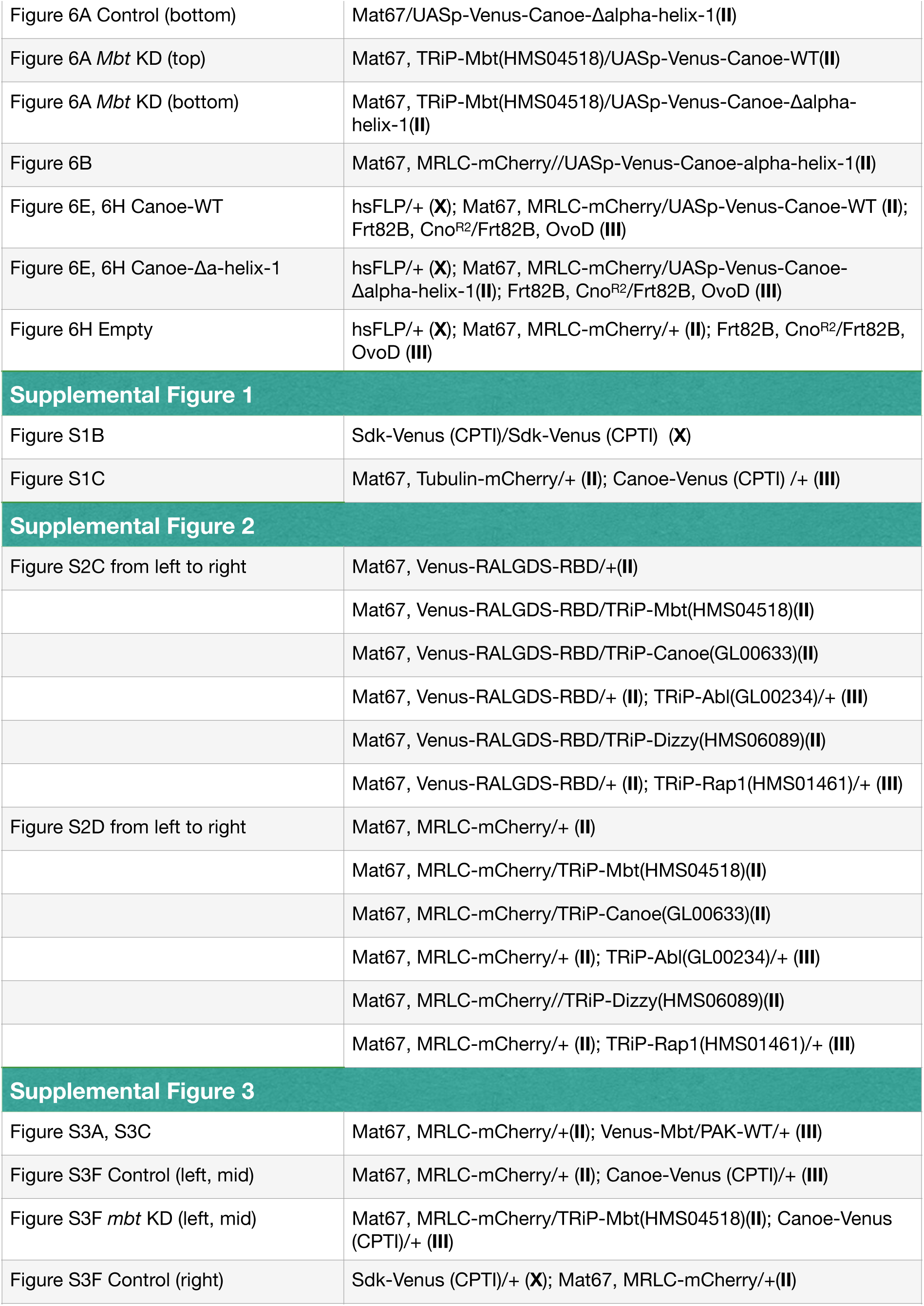
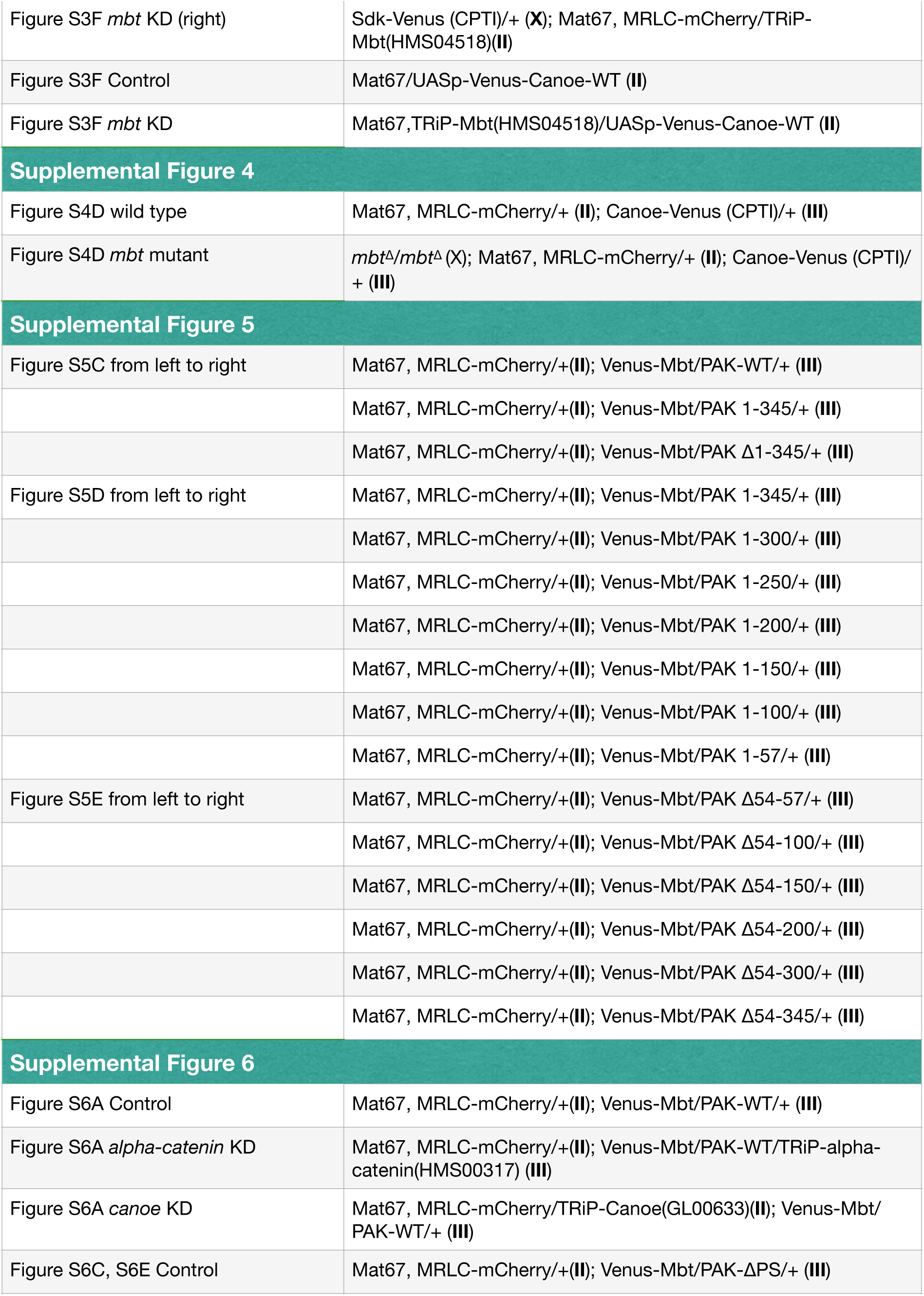
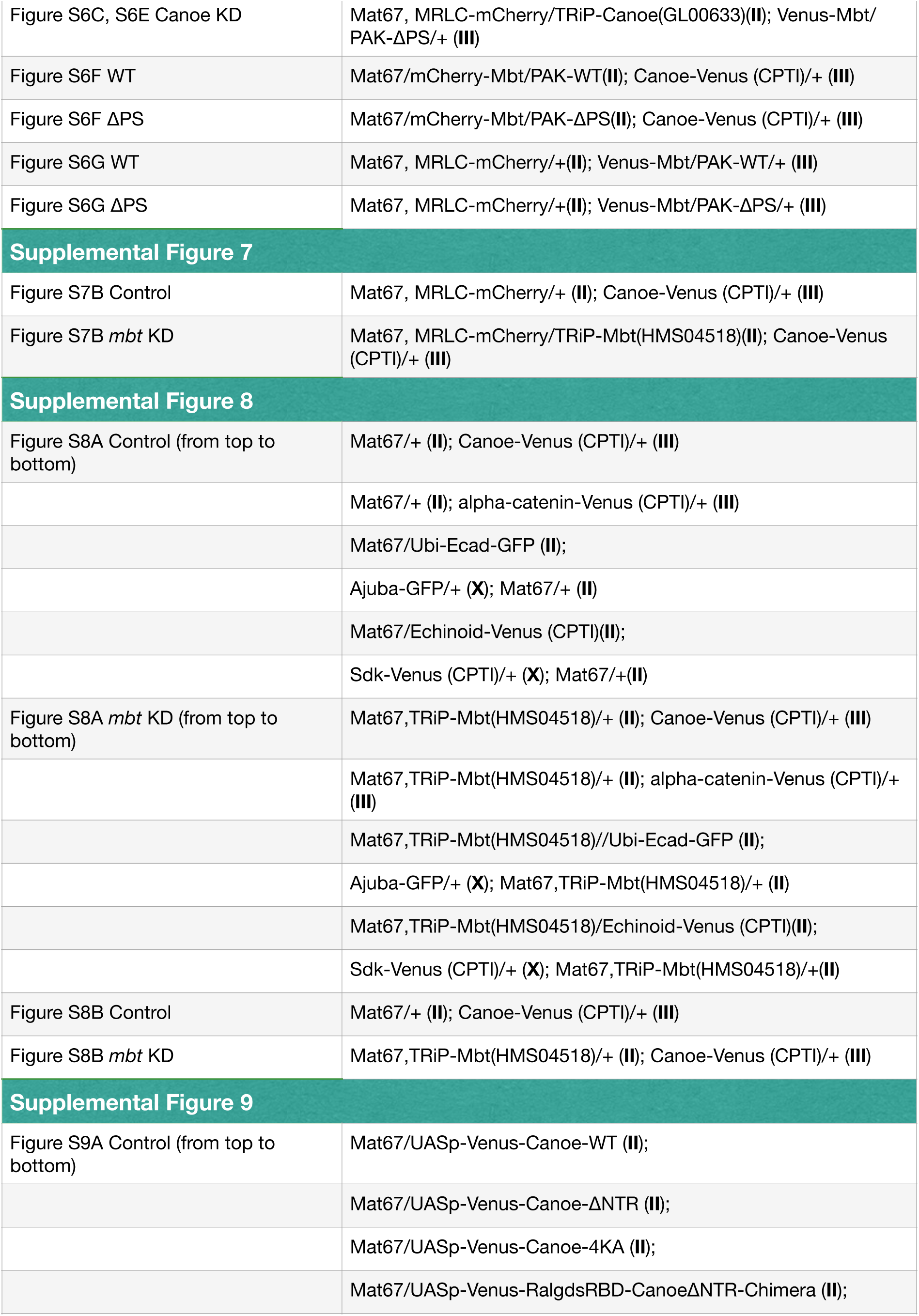
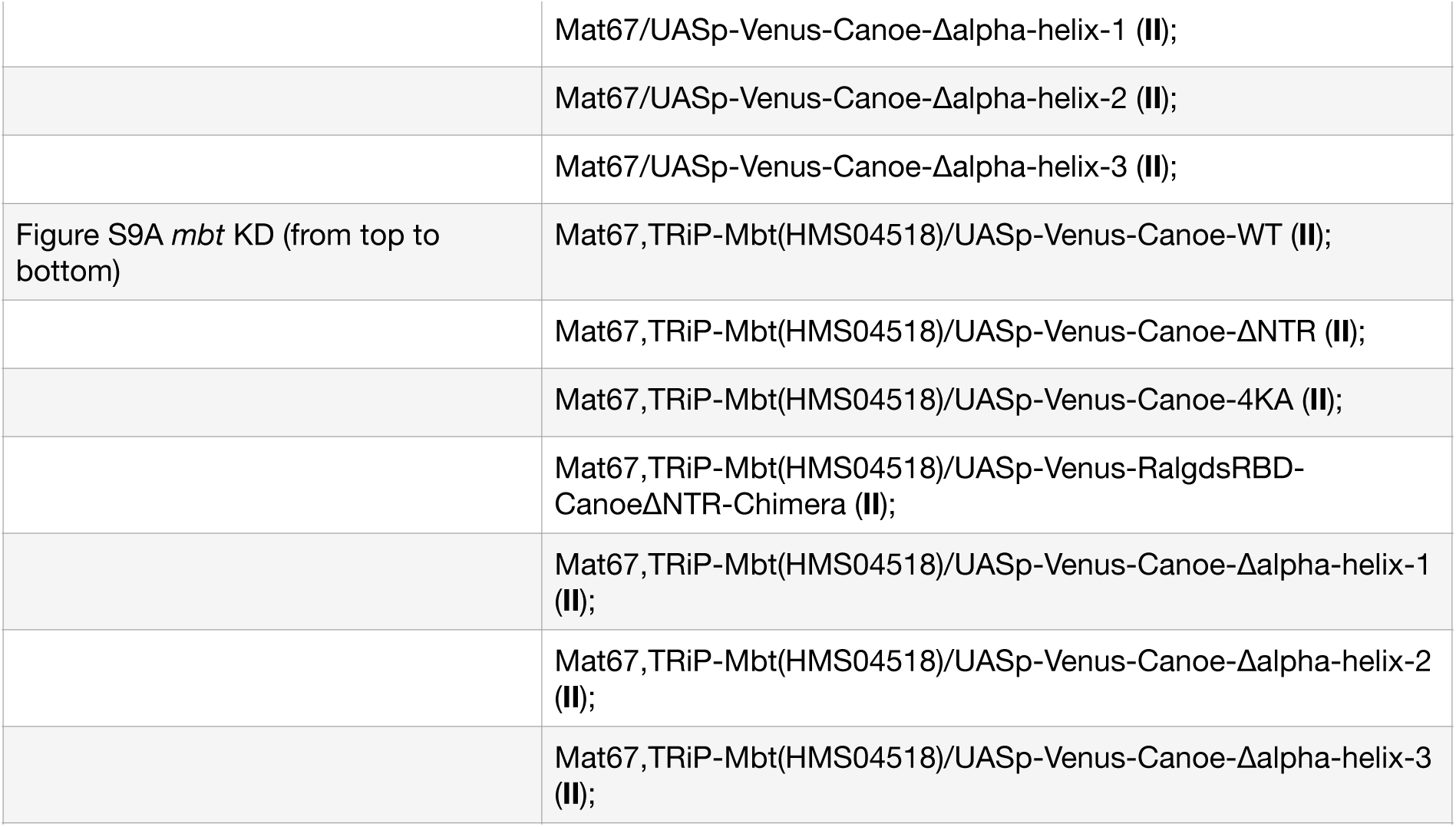

| Table S1 | Genotypes |
| --- | --- |
| **Figure 1** |  |
| Figure 1B | Sdk-Venus (CPTI)/Sdk-Venus (CPTI) (X) |
| Figure 1C, D | Mat67, Tubulin-mCherry/+ (II); Canoe-Venus (CPTI) /+ (III) |
| Figure 1E (Left), 1F (top) | Mat67, MRLC-mCherry/Venus-RALGDS-RBD (II) |
| Figure 1E (Middle), 1F (center) | Mat67, MRLC-mCherry/Mat67, MRLC-mCherry (II) |
| Figure 1E (Right), 1F (bottom) | Mat67, MRLC-mCherry/+; (II); Venus-Mbt/PAK-WT/+ (III) |
| **Figure 2** |  |
| Figure 2A, 2B Control | Mat67, MRLC-mCherry/+ (II); Canoe-Venus (CPTI)/+ (III) |
| Figure 2A ,2B *Mbt* KD | Mat67, MRLC-mCherry/TRiP-Mbt(HMS04518)(II); Canoe-Venus (CPTI)/+ (III) |
| Figure 2A *Mbt* mutant | *mbt<sup>Δ</sup>/mbt<sup>Δ</sup>* (X); Mat67, MRLC-mCherry/+ (II); Canoe-Venus (CPTI)/+ (III) |
| Figure 2H (top) | ♂ : X/Y and ♀ : MRLC-mCherry/+(II); paired-GAL4/+(III) |
| Figure 2H (bottom) | ♂ : UAST-ShroomA/Y and ♀ : MRLC-mCherry/+(II); paired-GAL4/+(III) |
| Figure 2I (top) | ♂ : X/Y and ♀ : Ajuba-GFP/+ (X); MRLC-mCherry/+(II); paired-GAL4/+(III) |
| Figure 2I (bottom) | ♂ : UAST-ShroomA/Y and ♀ : Ajuba-GFP/+ (X); MRLC-mCherry/+(II); paired-GAL4/+(III) |
| Figure 2J (left) | ♂ : UAST-ShroomA/Y and ♀ : Mat67, MRLC-mCherry/+(II); paired-GAL4/Canoe-Venus (CPTI)(III) |
| **Figure 3** |  |
| Figure 3B Control | Mat67, MRLC-mCherry/+ (II); Canoe-Venus (CPTI)/+ (III) |
| Figure 3B *Mbt* KD | Mat67, MRLC-mCherry/TRiP-Mbt(HMS04518)(II); Canoe-Venus (CPTI)/+ (III) |
| Figure 3C Control (WT) | Mat67, MRLC-mCherry/+ (II); Canoe-Venus (CPTI)/+ (III) |
| Figure 3C Control (*mbt<sup>Δ</sup>*) | *mbt<sup>Δ</sup>/mbt<sup>Δ</sup>* (X); Mat67, MRLC-mCherry/+ (II); Canoe-Venus (CPTI)/+ (III) |
| Figure 3C Stg/Cdc25 KD (WT) | Mat67, MRLC-mCherry/TRiP-Stg(GL00513) (II); Canoe-Venus (CPTI)/+ (III) |
| Figure 3C Stg/Cdc25 KD (*mbt<sup>Δ</sup>*) | *mbt<sup>Δ</sup>/mbt<sup>Δ</sup>* (X); Mat67, MRLC-mCherry/TRiP-Stg(GL00513) (II); Canoe-Venus (CPTI)/+ (III) |
| Figure 3E Control (WT) | Mat67, MRLC-mCherry/+ (II); Canoe-Venus (CPTI)/+ (III) |
| Figure 3E Control (*mbt<sup>Δ</sup>*) | *mbt<sup>Δ</sup>/mbt<sup>Δ</sup>* (X); Mat67, MRLC-mCherry/+ (II); Canoe-Venus (CPTI)/+ (III) |
| Figure 3E Stg/Cdc25 OE (WT) | Mat67, MRLC-mCherry/UAS-stg:HA (II); Canoe-Venus (CPTI)/+ (III) |
| Figure 3E Stg/Cdc25 OE (*mbt<sup>Δ</sup>*) | *mbt<sup>Δ</sup>/mbt<sup>Δ</sup>* (X); Mat67, MRLC-mCherry/UAS-stg:HA (II); Canoe-Venus (CPTI)/+ (III) |
| **Figure 4** |  |
| Figure 4A (from left to right) | Mat67, MRLC-mCherry/+ (II); Venus-Mbt/PAK-WT/+ (III) |
|  | Mat67, MRLC-mCherry/+ (II); Venus-Mbt/PAK-ΔPS/+ (III) |
|  | Mat67, MRLC-mCherry/+ (II); Venus-Mbt/PAK-Kinase-dead/+ (III) |
|  | Mat67, MRLC-mCherry/+ (II); Venus-Mbt/PAK-ΔCdc42/+ (III) |
|  | Mat67, MRLC-mCherry/+ (II); Venus-Mbt/PAK-ΔCdc42+ΔPS/+ (III) |
|  | Mat67, MRLC-mCherry/+ (II); Venus-Mbt/PAK-ΔAJM+ΔPS/+ (III) |
| Figure 4E, 4G Control | Mat67, MRLC-mCherry/+ (II); Venus-Mbt/PAK-WT/+ (III) |
| Figure 4E, 4G Canoe KD | Mat67, MRLC-mCherry/TRiP-Canoe(GL00633)(II); Venus-Mbt/PAK-WT/+ (III) |
| Figure 4H (from left to right) | *mbt<sup>Δ</sup>/mbt<sup>Δ</sup>* (X); Mat67, MRLC-mCherry/+ (II); |
|  | *mbt<sup>Δ</sup>/mbt<sup>Δ</sup>* (X); Mat67, MRLC-mCherry/+ (II); Venus-Mbt/PAK-WT/+ (III) |
|  | *mbt<sup>Δ</sup>/mbt<sup>Δ</sup>* (X); Mat67, MRLC-mCherry/+ (II); Venus-Mbt/PAK-ΔCdc42/+ (III) |
|  | *mbt<sup>Δ</sup>/mbt<sup>Δ</sup>* (X); Mat67, MRLC-mCherry/+ (II); Venus-Mbt/PAK-ΔPS/+ (III) |
|  | *mbt<sup>Δ</sup>/mbt<sup>Δ</sup>* (X); Mat67, MRLC-mCherry/+ (II); Venus-Mbt/PAK-ΔAJM+ΔPS/+ (III) |
|  | *mbt<sup>Δ</sup>/mbt<sup>Δ</sup>* (X); Mat67, MRLC-mCherry/+ (II); Venus-Mbt/PAK-Kinase-dead/+ (III) |
| Figure 4J (Empty) | *mbt<sup>Δ</sup>/mbt<sup>Δ</sup>* (X); Mat67/+ (II); Canoe-Venus (CPTI)/+ (III) |
| Figure 4J (Kinase-Dead) | *mbt<sup>Δ</sup>/mbt<sup>Δ</sup>* (X); Mat67/UASp-mCherry-Mbt/PAK-Kinase-Dead(II); Canoe-Venus (CPTI)/+ (III) |
| **Figure 5** |  |
| Figure 5A, 5E, 5G Control | Mat67, MRLC-mCherry/+ (II); Canoe-Venus (CPTI)/+ (III) |
| Figure 5A, 5E, 5G *Mbt* KD | Mat67, MRLC-mCherry/TRiP-Mbt(HMS04518)(II); Canoe-Venus (CPTI)/+ (III) |
| Figure 5A, 5G *Mbt* mutant | *mbt<sup>Δ</sup>/mbt<sup>Δ</sup>* (X); Mat67, MRLC-mCherry/+ (II); Canoe-Venus (CPTI)/+ (III) |
| Figure 5C, 5I | Mat67, MRLC-mCherry/TRiP-Mbt(HMS04518)(II); Canoe-Venus (CPTI)/+ (III) |
| **Figure 6** |  |
| Figure 6A Control (top) | Mat67/UASp-Venus-Canoe-WT(II) |
| Figure 6A Control (bottom) | Mat67/UASp-Venus-Canoe-Δalpha-helix-1(II) |
| Figure 6A *Mbt* KD (top) | Mat67, TRiP-Mbt(HMS04518)/UASp-Venus-Canoe-WT(II) |
| Figure 6A *Mbt* KD (bottom) | Mat67, TRiP-Mbt(HMS04518)/UASp-Venus-Canoe-Δalpha-helix-1(II) |
| Figure 6B | Mat67, MRLC-mCherry//UASp-Venus-Canoe-alpha-helix-1(II) |
| Figure 6E, 6H Canoe-WT | hsFLP/+ (X); Mat67, MRLC-mCherry/UASp-Venus-Canoe-WT (II); Frt82B, Cno<sup>R2</sup>/Frt82B, OvoD (III) |
| Figure 6E, 6H Canoe-Δa-helix-1 | hsFLP/+ (X); Mat67, MRLC-mCherry/UASp-Venus-Canoe-Δalpha-helix-1(II); Frt82B, Cno<sup>R2</sup>/Frt82B, OvoD (III) |
| Figure 6H Empty | hsFLP/+ (X); Mat67, MRLC-mCherry/+ (II); Frt82B, Cno<sup>R2</sup>/Frt82B, OvoD (III) |
| **Supplemental Figure 1** |  |
| Figure S1B | Sdk-Venus (CPTI)/Sdk-Venus (CPTI) (X) |
| Figure S1C | Mat67, Tubulin-mCherry/+ (II); Canoe-Venus (CPTI) /+ (III) |
| **Supplemental Figure 2** |  |
| Figure S2C from left to right | Mat67, Venus-RALGDS-RBD/+ (II) |
|  | Mat67, Venus-RALGDS-RBD/TRiP-Mbt(HMS04518)(II) |
|  | Mat67, Venus-RALGDS-RBD/TRiP-Canoe(GL00633)(II) |
|  | Mat67, Venus-RALGDS-RBD/+ (II); TRiP-Abl(GL00234)/+ (III) |
|  | Mat67, Venus-RALGDS-RBD/TRiP-Dizzy(HMS06089)(II) |
|  | Mat67, Venus-RALGDS-RBD/+ (II); TRiP-Rap1(HMS01461)/+ (III) |
| Figure S2D from left to right | Mat67, MRLC-mCherry/+ (II) |
|  | Mat67, MRLC-mCherry/TRiP-Mbt(HMS04518)(II) |
|  | Mat67, MRLC-mCherry/TRiP-Canoe(GL00633)(II) |
|  | Mat67, MRLC-mCherry/+ (II); TRiP-Abl(GL00234)/+ (III) |
|  | Mat67, MRLC-mCherry//TRiP-Dizzy(HMS06089)(II) |
|  | Mat67, MRLC-mCherry/+ (II); TRiP-Rap1(HMS01461)/+ (III) |
| **Supplemental Figure 3** |  |
| Figure S3A, S3C | Mat67, MRLC-mCherry/+ (II); Venus-Mbt/PAK-WT/+ (III) |
| Figure S3F Control (left, mid) | Mat67, MRLC-mCherry/+ (II); Canoe-Venus (CPTI)/+ (III) |
| Figure S3F *mbt* KD (left, mid) | Mat67, MRLC-mCherry/TRiP-Mbt(HMS04518)(II); Canoe-Venus (CPTI)/+ (III) |
| Figure S3F Control (right) | Sdk-Venus (CPTI)/+ (X); Mat67, MRLC-mCherry/+ (II) |
| Figure S3F *mbt* KD (right) | Sdk-Venus (CPTI)/+ (X); Mat67, MRLC-mCherry/TRiP-Mbt(HMS04518)(II) |
| Figure S3F Control | Mat67/UASp-Venus-Canoe-WT (II) |
| Figure S3F *mbt* KD | Mat67,TRiP-Mbt(HMS04518)/UASp-Venus-Canoe-WT (II) |
| **Supplemental Figure 4** |  |
| Figure S4D wild type | Mat67, MRLC-mCherry/+ (II); Canoe-Venus (CPTI)/+ (III) |
| Figure S4D *mbt* mutant | *mbt<sup>Δ</sup>/mbt<sup>Δ</sup>* (X); Mat67, MRLC-mCherry/+ (II); Canoe-Venus (CPTI)/+ (III) |
| **Supplemental Figure 5** |  |
| Figure S5C from left to right | Mat67, MRLC-mCherry/+(II); Venus-Mbt/PAK-WT/+ (III) |
|  | Mat67, MRLC-mCherry/+(II); Venus-Mbt/PAK 1-345/+ (III) |
|  | Mat67, MRLC-mCherry/+(II); Venus-Mbt/PAK Δ1-345/+ (III) |
| Figure S5D from left to right | Mat67, MRLC-mCherry/+(II); Venus-Mbt/PAK 1-345/+ (III) |
|  | Mat67, MRLC-mCherry/+(II); Venus-Mbt/PAK 1-300/+ (III) |
|  | Mat67, MRLC-mCherry/+(II); Venus-Mbt/PAK 1-250/+ (III) |
|  | Mat67, MRLC-mCherry/+(II); Venus-Mbt/PAK 1-200/+ (III) |
|  | Mat67, MRLC-mCherry/+(II); Venus-Mbt/PAK 1-150/+ (III) |
|  | Mat67, MRLC-mCherry/+(II); Venus-Mbt/PAK 1-100/+ (III) |
|  | Mat67, MRLC-mCherry/+(II); Venus-Mbt/PAK 1-57/+ (III) |
| Figure S5E from left to right | Mat67, MRLC-mCherry/+(II); Venus-Mbt/PAK Δ54-57/+ (III) |
|  | Mat67, MRLC-mCherry/+(II); Venus-Mbt/PAK Δ54-100/+ (III) |
|  | Mat67, MRLC-mCherry/+(II); Venus-Mbt/PAK Δ54-150/+ (III) |
|  | Mat67, MRLC-mCherry/+(II); Venus-Mbt/PAK Δ54-200/+ (III) |
|  | Mat67, MRLC-mCherry/+(II); Venus-Mbt/PAK Δ54-300/+ (III) |
|  | Mat67, MRLC-mCherry/+(II); Venus-Mbt/PAK Δ54-345/+ (III) |
| **Supplemental Figure 6** |  |
| Figure S6A Control | Mat67, MRLC-mCherry/+(II); Venus-Mbt/PAK-WT/+ (III) |
| Figure S6A *alpha-catenin* KD | Mat67, MRLC-mCherry/+(II); Venus-Mbt/PAK-WT/TRiP-*alpha-catenin*(HMS00317) (III) |
| Figure S6A *canoe* KD | Mat67, MRLC-mCherry/TRiP-Canoe(GL00633)(II); Venus-Mbt/PAK-WT/+ (III) |
| Figure S6C, S6E Control | Mat67, MRLC-mCherry/+(II); Venus-Mbt/PAK-ΔPS/+ (III) |

| **Table S1** | **Genotypes** |
| --- | --- |
| Figure S6C, S6E Canoe KD | Mat67, MRLC-mCherry/TRiP-Canoe(GL00633)(II); Venus-Mbt/PAK-ΔPS/+ (III) |
| Figure S6F WT | Mat67/mCherry-Mbt/PAK-WT(II); Canoe-Venus (CPTI)/+ (III) |
| Figure S6F ΔPS | Mat67/mCherry-Mbt/PAK-ΔPS(II); Canoe-Venus (CPTI)/+ (III) |
| Figure S6G WT | Mat67, MRLC-mCherry/+ (II); Venus-Mbt/PAK-WT/+ (III) |
| Figure S6G ΔPS | Mat67, MRLC-mCherry/+ (II); Venus-Mbt/PAK-ΔPS/+ (III) |
| **Supplemental Figure 7** |  |
| Figure S7B Control | Mat67, MRLC-mCherry/+ (II); Canoe-Venus (CPTI)/+ (III) |
| Figure S7B *mbt* KD | Mat67, MRLC-mCherry/TRiP-Mbt(HMS04518)(II); Canoe-Venus (CPTI)/+ (III) |
| **Supplemental Figure 8** |  |
| Figure S8A Control (from top to bottom) | Mat67/+ (II); Canoe-Venus (CPTI)/+ (III) |
|  | Mat67/+ (II); alpha-catenin-Venus (CPTI)/+ (III) |
|  | Mat67/Ubi-Ecad-GFP (II); |
|  | Ajuba-GFP/+ (X); Mat67/+ (II) |
|  | Mat67/Echinoid-Venus (CPTI)(II); |
|  | Sdk-Venus (CPTI)/+ (X); Mat67/+(II) |
| Figure S8A *mbt* KD (from top to bottom) | Mat67,TRiP-Mbt(HMS04518)/+ (II); Canoe-Venus (CPTI)/+ (III) |
|  | Mat67,TRiP-Mbt(HMS04518)/+ (II); alpha-catenin-Venus (CPTI)/+ (III) |
|  | Mat67,TRiP-Mbt(HMS04518)//Ubi-Ecad-GFP (II); |
|  | Ajuba-GFP/+ (X); Mat67,TRiP-Mbt(HMS04518)/+ (II) |
|  | Mat67,TRiP-Mbt(HMS04518)/Echinoid-Venus (CPTI)(II); |
|  | Sdk-Venus (CPTI)/+ (X); Mat67,TRiP-Mbt(HMS04518)/+(II) |
| Figure S8B Control | Mat67/+ (II); Canoe-Venus (CPTI)/+ (III) |
| Figure S8B *mbt* KD | Mat67,TRiP-Mbt(HMS04518)/+ (II); Canoe-Venus (CPTI)/+ (III) |
| **Supplemental Figure 9** |  |
| Figure S9A Control (from top to bottom) | Mat67/UASp-Venus-Canoe-WT (II); |
|  | Mat67/UASp-Venus-Canoe-ΔNTR (II); |
|  | Mat67/UASp-Venus-Canoe-4KA (II); |
|  | Mat67/UASp-Venus-RalgdsRBD-CanoeΔNTR-Chimera (II); |
|  | Mat67/UASp-Venus-Canoe-Δalpha-helix-1 (II); |
|  | Mat67/UASp-Venus-Canoe-Δalpha-helix-2 (II); |
|  | Mat67/UASp-Venus-Canoe-Δalpha-helix-3 (II); |
| Figure S9A *mbt* KD (from top to bottom) | Mat67,TRiP-Mbt(HMS04518)/UASp-Venus-Canoe-WT (II); |
|  | Mat67,TRiP-Mbt(HMS04518)/UASp-Venus-Canoe-ΔNTR (II); |
|  | Mat67,TRiP-Mbt(HMS04518)/UASp-Venus-Canoe-4KA (II); |
|  | Mat67,TRiP-Mbt(HMS04518)/UASp-Venus-RalgdsRBD-CanoeΔNTR-Chimera (II); |
|  | Mat67,TRiP-Mbt(HMS04518)/UASp-Venus-Canoe-Δalpha-helix-1 (II); |
|  | Mat67,TRiP-Mbt(HMS04518)/UASp-Venus-Canoe-Δalpha-helix-2 (II); |
|  | Mat67,TRiP-Mbt(HMS04518)/UASp-Venus-Canoe-Δalpha-helix-3 (II); |

**Table S2.**
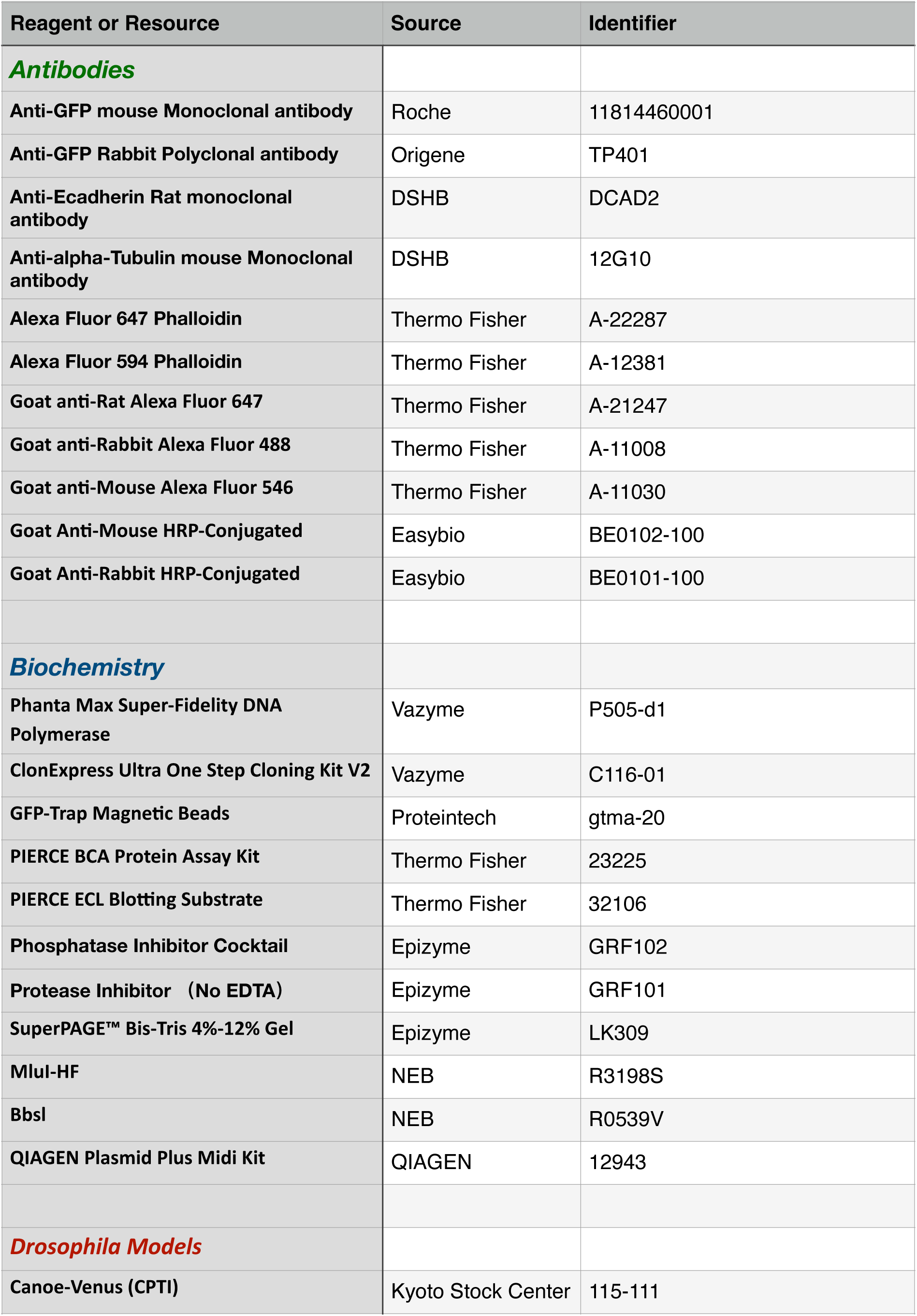
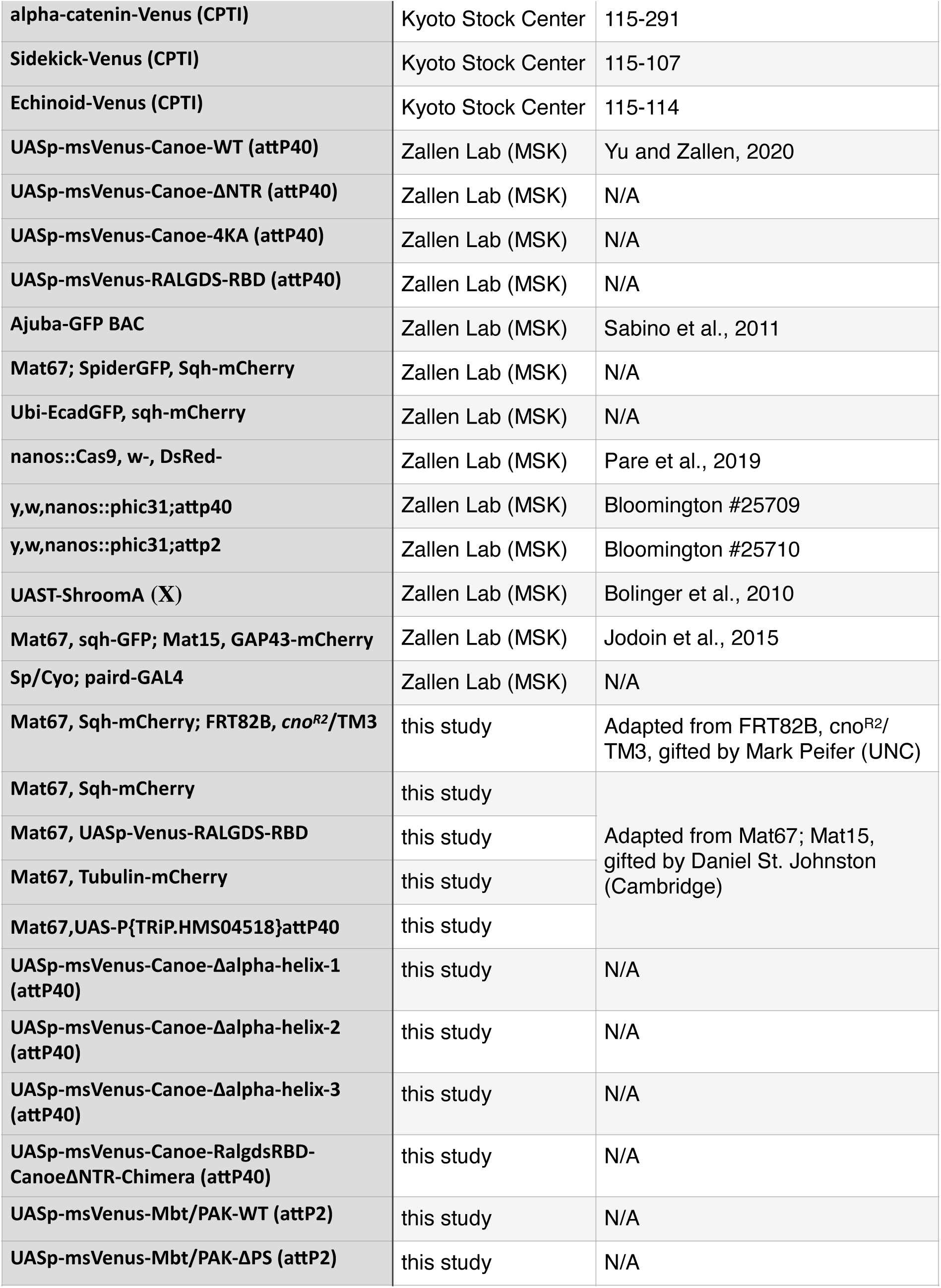
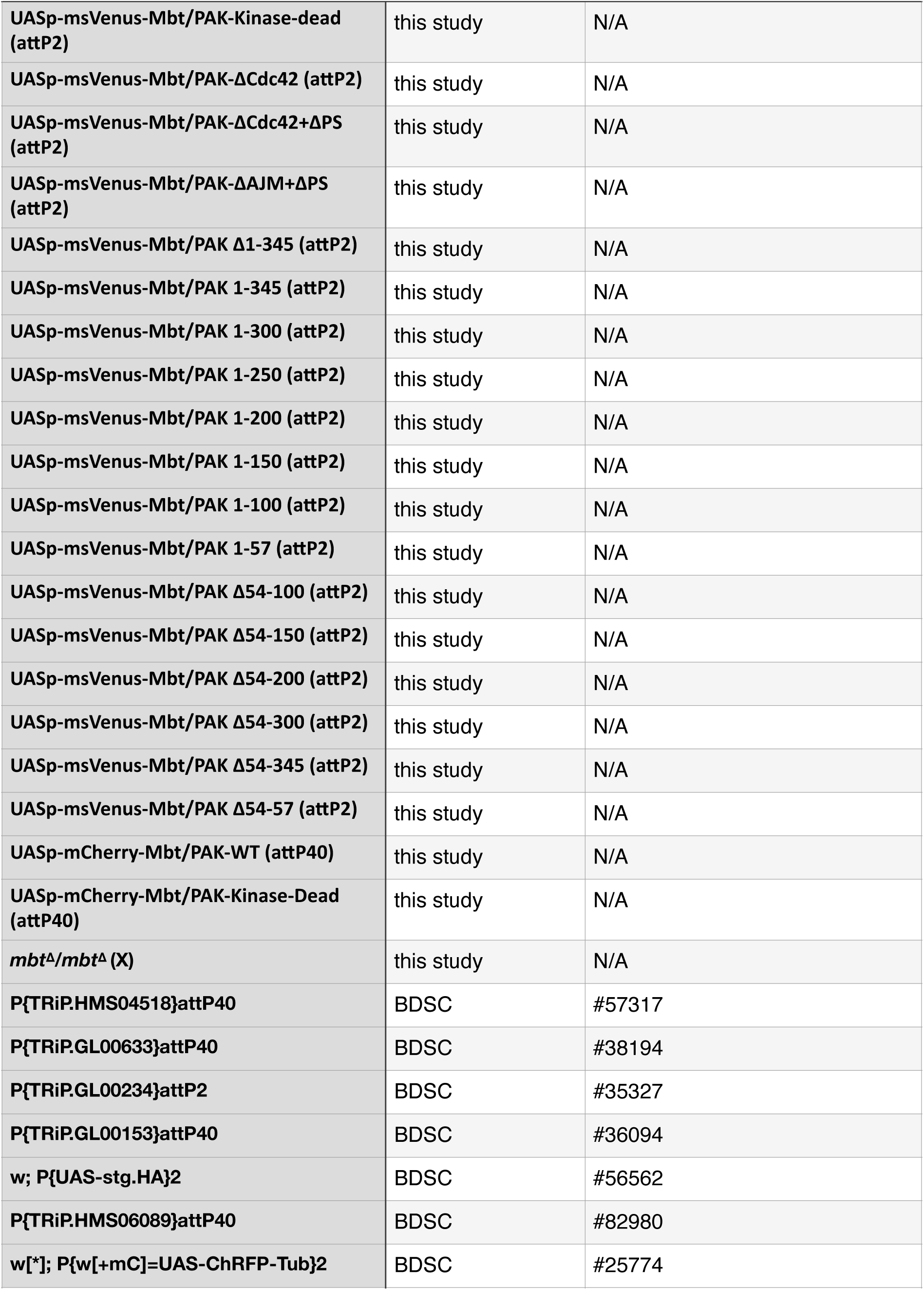
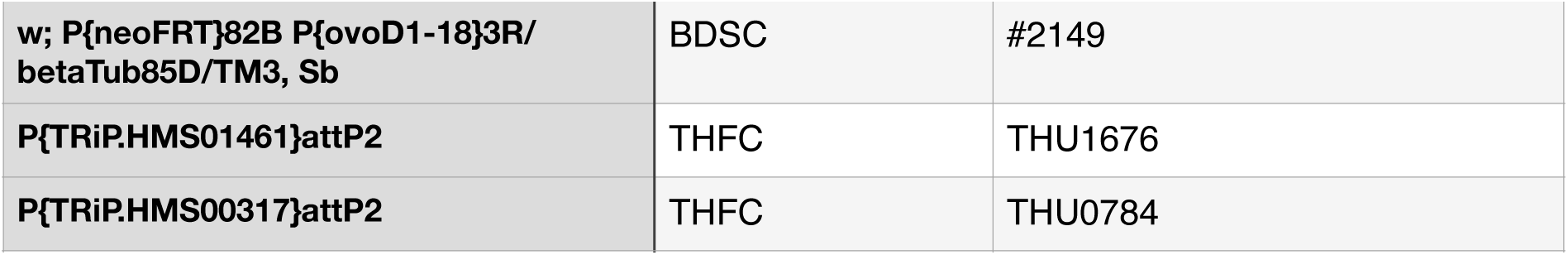

**Table S3:**
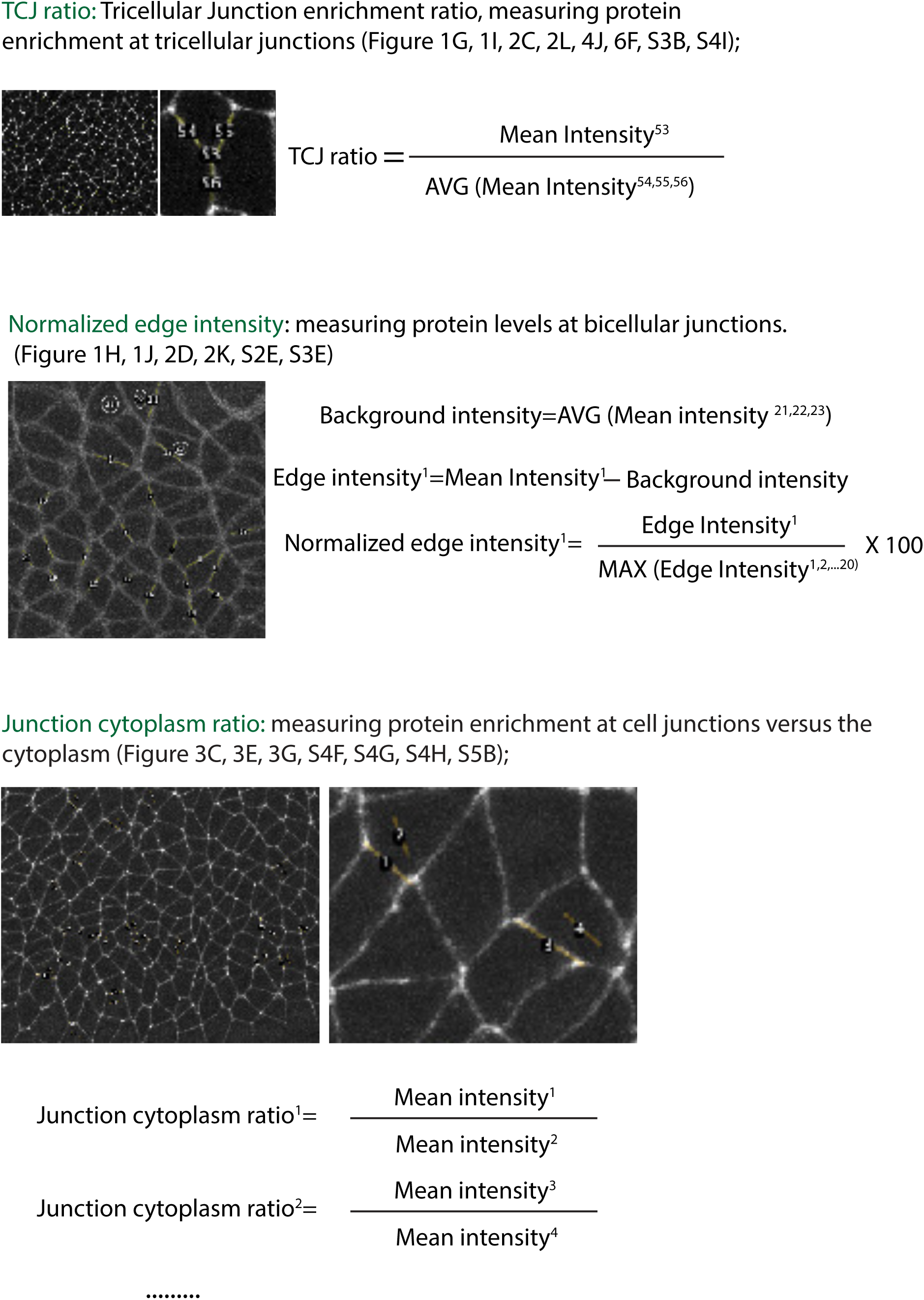
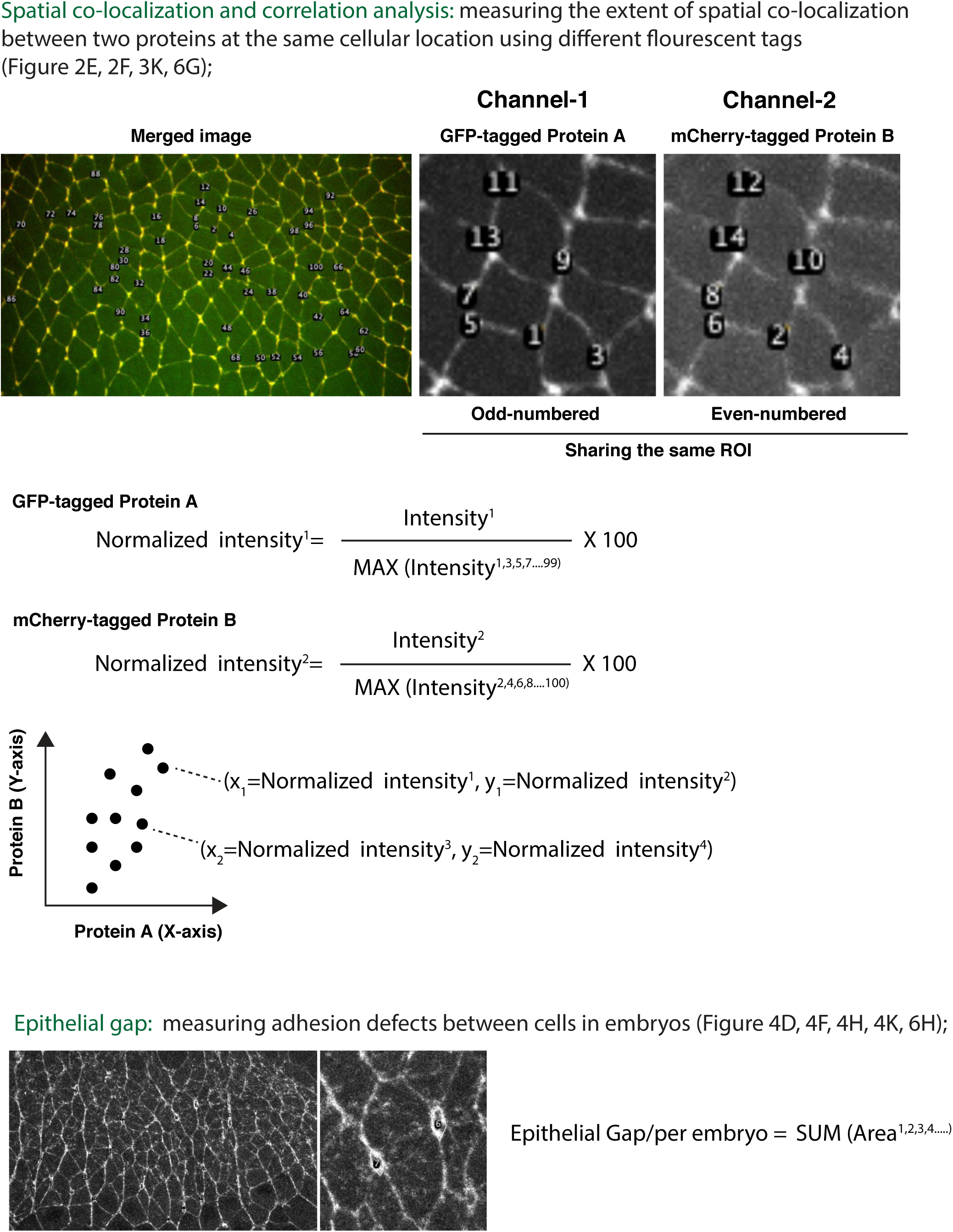
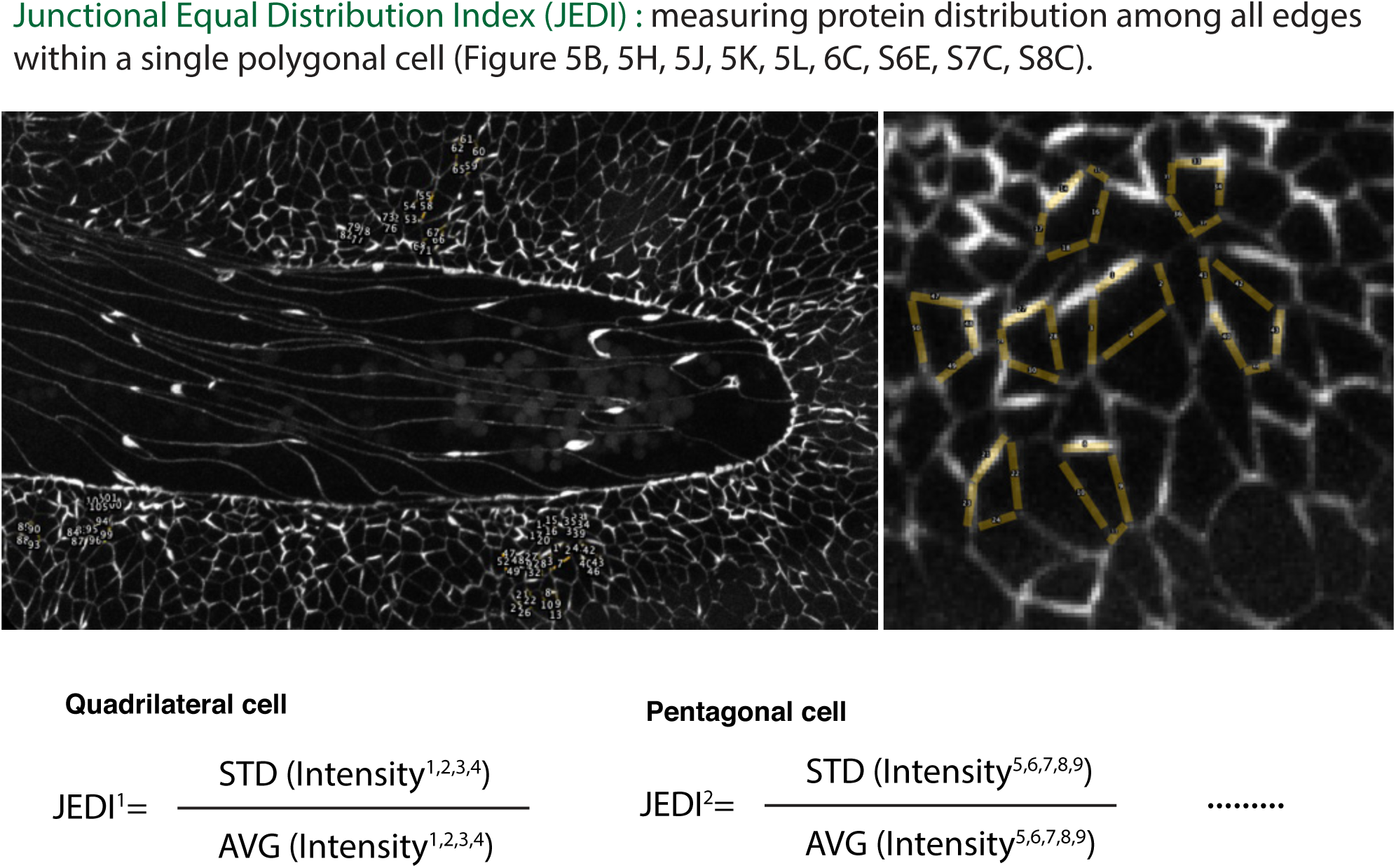
Quantification Manual

**Table S4.**
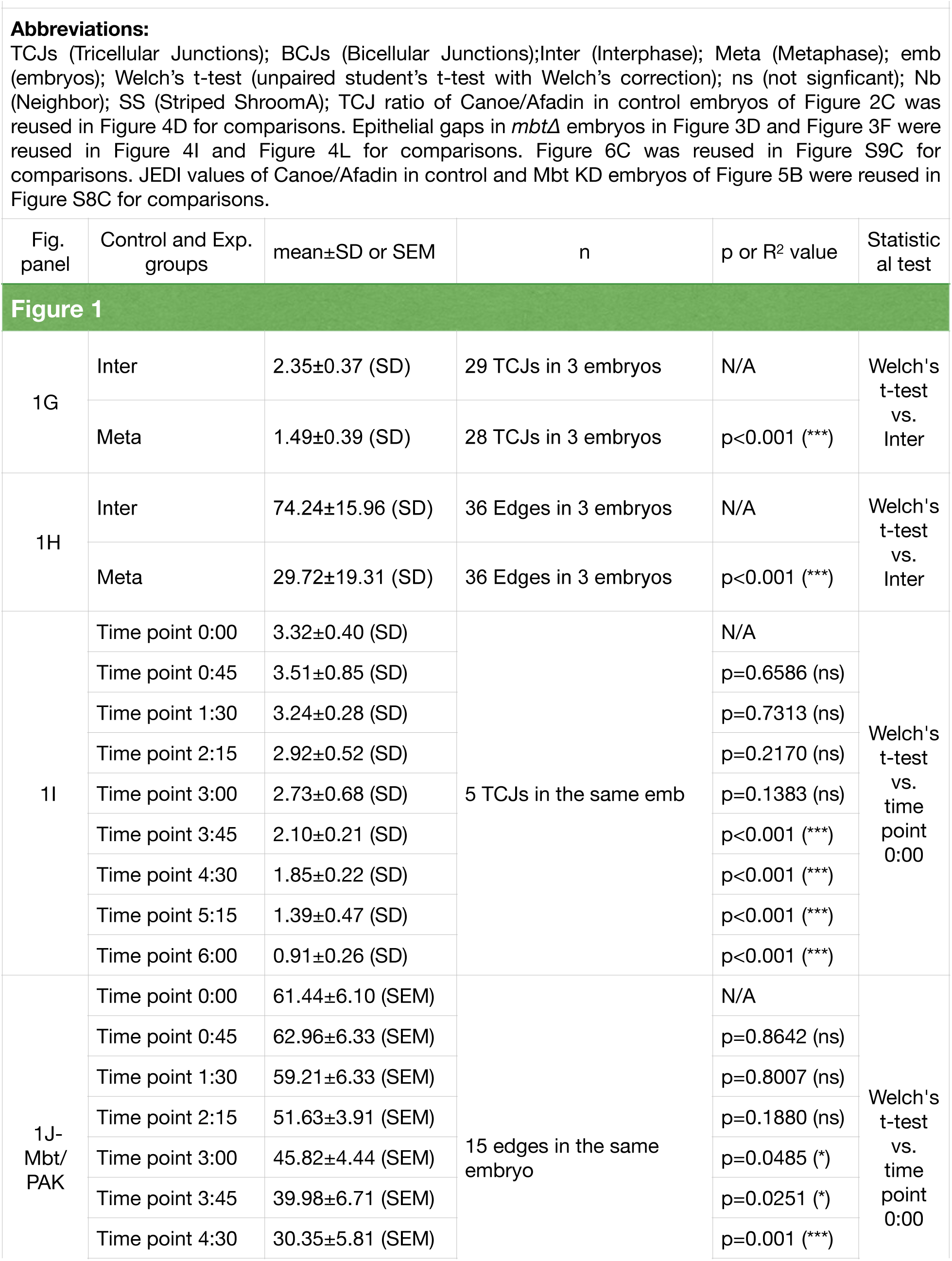
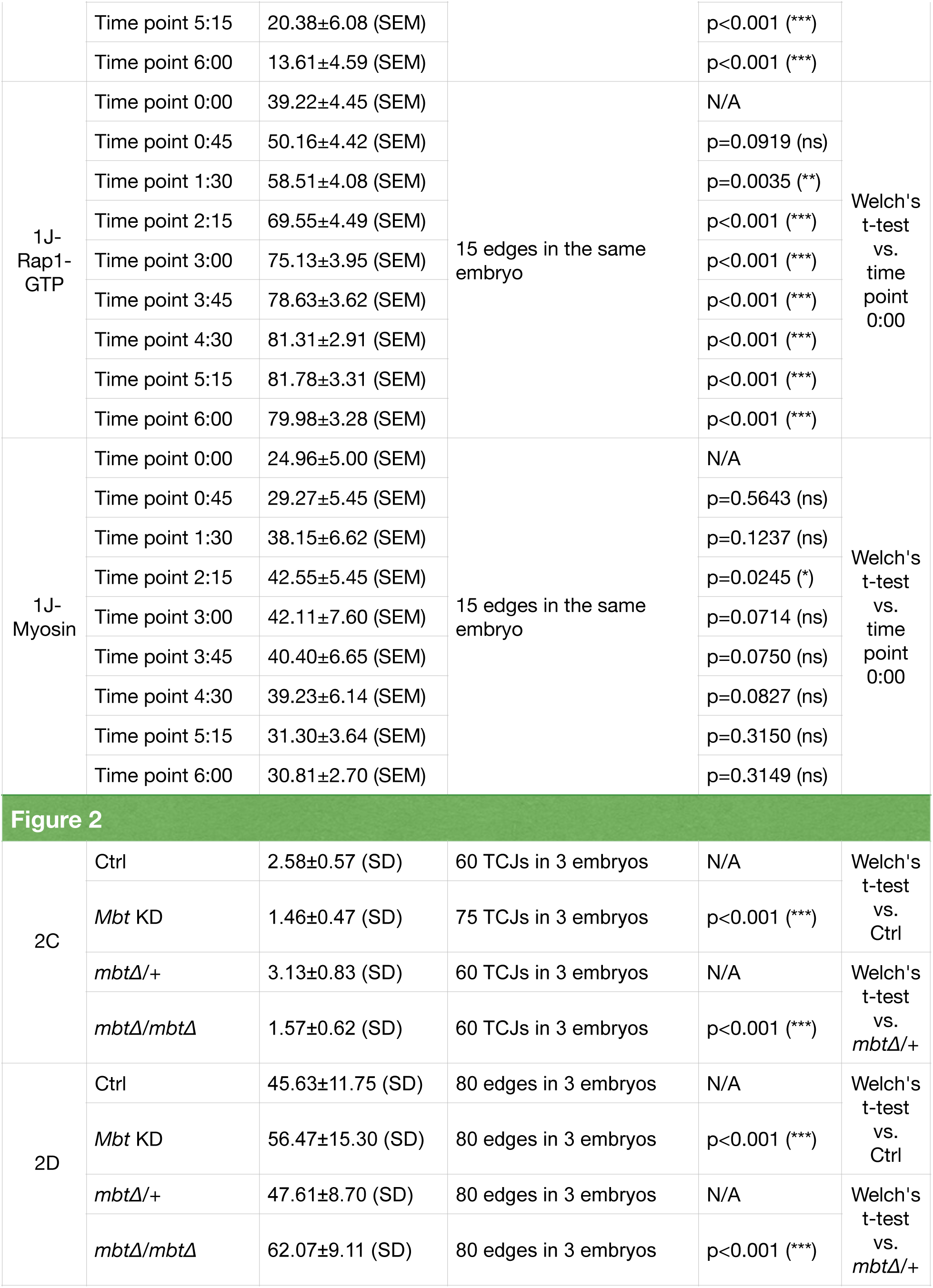
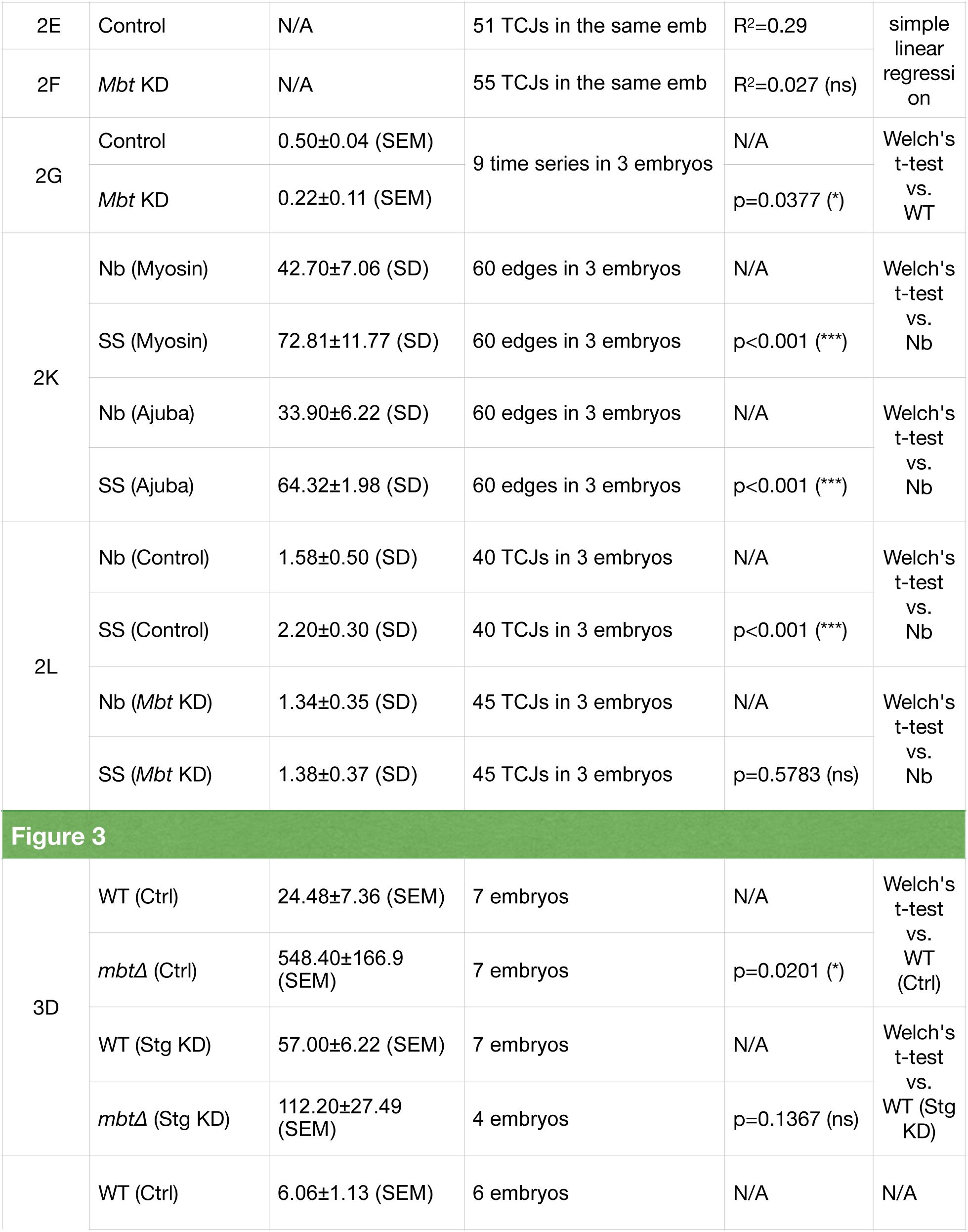
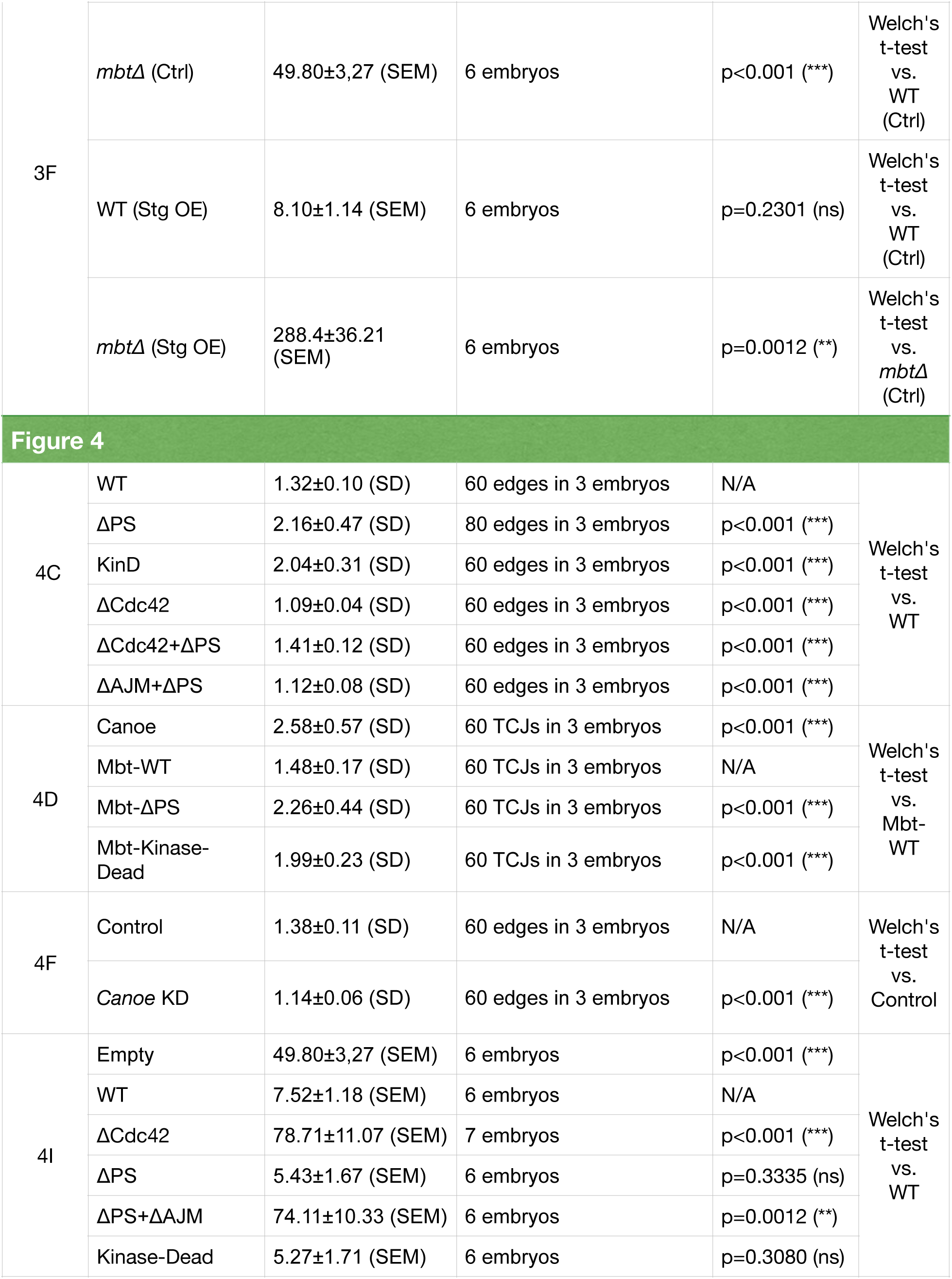
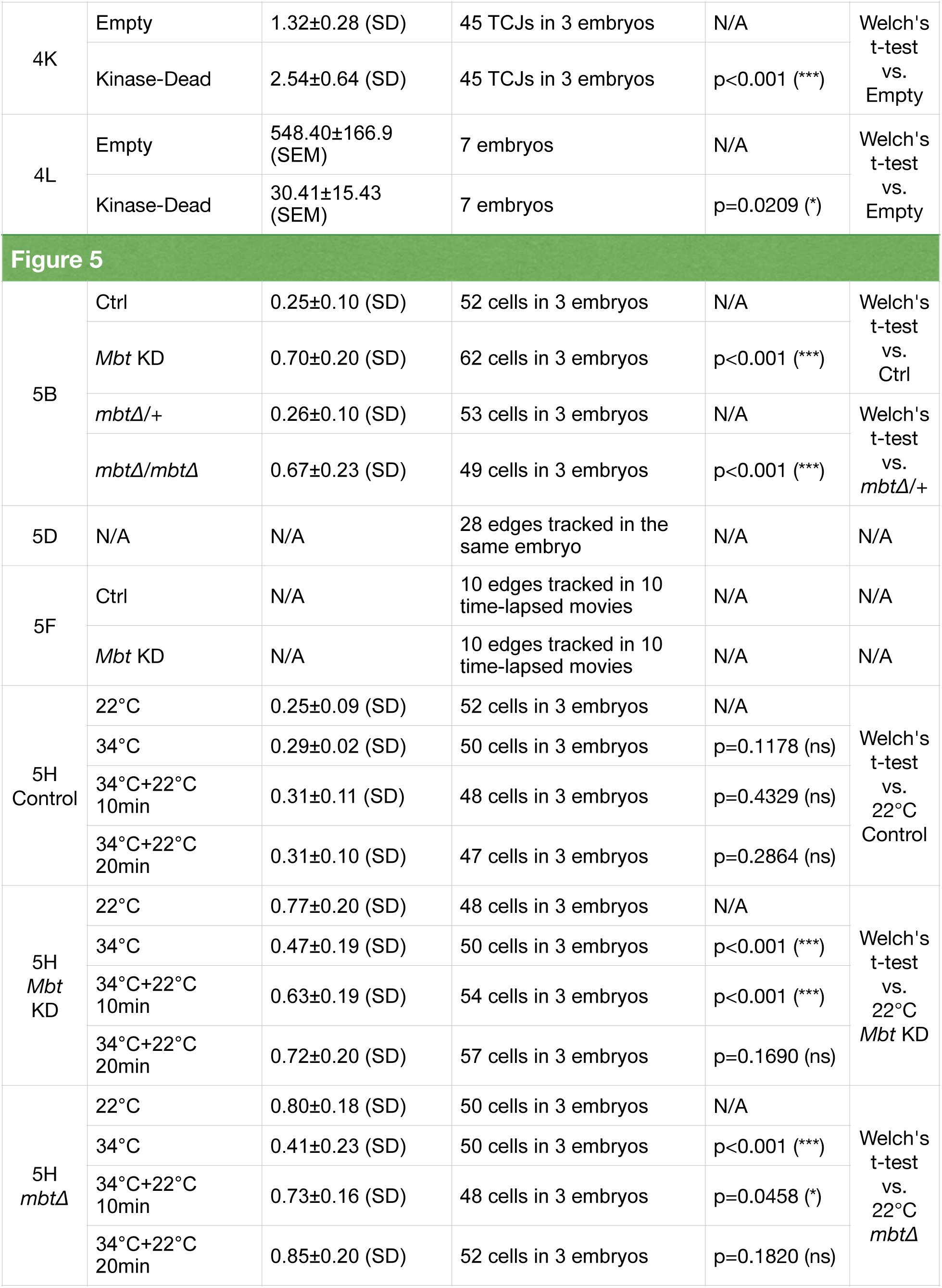
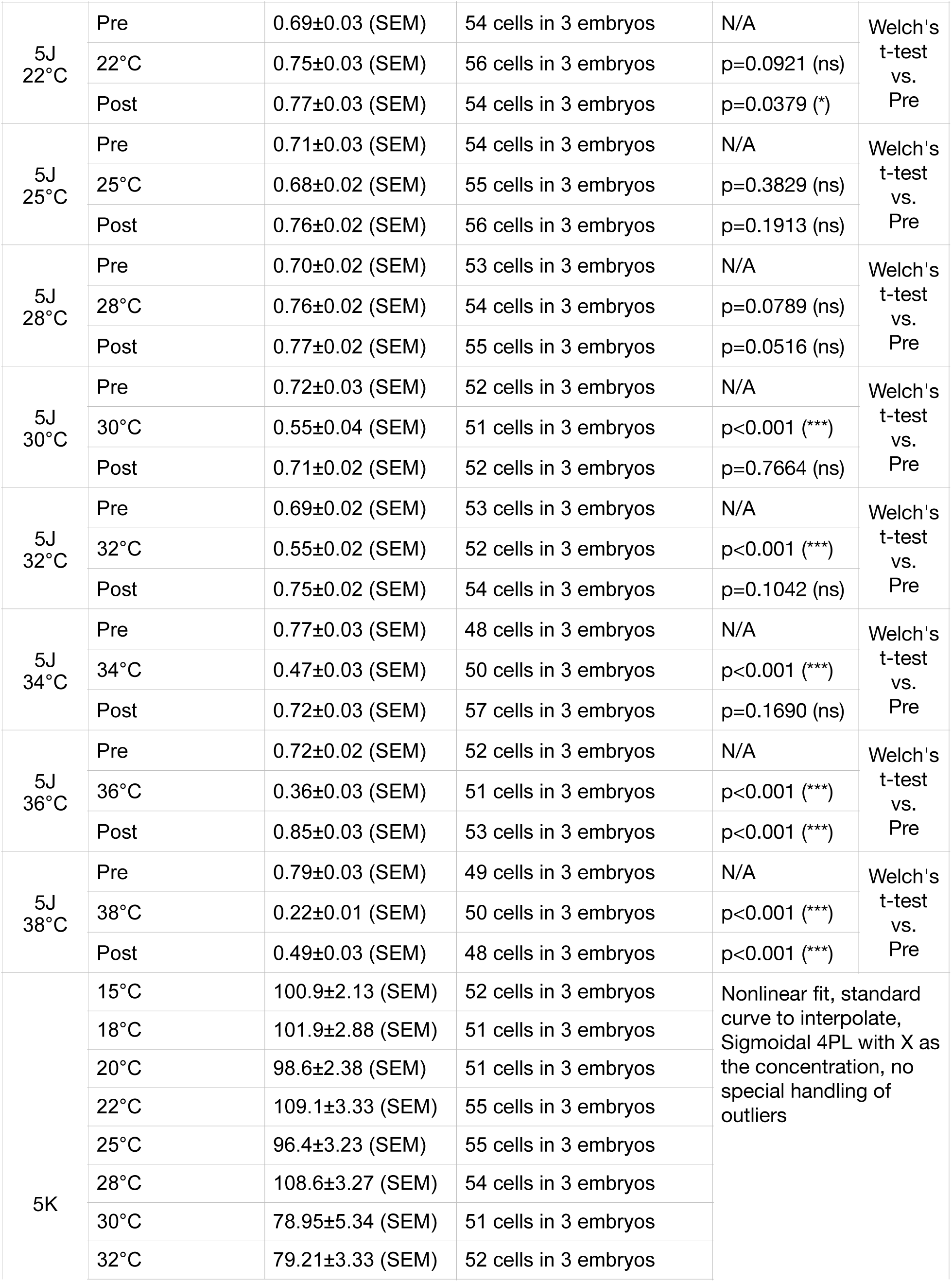
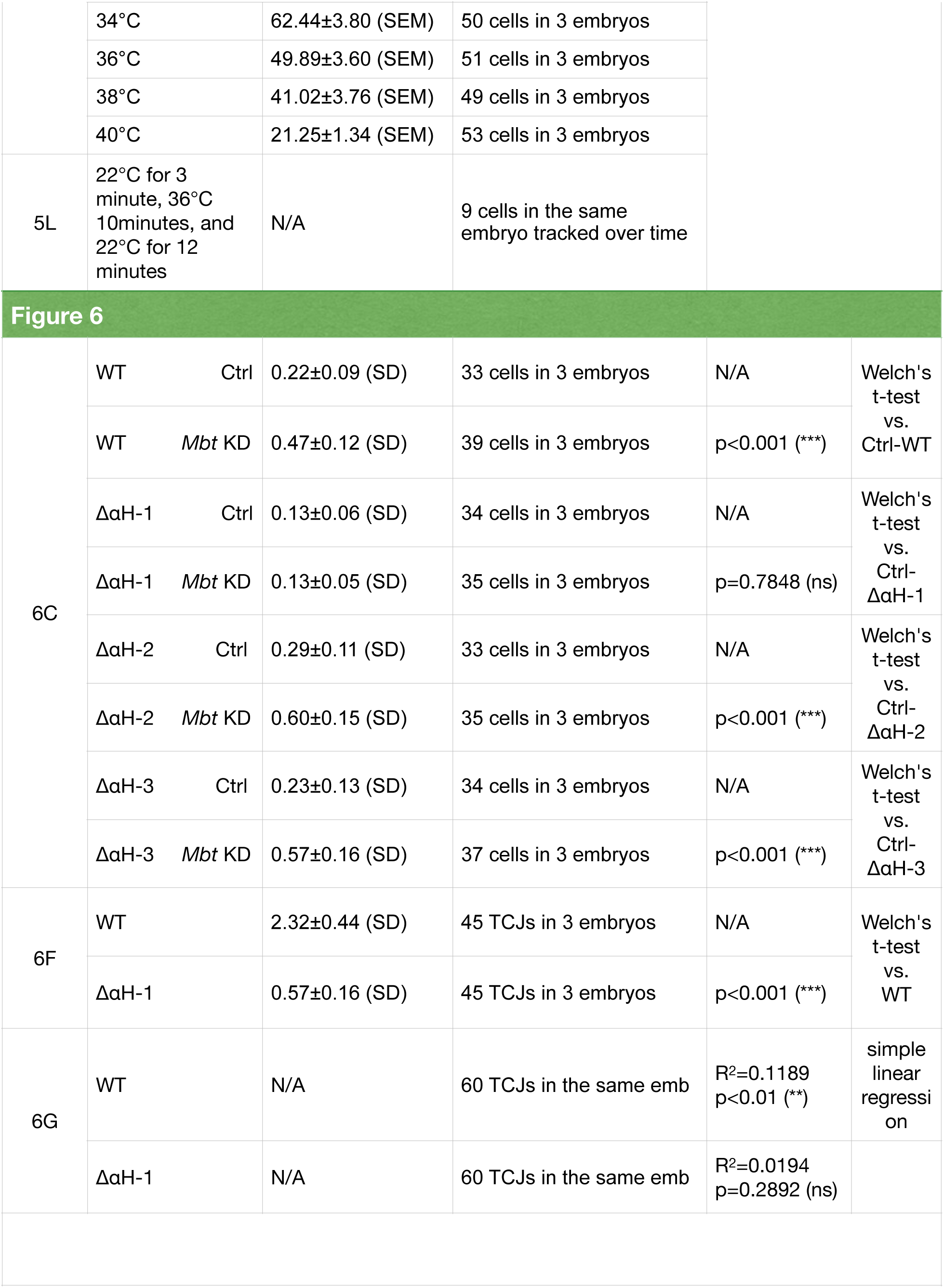
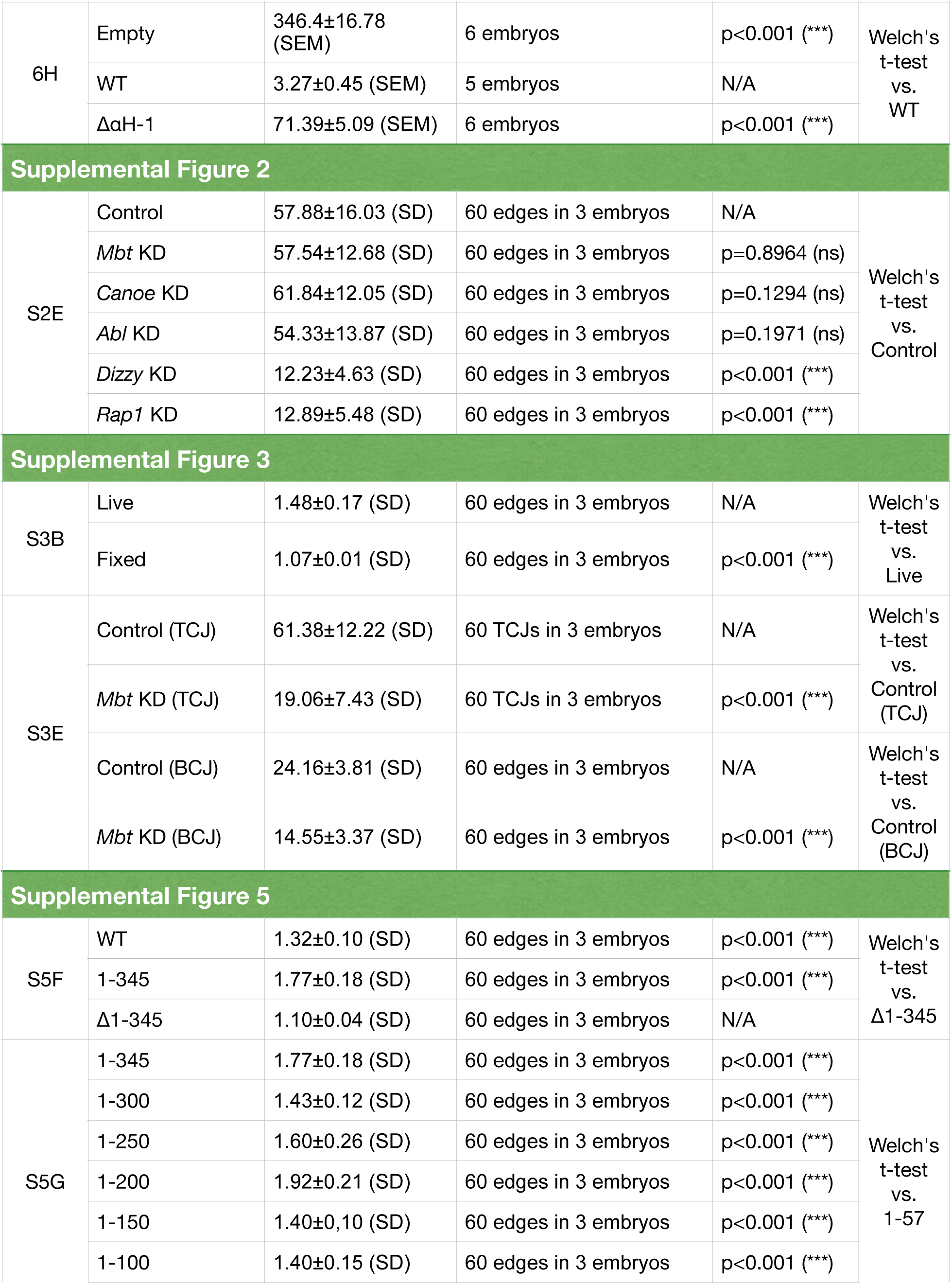
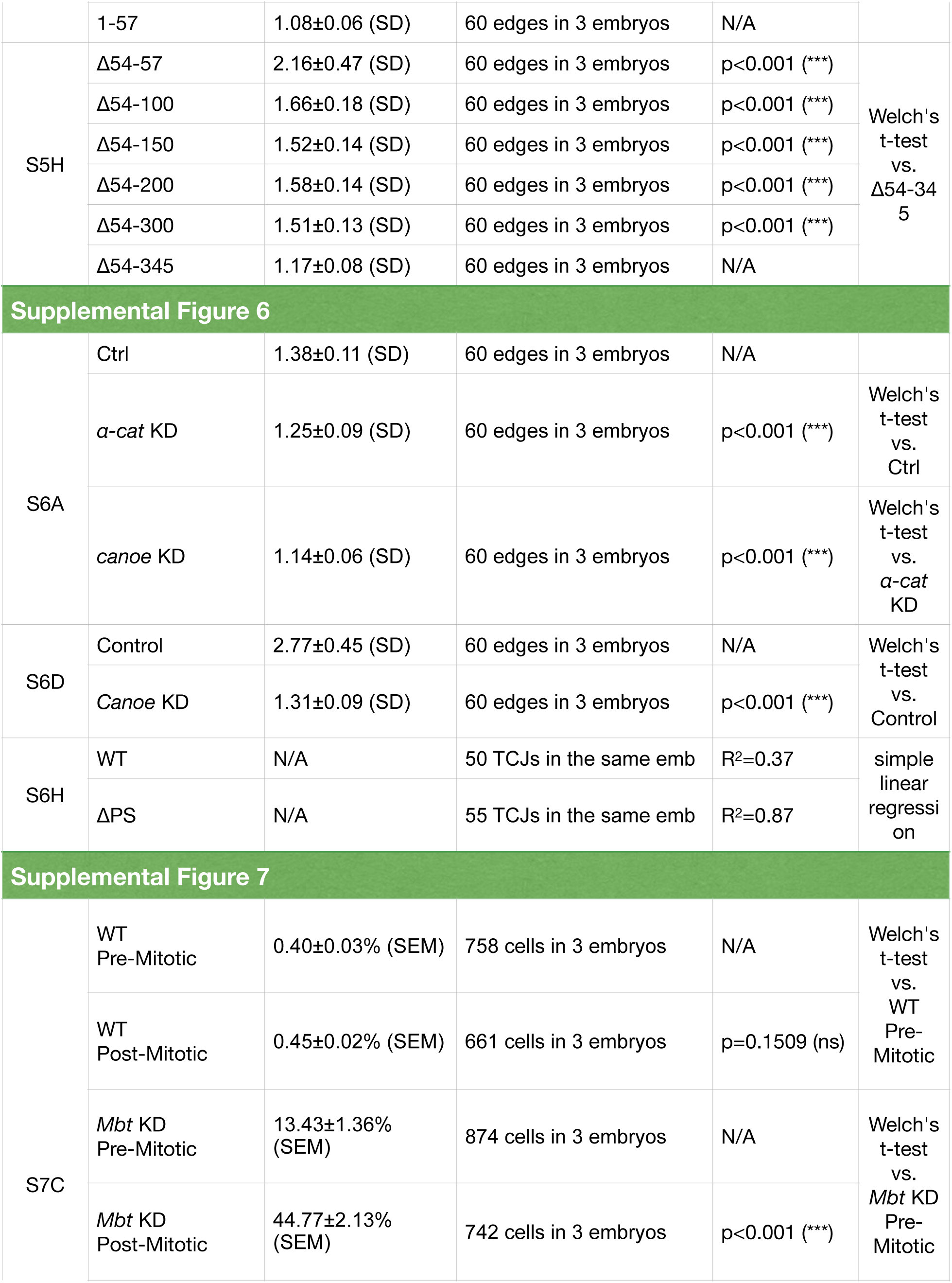
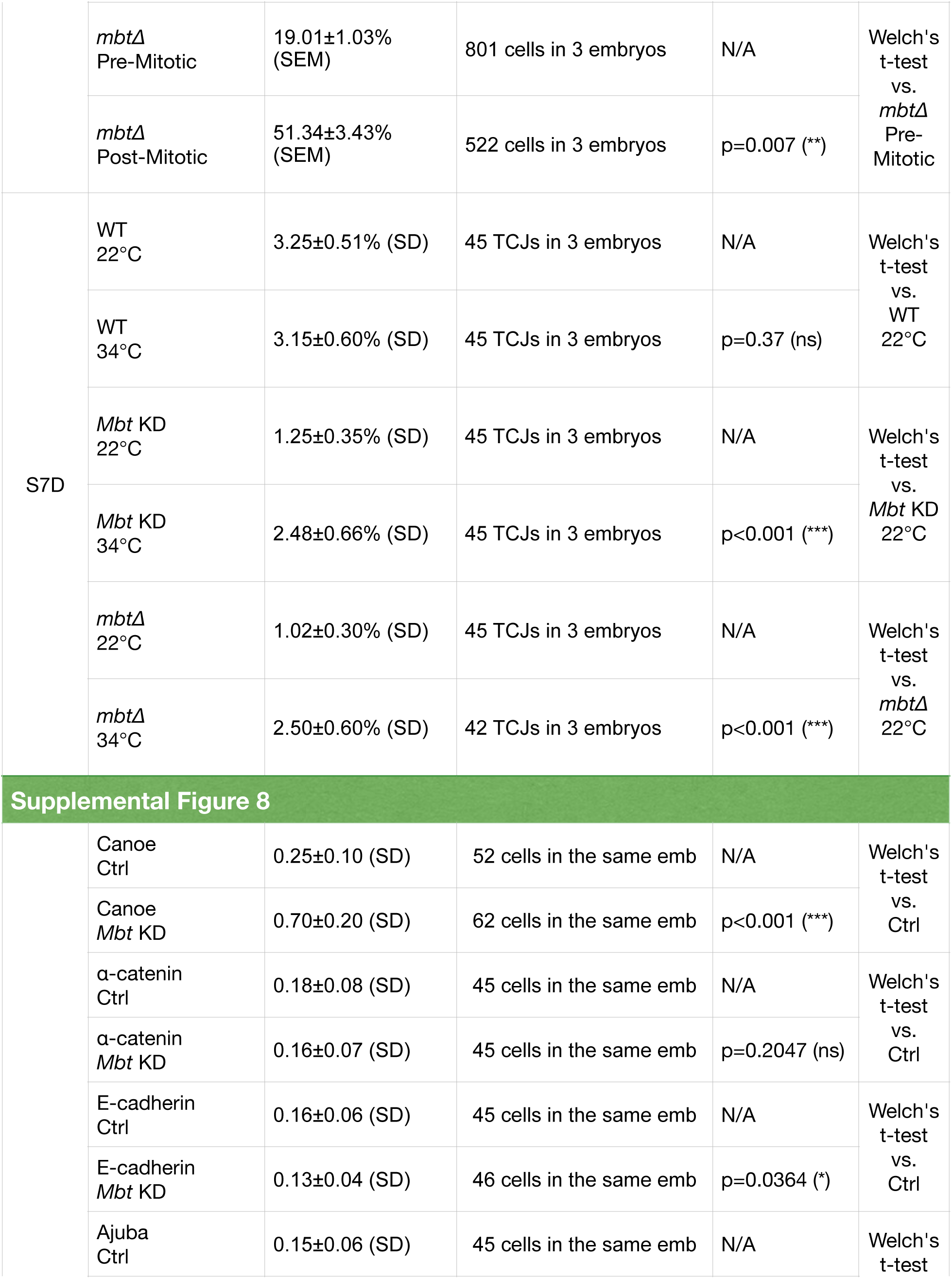
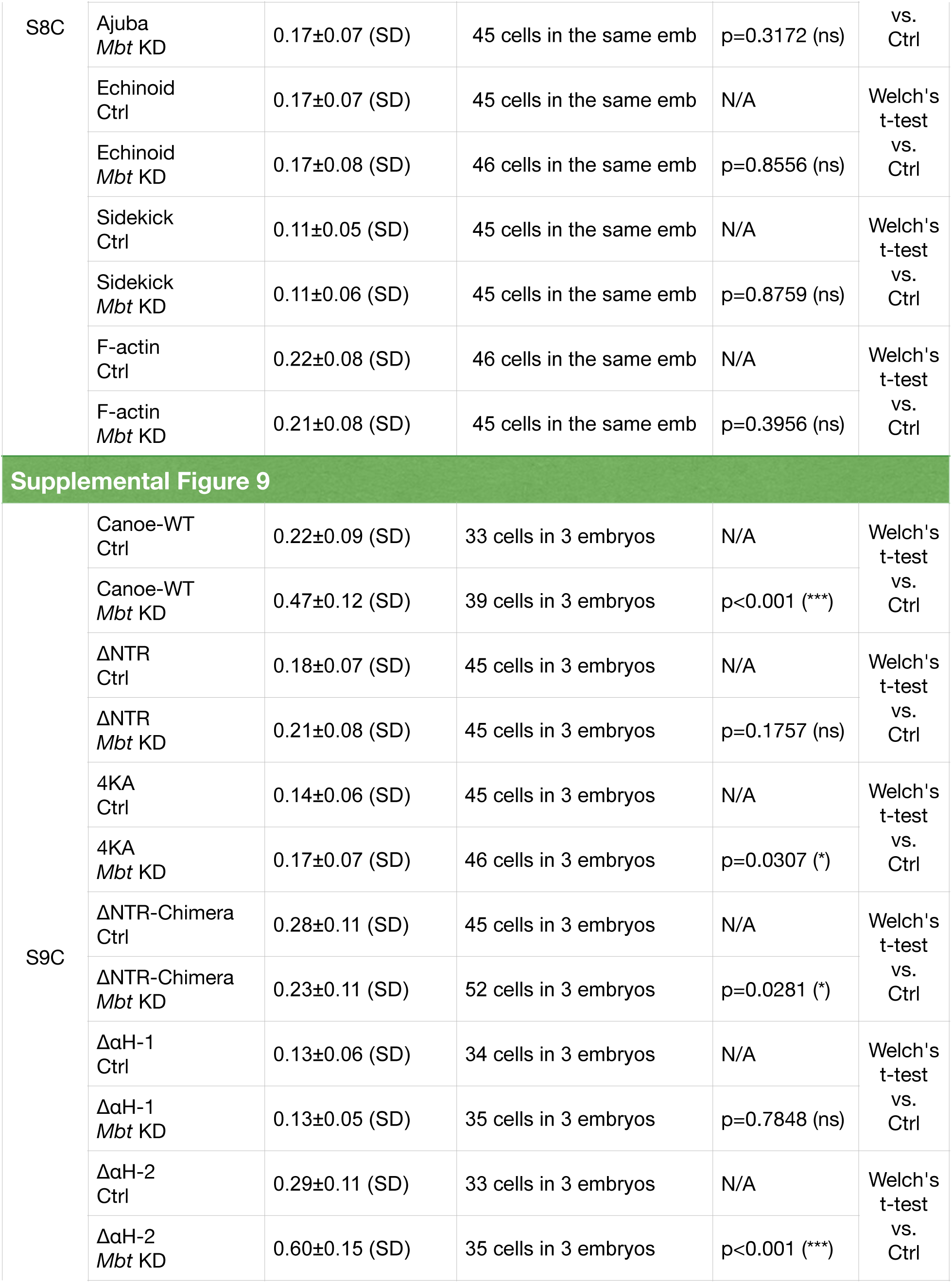
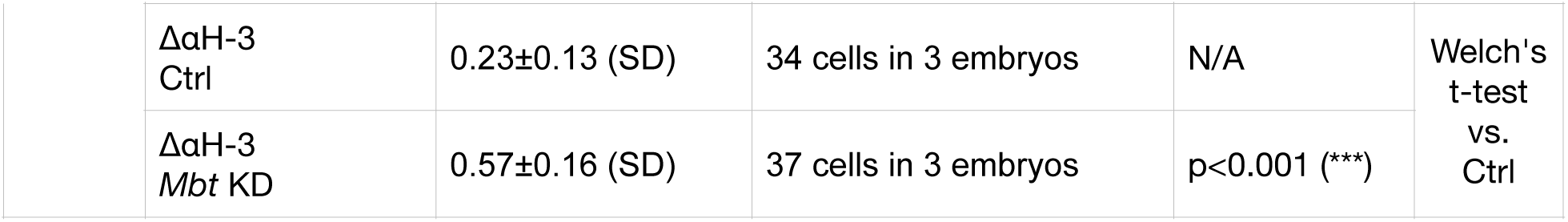
Summary of N and p values

